# Dissecting indirect genetic effects from peers in laboratory mice

**DOI:** 10.1101/302349

**Authors:** Amelie Baud, Francesco Paolo Casale, Amanda M. Barkley-Levenson, Nilgoun Farhadi, Charlotte Montillot, Binnaz Yalcin, Jerome Nicod, Abraham A. Palmer, Oliver Stegle

**Affiliations:** European Molecular Biology Laboratory, European Bioinformatics Institute, Wellcome Genome Campus, CB10 1SD Hinxton, Cambridge, UK; Department of Psychiatry, University of California San Diego, La Jolla, CA, 92093, USA; Microsoft Research New England, Cambridge, Massachusetts, USA; INSERM U1231 GAD Laboratory, 21070 Dijon, France; The Francis Crick Institute, London NW1 1AT, UK; Institute for Genomic Medicine, University of California San Diego, La Jolla, CA, 92093, USA; European Molecular Biology Laboratory, Genome Biology Unit, Heidelberg, DE

**Keywords:** Indirect genetic effects, Social genetic effects, Peer effects, Complex traits, Genotype to phenotype, Genome-wide association study

## Abstract

The phenotype of one individual can be affected not only by the individual’s own genotypes (direct genetic effects, DGE) but also by genotypes of interacting partners (indirect genetic effects, IGE). IGE have been detected using polygenic models in multiple species, including laboratory mice and humans. However, the underlying mechanisms remain largely unknown. Genome-wide association studies of IGE (igeGWAS) can point to IGE genes, but have not yet been applied to non-familial IGE arising from “peers” and affecting biomedical phenotypes. In addition, the extent to which igeGWAS will identify loci not identified by dgeGWAS remains an open question. Finally, findings from igeGWAS have not been confirmed by experimental manipulation.

We leveraged a dataset of 170 behavioural, physiological and morphological phenotypes measured in 1,812 genetically heterogeneous laboratory mice to study IGE arising between same-sex, adult, unrelated laboratory mice housed in the same cage. We developed methods for igeGWAS in this context and identified 24 significant IGE loci for 17 phenotypes (FDR < 10%). There was no overlap between IGE loci and DGE loci for the same phenotype, which was consistent with the moderate genetic correlations between DGE and IGE for the same phenotype estimated using polygenic models. Finally, we fine-mapped seven significant IGE loci to individual genes and confirmed, in an experiment with a knockout model, that *Epha4* gives rise to IGE on stress-coping strategy and wound healing.

Our results demonstrate the potential for igeGWAS to identify IGE genes and shed some light into the mechanisms of peer influence.

## Background

The phenotype of an individual can be affected not only by the individual’s own genotypes (direct genetic effects, DGE) but also by environmental factors, including the genotypes of other, interacting individuals (indirect genetic effects, IGE)(1–3) (**Figure 1a**). IGE arise when the phenotype of a focal individual is influenced by heritable traits of interacting partners (**Figure 1b**), which can include behavioural and non-behavioural traits of partners as well as modifications of the non-social environment by partners(4). IGE have been detected in many laboratory systems(5–14), livestock(15–17), crops(18), wild animals(19–21), and humans(22–27), demonstrating that they are an important component of the genotype to phenotype path and an aspect of the environment that can be studied using genetic approaches.

**Figure 1.**
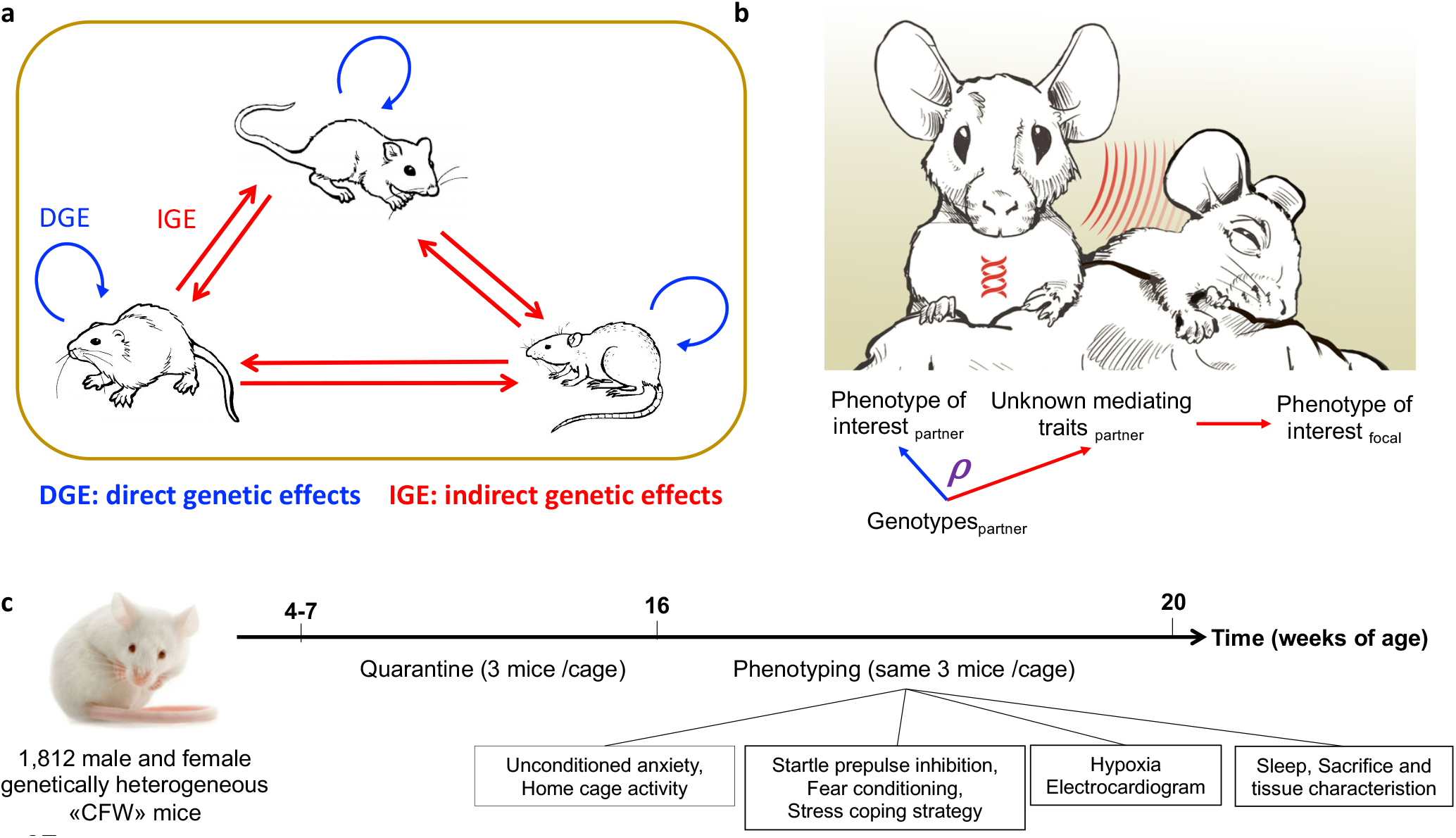
Definition of direct and indirect genetic effects and experimental design. **(a)** Direct genetic effects (DGE, blue) on an individual’s phenotype arise from the individual’s own genotypes; indirect genetic effects (IGE, red) arise from genotypes of interacting partners (cage mates). This panel illustrates a situation where all individuals are genetically heterogeneous and both DGE and IGE arise from each individual’s genotypes. **(b)** IGE on a phenotype of interest arise when two individuals interact and (unknown) heritable traits of one individual, the social partner, influence the phenotype of interest measured in the other individual. For a given phenotype of interest, the correlation ρ between DGE and IGE is equivalent to the correlation between DGE on the phenotype of interest and DGE on the traits mediating IGE on the phenotype of interest. Importantly, this correlation can be estimated even when the traits mediating IGE are not known or not measured. **(c)** Experimental design. A list of the 170 phenotypes collected on each mouse is presented in Supplementary Table 2.

Most prior studies of IGE have used polygenic modelling approaches to study aggregate genetic effects, either studying IGE mediated by <u>*specific traits of partners*</u> using trait-based models(2, 28) or polygenic risk scores(22, 25), or detecting IGE mediated by <u>*unknown heritable traits of partners*</u> using variance components models(9, 15, 29, 30). More recently, the genome-wide association study of IGE (igeGWAS) has been proposed as a strategy to identify individual genetic loci underlying IGE associations(5, 7, 8, 11, 31–35).

However, igeGWAS has only been applied in limited settings: in particular, it has not been used to study non-familial IGE from peers affecting biomedical phenotypes, despite growing evidence from polygenic models in laboratory mice(9) and in humans(25) that such effects are important. Moreover, the relationship between DGE and IGE affecting the same phenotype has not been fully addressed, such that the scope for igeGWAS to identify loci not detected by dgeGWAS is unknown. Finally, the results of igeGWAS have not yet been translated into experimentally validated genes causing IGE.

To address these issues, we leveraged a published dataset of 170 behavioural, physiological and morphological phenotypes measured in 1,812 male and female, genetically heterogeneous mice (**Figure 1c**), which we supplemented with previously unreported cage information (**Supplementary Table 1**). For each phenotype we investigated the relationship between DGE and IGE, using both polygenic analyses and GWAS. For 17 phenotypes, we fine-mapped IGE loci to identify putative causal genes underlying IGE. Finally, we validated one such gene using a knockout model.

## Results

We used the genome-wide genotypes (both LD-pruned and unpruned genotypes derived from low-coverage (0.15×), Illumina sequencing, see Methods) and 200 phenotypes for 2,073 commercially available, outbred Crl:CFW(SW)-US_P08(36) (herafter CFW) mice reported in Nicod et al. (37) and Davies et al. (38). In addition, we used previously unreported cage information provided by the authors of the original study upon request (**Supplementary Table 1**). Mice were housed in same-sex groups of three and interacted for at least nine weeks before phenotyping. We excluded any animal whose cage mates changed over the course of the experiment, as well as suspected siblings to rule out confounding from parental and litter effects. These steps resulted in a final sample size of 1,812 mice (927 females, 885 males) for analysis. We normalised each phenotype and excluded 30 phenotypes that could not be satisfactorily normalised (see Methods), yielding a total of 170 phenotypes measured in between 844 and 1,729 mice.

### Polygenic analysis of the correlation between DGE and IGE

Initially, we used polygenic models to assess the extent to which loci are shared between DGE and IGE affecting the same phenotype. Briefly, for each trait, we estimated the genetic correlation *ρ* between DGE and IGE. As this correlation is equivalent to the correlation between DGE on the phenotype of interest and DGE on the traits of partners mediating IGE (**Figure 1b**), a correlation coefficient of 0 would indicate that the traits mediating IGE are genetically uncorrelated (in the classical sense) to the phenotype of interest, whereas a correlation coefficient of ±1 would indicate that the phenotype of interest itself mediates IGE. For 28 traits with evidence for marginal DGE and IGE (>5% variance explained; **Supplementary Figure 1**), we performed hypothesis tests for both models (**Figure 2** and **Supplementary Table 2**). We found that *ρ* was different from zero for ten out of twenty eight phenotypes (P < 0.05), indicating that, often, the traits mediating IGE on a phenotype of interest are genetically correlated (in the classical sense) with the phenotype of interest. Evidence that *ρ* was different from zero was strongest for mean weight of the adrenal glands, which correlates with stress(39), mean platelet volume, LDL cholesterol levels, and rate of healing from an ear punch. Second, *ρ* was different from plus or minus one for ten phenotypes (P < 0.05), with the strongest evidence for a measure of stress-coping strategy (immobility in the forced swim test) and rate of healing from an ear punch. These results indicate that IGE on a phenotype of interest are often mediated by traits of partners other than the phenotype of interest. To uncover those traits, we turned to igeGWAS.

**Figure 2.**
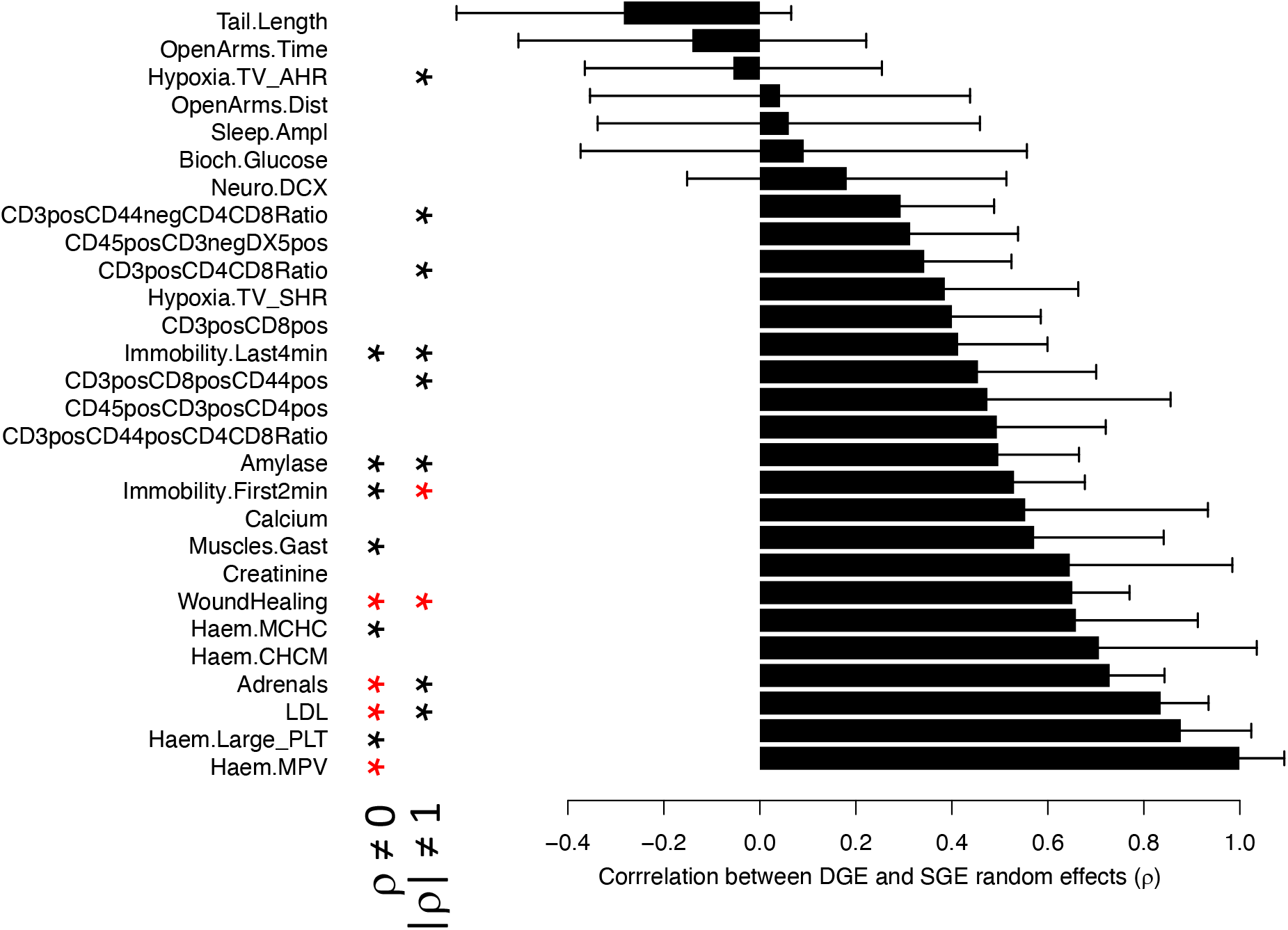
Correlation coefficients ρ between DGE and IGE estimated using polygenic models. ρ is shown for 28 phenotypes with marginal DGE and IGE greater than 5%. Error bars denote standard errors. Asterisks on the left show phenotypes for which ρ is significantly different from 0 (black: P < 0.05, red: Bonferroni-corrected P < 0.05). Asterisks on the right show phenotypes for which |ρ| is significantly different from 1 (black: P < 0.05, red: Bonferroni-corrected P < 0.05). Numerical values are provided in Supplementary Table 2.

### igeGWAS and dgeGWAS of 170 phenotypes

Next, to compare DGE and IGE at the level of individual loci, we considered the LD-pruned set of variants and performed igeGWAS and dgeGWAS in an analogous manner for each one of the 170 phenotypes. For igeGWAS, we estimated the “social genotype” of a mouse at a variant as the sum of the reference allele dosages across its two cage mates at the variant(31, 40), and tested for association between this social genotype and the phenotype of interest. To avoid spurious associations, we accounted for background IGE, background DGE and shared environmental (cage) effects using random effect components in a linear mixed model (Methods). Additionally, we included a fixed effect covariate for DGE arising from the tested variant in igeGWAS. This approach accounts for correlations between direct and social genotypes that arise when each individual serves as both focal individual and social partner in the analysis, a strategy that maximises sample size when all the individuals are genotyped and phenotyped(31, 40). Accounting for such correlations was required to obtain appropriately calibrated P values in our cohort (**Supplementary Figure 2**), and theoretical considerations show that it is required even when considering strictly unrelated samples (**Supplementary Note**). Finally, we adapted previous strategies(37, 41, 42) based on genome-wide permutations to control the per-phenotype FDR (see Methods), thereby accounting for the specific patterns of linkage disequilibrium present in the sample.

igeGWAS identified a total of 24 significant loci across 17 of the 170 tested phenotypes (FDR < 10%), including measures relevant to behavior, adult neurogenesis, blood biochemistry, red and white blood cells, apparent bone mineral content, electrocardiography, and ventilatory responses to acute hypoxia (**Supplementary Table 3**). The 17 phenotypes with one or more IGE loci tended to have a higher aggregate contribution of IGE (across the genome) than phenotypes without significant IGE loci (averages of 3.8% and 2.8% respectively), a trend that was not significant (one-sided t-test P = 0.14).

To enable a direct comparison between igeGWAS and dgeGWAS, we performed dgeGWAS for each phenotype using the same approach as taken for igeGWAS, including random effects for DGE and IGE polygenic effects and cage effects and including a fixed effect covariate for IGE arising from the tested variant. This identified 120 significant DGE loci for 63 phenotypes (FDR<10% **Supplementary Table 4**). Consistent with the difference in the number of discoveries, we observed that significant IGE loci had, on average, lower effect sizes (proportion of phenotypic variance explained) than DGE loci (**Supplementary Figure 3**). In light of the observed effect sizes and due to the winner’s curse (or Beavis effect (43, 44)), we expect a larger proportion of significant IGE loci to be false associations, compared to significant DGE loci.

There was no overlap between significant DGE and IGE loci for the same phenotype, or even for related phenotypes (**Figure 3**). This observation was expected based on the moderate values observed for the correlation *ρ* between DGE and IGE and the limited power of dgeGWAS and igeGWAS. However, we identified further reason why dgeGWAS and igeGWAS might identify different loci: using simulations to identify key parameters determining the power of igeGWAS, we found that both the number of cage mates and the mode of aggregation across cage mates (i.e. whether the IGE received by a focal mouse correspond to the sum or the average of the IGE emitted by its cage mates) are important, in addition to the parameters also determining the power of dgeGWAS, namely minor allele frequency (MAF) and allelic effect (**Supplementary Figure 4**). Thus, for a given MAF, allelic effect, and a number of cage mates equal to two as is the case in this study, dgeGWAS is expected to have greater power than igeGWAS if IGE get averaged across the two cage mates, but igeGWAS is expected to have greater power than dgeGWAS if IGE sum up across the two cage mates. As sample sizes increase for dgeGWAS and igeGWAS, the moderate genetic correlation *ρ* between DGE and IGE and the differences in power between dgeGWAS and igeGWAS dictated by the number of cage mates and the mode of aggregation across cage mates (sum or average) will continue to drive the identification of different loci by dgeGWAS and igeGWAS.

**Figure 3.**
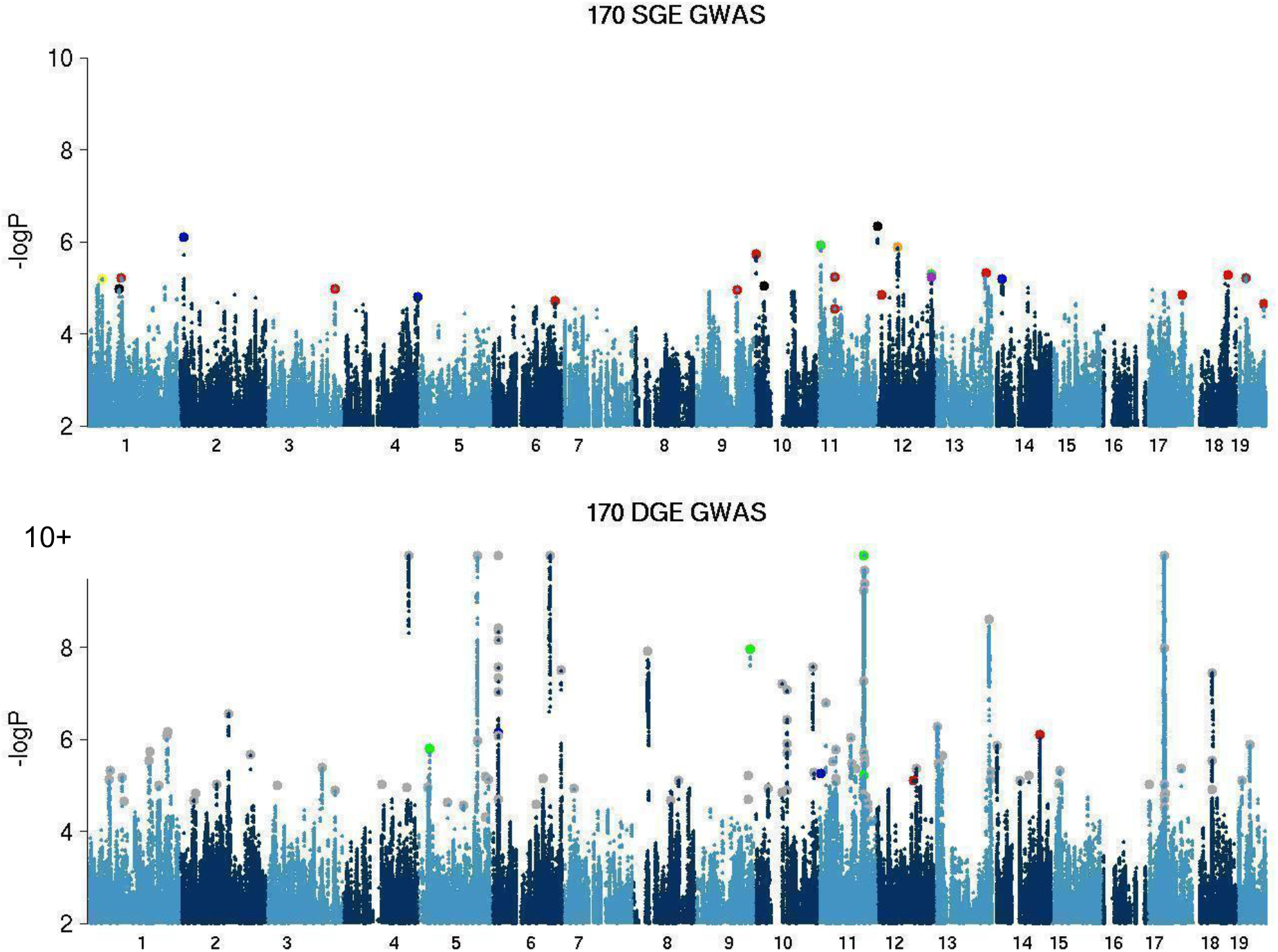
Superimposed Manhattan plots corresponding to igeGWAS (top panel) and dgeGWAS (bottom panel) of the same 170 phenotypes. DGE associations with a negative log P value greater than 10 were truncated at this threshold (as indicated by 10+); also, data points with negative log P values smaller than 2 are not shown. The larger dots correspond to the most significant variant at each significant IGE or DGE locus (FDR < 10%). In the IGE panel (top), each colour corresponds to a class of phenotypes: behavioural (red, includes 7 behavioural phenotypes with a significant IGE locus), adult neurogenesis (black, 2 phenotypes with a significant IGE locus), immune (orange, 1 phenotype with a significant IGE locus), haematological (yellow, 1 phenotype with a significant IGE locus), blood biochemistry (blue, 2 phenotypes with a significant IGE locus), bone phenotypes (green, 2 phenotypes with a significant IGE locus), heart function (brown, 1 phenotype with a significant IGE locus), and lung function (purple, 1 phenotype with a significant IGE locus). In the DGE panel (bottom), the same colouring scheme is used as in the IGE panel except for grey dots, which are for phenotypes that do not have any significant IGE locus.

### Identification of putative causal genes for experimental evaluation

Linkage disequilibrium decays faster in the CFW population than in many other mouse populations used for mapping, which facilitates identification of putative causal genes at associated loci(36, 37, 45). To identify such genes, we fine-mapped the 24 significant IGE loci using the full set of variants (rather than the pruned set used for igeGWAS) in the 1.5Mb window surrounding the most significant variant at the locus, which corresponds, in this sample, to the average 95% confidence interval for the association(37). We then identified, for each significant IGE locus, all of the genes that either overlapped the associated plateau or were located in direct proximity (see Methods, genes listed in **Supplementary Table 3** and local association plots in **locusZooms_SupplTable3.zip**). At seven loci there was a single putative causal gene: *Abca12* at a locus for adult neurogenesis, *Epha4* (stress-coping strategy), *Pkn2*, *Slit3* and *Pgk1-rs7* (at three different loci for sleep), *H60c* (home cage activity), and *Adcy1* (osteopetrosis).

One example of a putative causal IGE gene identified via this strategy is *Epha4*, which was identified at an IGE locus on chromosome 1 for immobility during the first two minutes of the forced swim test (FST), a measure of stress-coping strategy(46) (**Figure 4a** and **Supplementary Figure 5**). We focused on *Epha4* initially because it was the only putative causal gene at a significant locus, the locus was in the top half of the list in terms of significance, and a knockout mouse model was readily available from a neighbouring institute.

**Figure 4.**
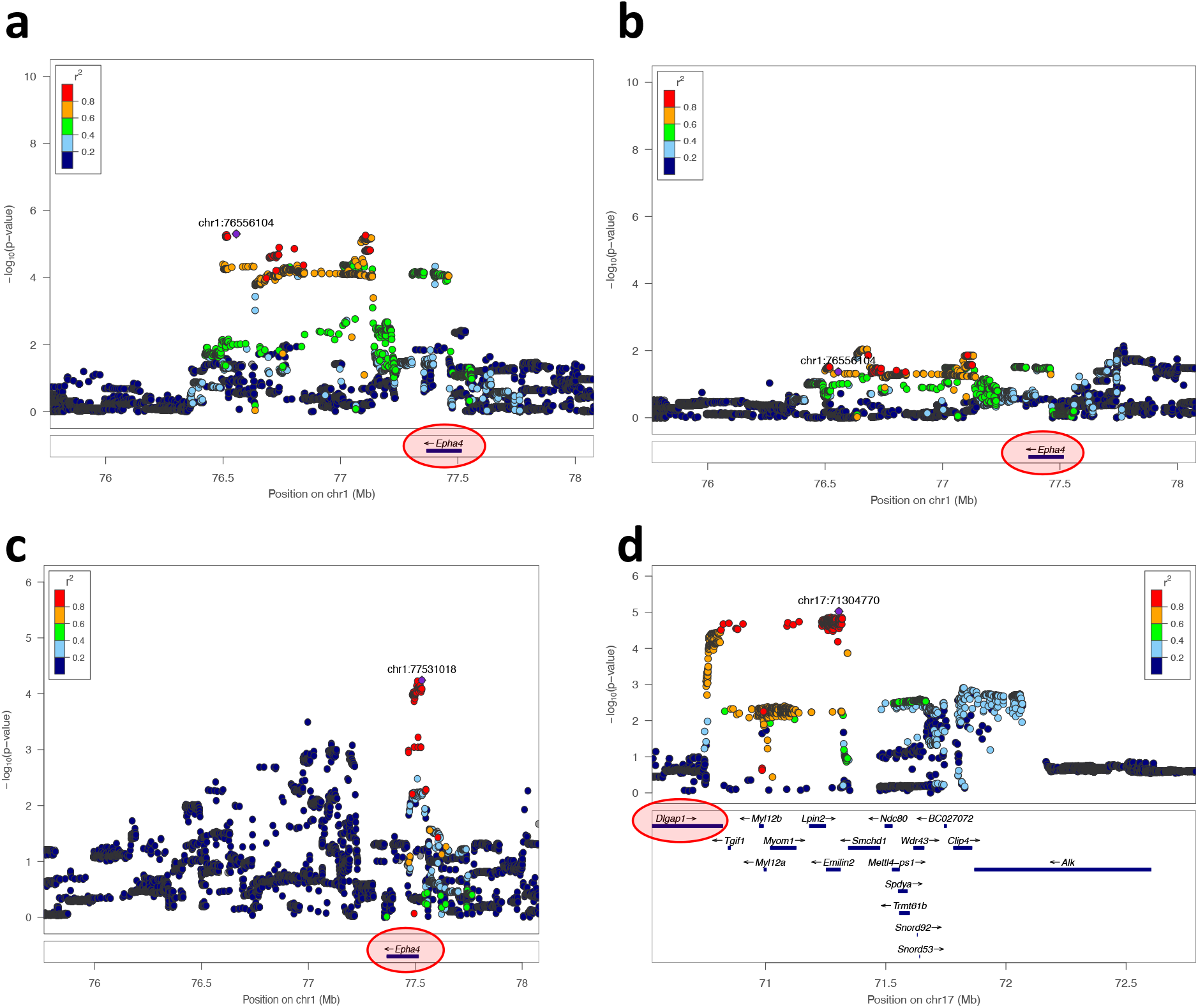
Locus zoom plots for the four associations in CFW outbred mice that were subsequently tested in an experiment with Epha4 and Dlgap1 knockout models. **(a)** Significant IGE locus on chromosome 1 for immobility during the first two minutes of the forced swim test (FST), a measure of stress-coping strategy. Epha4 was identified as the only putative causal gene at this locus (see Methods). **(b)** Same locus and phenotype but DGE, rather than IGE, are shown. The plot shows little evidence of DGE on FST immobility at the Epha4 locus. **(c)** Suggestive IGE association at the Epha4 locus with rate of healing from an ear punch, a phenotype of particular interest (-logP = 4.1, FDR > 10%). **(d)** Second significant (FDR < 10%) IGE locus for immobility in the FST (this time measured during the last four minutes of the test). Dlgap1 is one of eight putative causal genes at this locus. It was singled out because of its functional similarity and co-expression with Epha4 (see Text).

*Epha4* encodes a synaptic protein that plays an important role in synaptic plasticity in the hippocampus(47, 48) and DGE of *Epha4* on FST immobility have been reported(49, 50). Therefore we evaluated the possibility that *Epha4* directly influences stress-coping strategy and that the stress-coping strategy of a mouse in the weeks prior to or during the FST gets copied by the other mice in the cage (behavioural contagion), thereby giving rise to IGE on stress-coping strategy. To investigate this hypothesis, we tested whether *Epha4* had direct effects on FST immobility in CFW mice, using the full set of variants in the same 1.5Mb window including *Epha4* as for IGE analysis. We found little evidence that *Epha4* directly affects FST immobility in CFW mice (maximum -logP value at the locus: 2.14, **Figure 4b**), making it unlikely that behavioural contagion explains the detected IGE in CFW mice.

In addition to the significant IGE association between *Epha4* and FST immobility, we found suggestive evidence for an IGE association between *Epha4* and rate of healing from an ear punch (igeGWAS -logP value = 4.1, FDR > 10%, **Figure 4c**). This finding was of particular interest because the *Epha4* locus was among the three most significant IGE loci for wound healing (all three loci with -logP=4.1) and because IGE on wound healing seem to be ubiquitous in laboratory mice: indeed, we have found a significant *aggregate* contribution of IGE to rate of healing from an ear punch in all three mouse populations we have looked at to date (inbred C57BL/6J mice and outbred Heterogeneous Stock mice in Baud et al.(9), and CFW mice in this study). Thus, we were particularly interested in testing whether *Epha4* was involved in IGE on wound healing.

We found two additional significant IGE loci for FST immobility, more precisely for immobility during the last four minutes of the test (**Supplementary Table 3, Supplementary Figure 6a**). At the locus on chromosome 17, we identified eight genes as putatively causal but singled out *Dlgap1* (**Figure 4d**) for experimental validation because it encodes a synaptic protein(51), like *Epha4*, and because its expression in the hippocampus, which was measured in a separate cohort of 79 male CFW mice(45), was significantly and highly correlated with that of *Epha4* (Spearman r = 0.868, Bonferroni-corrected P = 2,3.10-19, **Supplementary Figure 6b**). As was the case for *Epha4*, we found no evidence of DGE arising from *Dlgap1* and affecting FST immobility in CFW mice (maximum -logP value at the locus 2.46).

### Evaluating the role of *Epha4* and *Dlgap1* in IGE using knockout models

We tested the hypotheses that *Epha4* can give rise to IGE on FST immobility and rate of healing using a constitutive *Epha4* knockout model on a mixed C56BL/6 & C56BL/10 genetic background. In addition, we tested for IGE from *Dlgap1* on FST immobility using a constitutive *Dlgap1* knockout model on a C57BL/6N background. At weaning, one *Epha4* mouse (heterozygote or wild-type, see Methods) or one *Dlgap1* mouse (homozygote knockout, heterozygote or wild-type) was co-housed with one focal FVB/NJ (FVB) mouse of the same sex (male or female). The FVB strain was chosen because it is the *inbred* strain whose genetic background is most similar to that of the *outbred* CFW mice used in igeGWAS, contributing 38% of all alleles in CFW mice(37). Focal FVB mice were ear punched prior to pairing, then the pairs of mice were left to interact in their cages for two months before they were all tested in the FST and the ears of FVB mice were analysed to measure the rate of healing (see Methods).

Although FVB mice are genetically similar to CFW mice, we observed that focal FVB mice showed much less immobility during the first two minutes of the FST than CFW mice (2.0 seconds on average across all FVB mice vs 12.2 seconds on average across all CFW mice). Therefore, in our analysis of FVB focal mice we focused on immobility during the last four minutes of the test, even though this measure showed a lower association in igeGWAS than immobility during the first two minutes of the test (-logP = 2.8 and 5.2 respectively).

When considering males and females together we found no effect of the genotype of cage mates on either FST immobility (P = 0.52, ANOVA, N = 81) or wound healing (P = 0.40, ANOVA, N = 85). However, model comparison using the Akaike Information Criterion (AIC) suggested there was an interaction between sex and genotype of the cage mate (i.e. IGE) for both FST immobility and wound healing, as the model including an interaction term between sex and genotype of the cage mate was favoured. Therefore, we considered the two sexes separately and observed, in males but not in females, IGE on FST immobility (P = 0.054, ANOVA, N = 35) and wound healing (P = 0.038, ANOVA, N = 38) (**Figure 5**). The detection of male-specific IGE from *Epha4* on wound healing is consistent with the observation of stronger IGE at the *Epha4* locus in male CFW mice compared to female CFW mice (**Supplementary Figure 7a**). The detection of male-specific IGE on FST immobility, on the other hand, was not expected from the analysis of CFW mice as similar effects were observed in males and females (**Supplementary Figures 7b and 7c**). A potential explanation for male-specific IGE on FST immobility in FVB focal mice is that FVB females showed lower immobility than FVB males, hindering our ability to detect genetic effects. Nevertheless, these experimental results support the hypothesis that *Epha4* can give rise to IGE on FST immobility and wound healing in laboratory mice.

**Figure 5.**
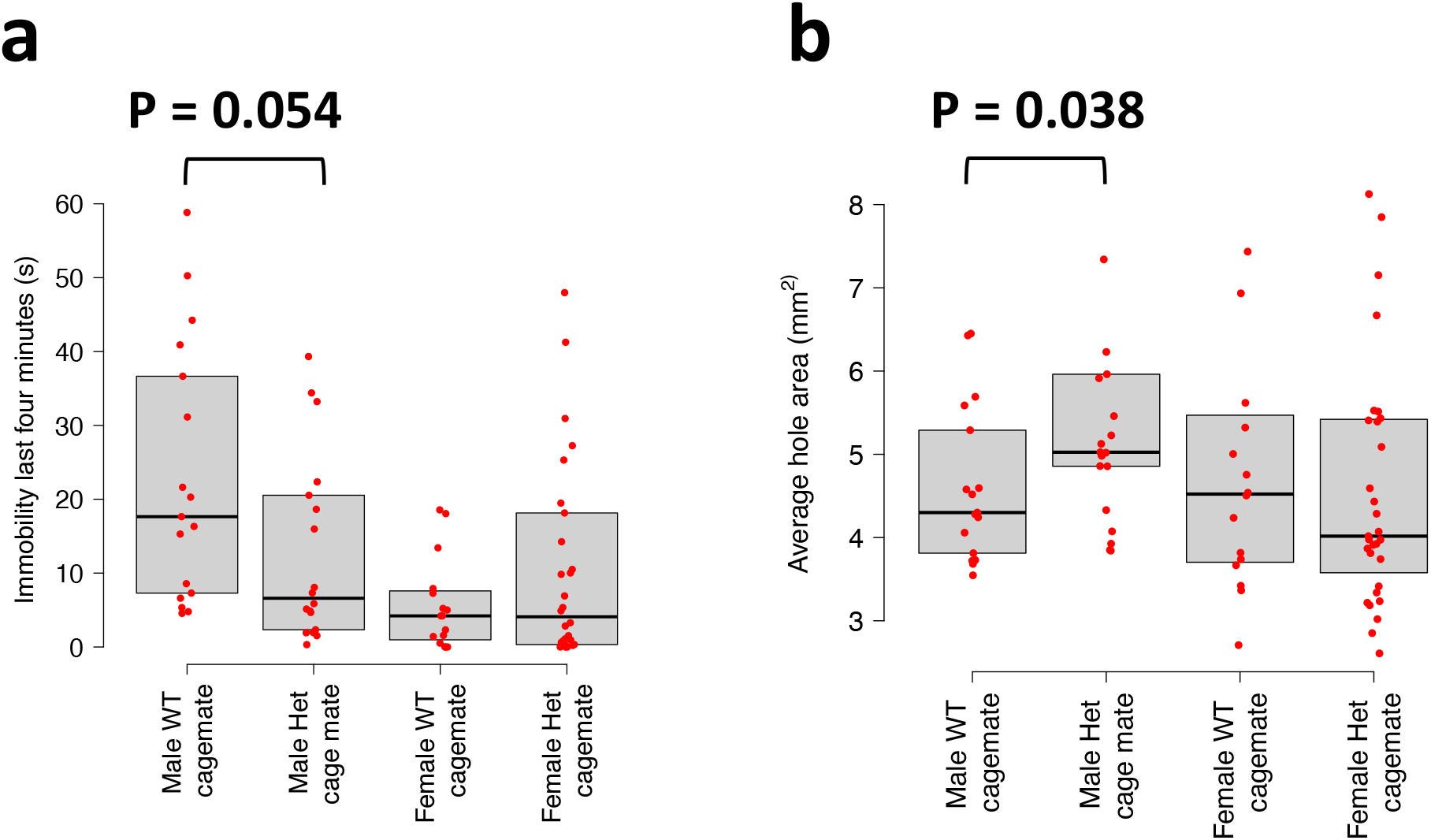
Results of an experiment in which FVB focal mice were co-housed with Epha4 knockout heterozygote (Het) or wild-type (WT) cage mates. Immobility during the last four minutes of the FST **(a)** and rate of healing from an ear punch **(b)** were measured in FVB focal mice after two months of co-housing.

As was the case in CFW mice, we did not observe a direct effect of *Epha4* on FST immobility whether all mice or males only were considered (P = 0.22 and 0.23 respectively, ANOVA, N = 81 and 35 respectively), indicating behavioural contagion is unlikely to explain these IGE.

Finally, we found no evidence of IGE from *Dlgap1* on FST immobility.

## Discussion

In this study, we leveraged a published dataset of 170 behavioural, physiological and morphological phenotypes measured in 1,812 genetically heterogeneous mice housed in same-sex groups of three to comprehensively assess the contribution of IGE to phenotypic variation and characterise the relationship between DGE and IGE for the same phenotype. Using polygenic models we showed that the genetic correlation ρ between DGE and IGE for a given phenotype is often significantly different from one, indicating IGE loci are different from DGE loci for the same phenotype. Consistently, we found that none of the 24 significant IGE loci identified for 17 phenotypes using igeGWAS overlapped with significant DGE loci identified using dgeGWAS. We fine-mapped seven significant IGE loci to a single putative causal gene and experimentally validated IGE from one of them, *Epha4*, on stress-coping strategy and wound healing using a knockout model.

The analysis of the genetic correlation ρ between DGE and IGE for the same phenotype provides insights into the overlap between DGE and IGE loci for a given phenotype and whether the traits mediating IGE on a phenotype of interest are genetically correlated (in the classical sense) with that phenotype. The correlation ρ was expected to be different from zero for many phenotypes, based on reports that emotions(52–54), behaviours(25, 55, 56), pathogens, and components of the gut microbiome(57) can “spread” between individuals and contribute to phenotypic variation, both in mice and in humans. In our study we found that ρ is significantly different from zero for a variety of phenotypes, which indicates some overlap between DGE loci and IGE loci for the same trait and is consistent with a genetic correlation (in the classical sense) between the phenotype of interest and the traits mediating IGE. However, we also found that ρ is significantly different from ±1 for ten out of twenty eight traits, reflecting differences between DGE and IGE loci and demonstrating that IGE on a phenotype of interest often involve traits of cage mates other than the phenotype of interest. This was true even for phenotypes that likely spread, namely stress and stress-coping strategies.

Consistent with the estimates of ρ from polygenic models, we found no overlap between the 24 loci identified by igeGWAS for 17 phenotypes and the loci identified by dgeGWAS for the same phenotypes. Our survey of a large number of phenotypes suggests that the loci identified by igeGWAS will, generally, be different from those identified by dgeGWAS, meaning igeGWAS holds great potential to uncover new loci underlying phenotypic variation and that these loci will point to traits of cage mates different from the phenotype studied.

Identifying IGE genes using igeGWAS has been previously attempted(5, 7, 8, 11, 31–35), but there has been limited evidence that this approach can indeed identify genes that are causally involved in IGE. The results of our igeGWAS and fine-mapping analyses identified a single putative causal gene at seven IGE loci: *Abca12* at a locus for adult neurogenesis, *Epha4* at a locus for stress-coping strategy, *Pkn2*, *Slit3* and Pgk1-rs7 at three different loci for sleep, *H60c* at a locus for home cage activity, and *Adcy1* at a locus for osteopetrosis. We tested one of these genes, *Epha4*, as well as another gene, *Dlgap1*, in experiments with knockout models. *Epha4* and *Dlgap1* were putative causal genes at two different IGE loci for stress-coping strategy and both encode synaptic proteins. However, only *Epha4* was at a locus with a single putative causal gene, making it a stronger candidate than *Dlgap1*. We confirmed the role of *Epha4* in giving rise to IGE on stress-coping strategy and wound healing in laboratory mice, but did not find evidence of IGE from *Dlgap1*. A limitation of our experiment is that FVB focal mice showed little to no immobility during the first two minutes of the FST, in contrast with the CFW mice used in igeGWAS. Hence, even though the significant igeGWAS locus was for immobility during the first two minutes of the test, we had to focus on immobility during the last four minutes when analysing the behaviour of FVB mice. Similarly, immobility during the last four minutes was lower in FVB female mice than it was in FVB male mice, which may explain why only observed IGE from *Epha4* in FVB male. Effects of the genetic background of knockout models have been reported in studies of DGE(58); our results show that in studies of IGE both the genetic background of the focal individuals matter too. In the future we will consider a broader range of genetic backgrounds for focal mice. The seven genes listed above as single putative causal genes at IGE loci as well as the experimental system we have developed to test *Epha4* and *Dlgap1* will serve as valuable starting points to gain further insights into the mechanisms of IGE in the future.

Finally, we identified challenges and solutions to different sources of confounding in igeGWAS. In particular, we demonstrated that correlations between direct and social genotypes arise when study individuals play both roles of focal individuals and social partners and that, counter-intuitively, these correlations arise even when all individuals are strictly unrelated. We showed that accounting for direct effects of the locus tested in the null model for igeGWAS permits avoiding spurious IGE associations. These insights, combined with the light we shed on two key parameters determining the power of igeGWAS, namely the number of cage mates and the mode of aggregation of IGE across cage mates, will inform the design and analysis of future igeGWAS.

## Conclusions

Our results demonstrate the potential for igeGWAS to uncover genetic effects expressed only in the context of social interactions and to serve as a starting point for follow up analyses and experiments that will improve our understanding of peer effects on health and disease.

## Methods

### Phenotypes and experimental variables

Phenotypes and experimental variables (covariates) for 1,934 male and female Crl:CFW(SW)-US_P08 (CFW) mice were retrieved from http://wp.cs.ucl.ac.uk/outbredmice/. Phenotypes were normalized using the boxcox function (MASS package(59)) in R; phenotypes that could not be normalised satisfactorily (transformation parameter lambda outside of −2 to 2 interval) were excluded. Because data for some phenotypes were missing for some mice, the sample size varied. The sample size for each phenotype after all filtering (see below) is indicated in **Supplementary Table 2**. The subset of covariates used for each phenotype, which always included sex, is indicated in **Supplementary Table 2**. For those phenotypes where body weight was included as a covariate, we checked that this did not lead to systematically increased (or decreased) estimates of the aggregate contribution of IGE (collider bias).

### Cage information

Mice were four to seven weeks old when they arrived at the phenotyping facility and were housed in same-sex groups of three mice. They were left undisturbed for nine to twelve weeks during their time in quarantine and spent another four weeks together during phenotyping.

Cage assignments were not included in the publicly available dataset but were provided by the authors upon request and are now provided in **Supplementary Table 1**. Cage assignments were recorded at eleven time points throughout the study and showed that a few mice were taken out of their original cages and singly housed, presumably because they were too aggressive. We only included in our analyses mice that had the same two cage mates throughout the experiment. We further excluded a subset of mice based on their genotype-based genetic similarity, as described below. Finally, all mice were singly housed during the sleep test and until sacrifice a few days later. Hence, we investigated “persistent” IGE on sleep and tissue phenotypes.

### Genome-wide genotypes

From http://wp.cs.ucl.ac.uk/outbredmice/ we retrieved both allele dosages for 7 million variants and allele dosages for a subset of 353,697 high quality, LD-pruned variants (as described in Nicod et al.(37); genotyping based on sparse sequencing data). We used LD-pruned variants for all analyses but the identification of putative causal genes at IGE loci (see below), for which we used the full set of variants.

### Genetic relatedness matrix (GRM) and exclusion of presumed siblings

The genetic relatedness matrix was calculated as the cross-product of the LD-pruned dosage matrix after standardizing the dosages for each variant to mean 0 and variance 1. A few pairs of mice were outliers in the distribution of GRM values, which made us suspect that siblings had been included in the sample even though they were not supposed to be (siblings were excluded by design). To mitigate confounding of DGE and IGE analyses by litter effects, we excluded 19 cages (57 mice) from all analyses.

### Variance components model

The same model as described in detail in Baud et al.(9) was used. Briefly, the model used is the following:

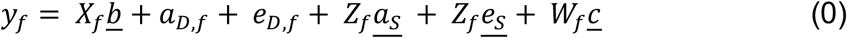

*y*_*f*_ is the phenotypic value of the focal mouse *f*, X_*f*_ is a row of the matrix *X* of covariate values and *b* a column vector of corresponding estimated coefficients. <u>*a_D,f_*</u> is the additive direct genetic effects (DGE) of *f*. Z_*f*_ is a row of the matrix *Z* that indicates cage mates (importantly Z_*i,i*_ = 0) and <u>*a_s_*</u> the column vector of additive indirect (social) genetic effects (IGE). <u>*e_D_*</u> refers to direct environmental effects (DEE) and <u>*e_s_*</u> to indirect
(social) environmental effects (IEE). *W_f_* is a row of the matrix *W* that indicates cage assignment and *c* the column vector of cage effects.

The joint distribution of all random effects is defined as:

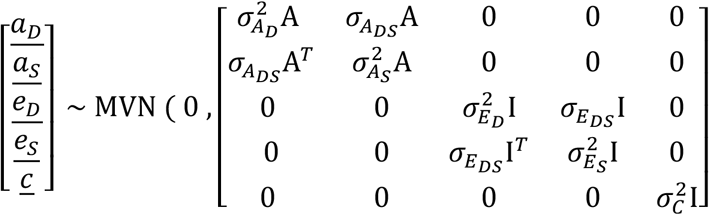

where A is the GRM matrix and I the identity matrix.

The phenotypic covariance is:

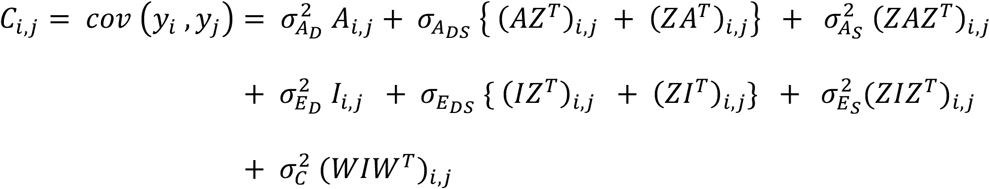

When all cages have the same number of mice, as is the case in this study, the non-genetic random effects are not identifiable(15, 60). An equivalent model can, in that case, be defined as(60):

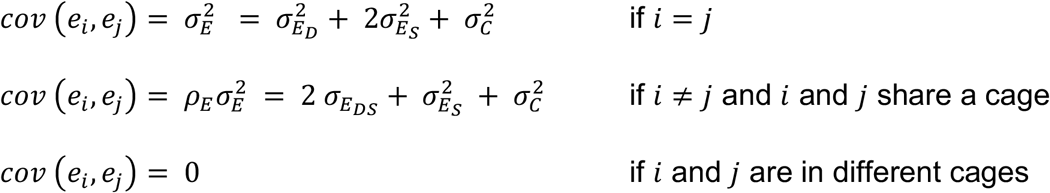

We checked that both model (0) and this alternative model yielded the same genetic estimates and maximum likelihoods. The alternative model was fitted using the SimplifNonIdableEnvs option in LIMIX(41, 61).

### Aggregate contributions of DGE and IGE

The aggregate contributions of DGE and IGE were calculated, respectively, as *sampleVar*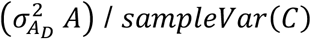 and *sampleVar*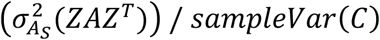, where *sampleVar* is the sample variance of the corresponding covariance matrix: suppose that we have a vector <u>*x*</u> of random variables with covariance matrix *M*, the sample variance of *M* is calculated as

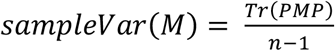

*T_r_* denotes the trace, *n* is the sample size, and 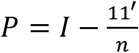.

Significance of the IGE variance component was assessed using a two-degree of freedom log likelihood ratio (LLR) test (for the variance component and the covariance with DGE). Note that this testing procedure is conservative. The Q value for the aggregate contribution of IGE was calculated for each phenotype using the R package qvalue(62). Significant IGE contributions were reported at FDR < 10% (corresponding to Q value < 0.1).

### Correlation between DGE and IGE

The correlation *ρ* between <u>*a_D_*</u> and <u>*a_s_*</u> was calculated as:

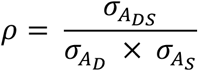

We tested whether *ρ* was significantly different from 0 and whether |ρ| was significantly different from 1 using a one-degree of freedom LLR test, which is conservative for the latter test.

### Simulations for Supplementary Figure 1

Phenotypes were simulated based on the genotypes and cage relationships of the full set of 1,812 mice. Phenotypes were drawn from model (0) with the following parameters: IGE explaining between 0 and 35.7% of phenotypic variance, DGE explaining 15% of phenotypic variance, 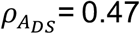, DEE explaining 22% of phenotypic variance, IEE explaining 16% of phenotypic variance, 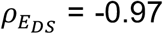, and cage effects explaining 26% of phenotypic variance. These variances correspond to the median value of estimates across traits with aggregate IGE and DGE > 5%. After building the phenotypic covariance matrix, the sample variance of the simulations was calculated and used to calculate “realised” simulation parameters from the “target” parameters above. The realised parameters were used for comparison with the parameters estimated from the simulations.

### Definition of “social genotype” for igeGWAS

We assumed additive effects across cage mates and calculated the “social genotype” of a mouse as the sum of the reference allele dosages of its cage mates. The same assumption was made by Biscarini *et al.(40)* and Brinker *et al.(31)* among others.

### Models used for igeGWAS and dgeGWAS

To test IGE of a particular variant in igeGWAS, we compared the following two models:

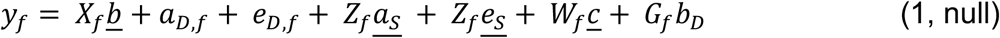

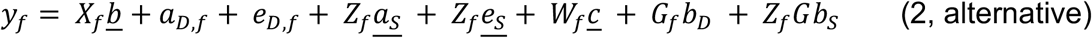

Here, *G* is the vector of direct genotypes at the tested variant; hence, *G_f_* is the genotype of the individual that is phenotyped (*f*) and *Z_f_G* is the sum of the genotypes of the two cage mates of *f*. *b_D_* the estimated coefficient for local DGE and *b_S_* the estimated coefficient for local IGE. Note that *Z_f_* could be defined as the average of the genotypes of the two cage mates of *f*, in which case <u>*b_S_*</u> would be doubled but the igeGWAS P values would remain unchanged. In igeGWAS, we refer to the inclusion of *G_f_b_D_* in model (1, null) as “conditioning”.

The models were fitted using LIMIX with the covariance of the model estimated only once per phenotype, in the null model with no local genetic effect (model 0).

The significance of local IGE was calculated by comparing models (1) and (2) with a 1-degree of freedom LLR test.

dgeGWAS was carried out by comparing model (2) above to the null model (3) below:

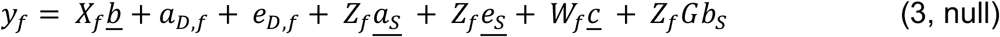

In dgeGWAS, we refer to the inclusion of *Z_f_Gb_S_* in model (3, null) as “conditioning”.

### Identification of significant associations

We used a genome-wide permutation strategy to control the FDR for each phenotype, as done by Nicod et al.(37). This strategy takes into account the specific patterns of linkage disequilibrium present in the sample and identifies significant associations *for each phenotype independently of the results for the other phenotypes in the dataset*. More precisely, for each phenotype and for each type of genetic effect (direct and indirect), we performed 100 “permuted GWAS” by permuting the rows of the matrix of social (respectively direct) genotypes, and testing each variant at a time using the permuted genotypes together with the un-permuted phenotypes, un-permuted covariates, un-permuted GRM and un-permuted matrix of direct (respectively social) genotypes (for conditioning)(41, 42). For a given P value x, the per-phenotype FDR can be calculated as:

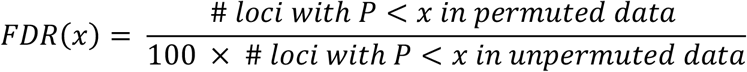

We reported those loci with FDR < 10%.

### Definition of putative causal genes at associated loci

At each significantly associated locus we defined a 1.5Mb window centred on the lead variant corresponding, in this sample, to the 95% confidence interval for the association(37). We identified all the variants that segregate in this window based on the full set of 7M variants and reran igeGWAS and dgeGWAS locally using all the variants at the locus. We defined “putative causal genes” as those genes that either overlapped the associated plateau or were located in direct proximity, and whose MGI symbol does not start by ‘Gm’, ‘Rik’, ‘Mir’, ‘Fam’, or ‘Tmem’ in order to focus on genes with known function and generate more tractable hypotheses on the pathways of social effects.

We identified putative causal genes using locusZoom plots(63). To create them, we used the standalone version of locusZoom (https://genome.sph.umich.edu/wiki/LocusZoom_Standalone). The plots for all 24 significant IGE loci reported in Supplementary Table 3 are provided in locusZooms_SupplTable3.zip.

### Gene expression in the hippocampus of an independent sample of CFW mice

Gene expression in the hippocampus of an independent sample of 79 male CFW mice, initially published in Parker et al.(45), was available from GeneNetwork (http://gn2.genenetwork.org/)(64, 65). The data are accessible by selecting Mouse as *Species*, CFW Outbred GWAS as *Group*, Hippocampus mRNA as *Type*, and UCSD CFW Hippocampus (Jan17) RNA-Seq Log2 Z-score as *Dataset*. To retrieve the genes whose expression is most highly correlated with that of *Epha4*, we entered “Epha4” in the *Get Any* field. Following selection of the Epha4 record (click on ENSMUSG00000026235), we used *Calculate Correlations* with Sample r as *Method*, UCSD CFW Hippocampus (Jan17) RNA-Seq Log2 Z-score as *Database*, and Spearman rank as correlation *Type*. **Supplementary Figure 6b** was obtained by clicking on the value of the correlation between Epha4 and Dlgap1 expression levels (column *Sample rho*).

### Variance explained by a significant association

The variance explained by a significant IGE association was estimated in an extension of model (0) with additional fixed effects for both direct and social effects of lead SNPs at all significant IGE loci (the lead SNP being the SNP with the most significant P value at the locus in the igeGWAS). After fitting the model, the variance was calculated as:

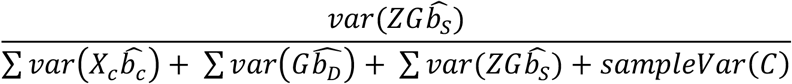

where *sampleVar(C)* is the sample variance of the covariance matrix in this model.

The variance explained by a significant DGE association was estimated in a similar model but considering all significant DGE associations and calculated as:

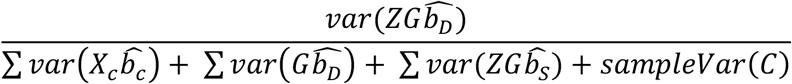

### Simulations for Supplementary Figure 2b and 2c

Phenotypes were simulated based on the genotypes and cage relationships of the 1,812 mice. Null phenotypes (no local IGE) were simulated from model (1) as the sum of random effects and local DGE. The following parameters were used for the random effects: 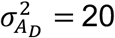 and 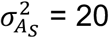(which correspond to high polygenic effects in the real data), 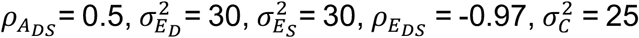 (which are close to the median of the corresponding estimates from the real data). Local DGE were simulated at random variants in the genome to account for 20% of the phenotypic variance.

### Simulations for Supplementary Figure 4

Phenotypes were simulated based on the real genotypes but random cages. Phenotypes were simulated as the sum of random and fixed effects using the following models:

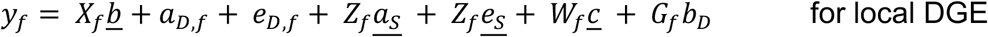

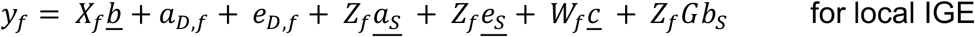

The following parameter values were used for the random effects: 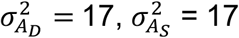 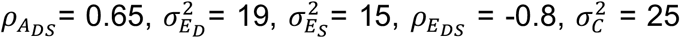. These values correspond to the median estimates for phenotypes with aggregate IGE and DGE > 0.1.

Local DGE and IGE were simulated at variants with low MAF (MAF < 0.05), medium MAF (0.225<MAF<0.275) or high MAF (MAF>0.45). Local IGE were simulated using two alternative generative models: an “additive” model by using *Z* as in model (2) (i.e. filled with 0s and 1s) or an “average” model by using 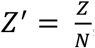, where N = 2. In all cases (DGE, additive IGE and average IGE) we simullated an allelic effect of 0.2, which is similar to the average allelic effect estimated in the igeGWAS. Power was calculated at a genome-wide significance threshold of negative log P 5, which is similar to the significance of associations detected at FDR < 10%.

### Experiment with *Epha4* and *Dlgap1* knockout mice

#### Experimental design

All animal procedures were approved by the Institutional Animal Care and Use Committee of the University of California San Diego (UCSD) and were conducted in accordance with the NIH Guide for the Care and Use of Laboratory Animals. FVB/NJ breeder mice were originally purchased from the Jackson Laboratory (Bar Harbor, MA, USA), then bred on site. *Epha4* knockout mice (allele name: Epha4tm1Byd) on a mixed C56BL/6 C56BL/10 genetic background, originally created by Dottori et al.(66), were generously donated by Prof. Elena Pasquale (Sanford Burnham Prebys, San Diego, CA, USA) then bred at UCSD. The mouse line C57BL/6N-Dlgap1<em1(IMPC)Tcp> was made as part of the KOMP2-Phase2 project at The Centre for Phenogenomics, Toronta, Canada. It was obtained from the Canadian Mouse Mutant Repository and bred at UCSD. Breeding from heterozygous parents produced, for *Dlgap1*, wild-type (WT), heterozygote (Het) and homozygote knockout (KO) offspring. For *Epha4*, homozygote knockout offspring usually died before weaning, leaving Het and WT offspring only. Within three days of weaning, we paired one focal FVB mouse with either a *Dlgap1* (WT, Het or KO) or an *Epha4* (WT or Het) cage mate of the same sex (male or female). Immediately prior to pairing, the FVB/NJ mice were ear punched on each ear using 2-mm ear punch scissors. Pairs of mice were then left to interact for two months before all mice were phenotyped in the forced swim test (FST), sacrificed and the ears of FVB/NJ mice were collected. The sample size was 52 *Epha4* Het mice and 33 *Epha4* WT mice for wound healing; for FST, there were only 48 *Epha4* Het mice as one mouse died during the FST, two mice had to be separated from their cage mate due to fighting in the days before the FST (but their ears were still collected as this did not significantly change the healing time), and the battery of the camera recording the FST ran out during the FST of the fourth mouse. A small subset of mice were video recorded in a new enclosure for 24h a few days before the FST but the data from this pilot project are not reported here. Throughout the experiment all mice were housed on a 12h:12h light-dark cycle, with lights on at 06:00, and all behavioural testing occurred during the light phase of the light-dark cycle.

#### Forced swim test

Following the same protocol as in the CFW study(37), mice were tested in the forced swim test: they were placed for 6 minutes in 6’’ wide × 12’’ tall glass buckets filed with water at 24-26°C. Mice were video recorded from the side and their immobility during the first 2 and last 4 minutes of the test was scored by an observer blind to the genotypes of the black (*Epha4* and *Dlgap1*) mice. The analysis of IGE focused on immobility of FVB mice in the last four minutes of the test as FVB mice are rarely immobile during the first two minutes of the test.

#### Healing from an ear punch

Both ears of FVB/NJ mice were punched with a 2mm-diameter ear punch scissor just before the mice were paired with an *Epha4* or a *Dlgap1* cage mate at weaning. Following the same protocol as in the CFW study(37), the ears were collected two months later after sacrifice, stored in 10% buffered formalin phosphate until analysis. To measure the area of the hole, each ear was mounted on an histology slide and photos were taken from a fixed distance. Images were analysed with the ImageJ software(67) and the average across the two ears calculated.

#### Genotyping

For genotyping *Epha4* mice, tail or ear biopsies were sent to Transnetyx Inc. for genotyping (Transnetyx Genotyping Services, Cordova, TN). Transnetyx Inc. utilize real-time PCR and duplicate sample processing to ensure the accuracy of each mutation. Additionally, Sanger sequencing was performed to further validate the results of the Transnetyx assays.

For genotyping *Dlgap1* mice, we used a multiplex PCR with primers: *CCGTAAGTGAAGTCTCCATCAACAG (Fw1), CGGCTAGGATTTCAGAGTTTGTTC* (Fw2) and *CTTCCTCTCCTACACCATCAACAC (Rev1)*, yielding a 308bp band in the presence of a WT allele and a 392bp band in the presence of a knockout allele.

#### Statistical analysis

For both FST immobility and wound healing, five fixed-effect models were first compared using AIC: a model with intercept only, a model with the sex of the pair (focal animal and cage mate were always of the same sex), a model with the genotype of the cage mate, a model with both sex and genotype of cage mate, and finally a model with main effects of sex and genotype of cage mate and their interaction.

IGE were then tested in males only using an analysis of variance (ANOVA) with one degree of freedom.

## Supporting information

locusZoom plots for all significant IGE loci

## Declarations

### Ethics approval

All animal procedures were approved by the Institutional Animal Care and Use Committee of the University of California San Diego (UCSD) and were conducted in accordance with the NIH Guide for the Care and Use of Laboratory Animals.

### Availability of data and materials

Genotype and phenotype data from Nicod et al.(37) and Davies et al.(38) are available from http://wp.cs.ucl.ac.uk/outbredmice/. Cage information is provided in Supplementary Table 1.

All the scripts used in this study are available from http://github.com/limix/IGE. LIMIX can be downloaded from http://github.com/limix/limix.

### Authors’ contributions

AB, AAP and OS designed the study. AB and FPC performed the analyses. JN contributed the cage information. AB, ABL, NF, and CM performed the mouse knockout experiments. All authors contributed to the interpretation of the data. AB, FPC, AAP and OS wrote the manuscript.

## Acknowledgements

The authors want to thank Drs. Na Cai, Robert W. Davies and Richard Mott for facilitating access to the genotype data. The authors also thank Dr. Elena Pasquale for generously providing mice from the *EphA4* knockout line, which was produced thanks to funding from NIH grant NS087070.

## References

1. Griffing B. Selection in reference to biological groups I. Individual and group selection applied to populations of unordered groups. Australian Journal of Biological Sciences. 1967;20(1):127–40.

2. Moore AJ, Brodie III ED, Wolf JB. Interacting phenotypes and the evolutionary process: I. Direct and indirect genetic effects of social interactions. Evolution. 1997;51(5):1352–62.

3. Wolf JB, Brodie III ED, Cheverud JM, Moore AJ, Wade MJ. Evolutionary consequences of indirect genetic effects. Trends in ecology & evolution. 1998;13(2):64–9.

4. Wolf JB, Mutic JJ, Kover PX. Functional genetics of intraspecific ecological interactions in Arabidopsis thaliana. Philosophical Transactions of the Royal Society B: Biological Sciences. 2011;366(1569):1358–67.

5. Bailey NW, Hoskins JL. Detecting cryptic indirect genetic effects. Evolution. 2014;68(7):1871–82.

6. Bleakley BH, Brodie ED. Indirect genetic effects influence antipredator behavior in guppies: estimates of the coefficient of interaction psi and the inheritance of reciprocity. Evolution. 2009;63(7):1796–806.

7. Ashbrook DG, Gini B, Hager R. Genetic variation in offspring indirectly influences the quality of maternal behaviour in mice. Elife. 2015;4:11814.

8. Ashbrook DG, Sharmin N, Hager R, editors. Offspring genes indirectly influence sibling and maternal behavioural strategies over resource share. Proc R Soc B; 2017: The Royal Society.

9. Baud A, Mulligan MK, Casale FP, Ingels JF, Bohl CJ, Callebert J, et al. Genetic variation in the social environment contributes to health and disease. PLoS genetics. 2017;13(1):e1006498.

10. Head ML, Berry LK, Royle NJ, Moore AJ. Paternal care: direct and indirect genetic effects of fathers on offspring performance. Evolution. 2012;66(11):3570–81.

11. Mutic JJ, Wolf JB. Indirect genetic effects from ecological interactions in Arabidopsis thaliana. Molecular ecology. 2007;16(11):2371–81.

12. Petfield D, Chenoweth SF, Rundle HD, Blows MW. Genetic variance in female condition predicts indirect genetic variance in male sexual display traits. Proceedings of the National Academy of Sciences. 2005;102(17):6045–50.

13. Wilson AJ, Gelin U, Perron M-C, Réale D. Indirect genetic effects and the evolution of aggression in a vertebrate system. Proceedings of the Royal Society of London B: Biological Sciences. 2009;276(1656):533–41.

14. Moore AJ, Haynes KF, Preziosi RF, Moore PJ. The evolution of interacting phenotypes: genetics and evolution of social dominance. the american naturalist. 2002;160(S6):S186–S97.

15. Bergsma R, Kanis E, Knol EF, Bijma P. The contribution of social effects to heritable variation in finishing traits of domestic pigs (Sus scrofa). Genetics. 2008;178(3):1559–70.

16. Ellen ED, Visscher J, van Arendonk JA, Bijma P. Survival of laying hens: genetic parameters for direct and associative effects in three purebred layer lines. Poultry science. 2008;87(2):233–9.

17. Alemu SW, Bijma P, Møller SH, Janss L, Berg P. Indirect genetic effects contribute substantially to heritable variation in aggression-related traits in group-housed mink (Neovison vison). Genetics Selection Evolution. 2014;46(1):30.

18. e Silva JC, Potts B, Gilmour A, Kerr R. Genetic-based interactions among tree neighbors: identification of the most influential neighbors, and estimation of correlations among direct and indirect genetic effects for leaf disease and growth in Eucalyptus globulus. Heredity. 2017;119(3):125.

19. Wilson A, Coltman D, Pemberton J, Overall A, Byrne K, Kruuk L. Maternal genetic effects set the potential for evolution in a free‐living vertebrate population. Journal of evolutionary biology. 2005;18(2):405–14.

20. McAdam AG, Boutin S, Réale D, Berteaux D. Maternal effects and the potential for evolution in a natural population of animals. Evolution. 2002;56(4):846–51.

21. Hadfield JD, Burgess MD, Lord A, Phillimore AB, Clegg SM, Owens IP. Direct versus indirect sexual selection: genetic basis of colour, size and recruitment in a wild bird. Proceedings of the Royal Society B: Biological Sciences. 2006;273(1592):1347–53.

22. Kong A, Thorleifsson G, Frigge ML, Vilhjalmsson BJ, Young AI, Thorgeirsson TE, et al. The nature of nurture: Effects of parental genotypes. Science. 2018;359(6374):424–8.

23. Bates TC, Maher BS, Medland SE, McAloney K, Wright MJ, Hansell NK, et al. The Nature of Nurture: Using a Virtual-Parent Design to Test Parenting Effects on Children’s Educational Attainment in Genotyped Families. Twin Research and Human Genetics. 2018:1–11.

24. Warrington NM, Beaumont RN, Horikoshi M, Day FR, Helgeland Ø, Laurin C, et al. Maternal and fetal genetic effects on birth weight and their relevance to cardio-metabolic risk factors. Nature genetics. 2019:1.

25. Sotoudeh R, Harris KM, Conley D. Effects of the peer metagenomic environment on smoking behavior. Proceedings of the National Academy of Sciences. 2019;116(33):16302–7.

26. Xia C, Canela-Xandri O, Rawlik K, Tenesa A. Evidence of horizontal indirect genetic effects in humans. Nature Human Behaviour. 2021;5(3):399–406.

27. Demange PA, Hottenga JJ, Abdellaoui A, Malanchini M, Domingue BW, de Zeeuw EL, et al. Parental influences on offspring education: indirect genetic effects of non-cognitive skills. biorxiv. 2020.

28. McGlothlin JW, Brodie III ED. HOW TO MEASURE INDIRECT GENETIC EFFECTS: THE CONGRUENCE OF TRAIT‐BASED AND VARIANCE‐PARTITIONING APPROACHES. Evolution. 2009;63(7):1785–95.

29. Bijma P. The quantitative genetics of indirect genetic effects: a selective review of modelling issues. Heredity. 2014;112(1):61.

30. Wilson AJ, Morrissey M, Adams M, Walling CA, Guinness FE, Pemberton JM, et al. Indirect genetics effects and evolutionary constraint: an analysis of social dominance in red deer, Cervus elaphus. Journal of evolutionary biology. 2011;24(4):772–83.

31. Brinker T, Bijma P, Vereijken A, Ellen ED. The genetic architecture of socially-affected traits: a GWAS for direct and indirect genetic effects on survival time in laying hens showing cannibalism. Genetics Selection Evolution. 2018;50(1):38.

32. Wu P, Wang K, Yang Q, Zhou J, Chen D, Liu Y, et al. Whole-genome re-sequencing association study for direct genetic effects and social genetic effects of six growth traits in Large White pigs. Scientific reports. 2019;9(1):1–12.

33. Hong J-K, Lee J-B, Ramayo-Caldas Y, Kim S-D, Cho E-S, Kim Y-S, et al. Single-step genome-wide association study for social genetic effects and direct genetic effects on growth in Landrace pigs. Scientific reports. 2020;10(1):1–11.

34. Warrington NM, Beaumont RN, Horikoshi M, Day FR, Helgeland Ø, Laurin C, et al. Maternal and fetal genetic effects on birth weight and their relevance to cardio-metabolic risk factors. Nature genetics. 2019;51(5):804–14.

35. McGinnis R, Steinthorsdottir V, Williams NO, Thorleifsson G, Shooter S, Hjartardottir S, et al. Variants in the fetal genome near FLT1 are associated with risk of preeclampsia. Nature genetics. 2017;49(8):1255.

36. Yalcin B, Nicod J, Bhomra A, Davidson S, Cleak J, Farinelli L, et al. Commercially available outbred mice for genome-wide association studies. PLoS genetics. 2010;6(9):e1001085.

37. Nicod J, Davies RW, Cai N, Hassett C, Goodstadt L, Cosgrove C, et al. Genome-wide association of multiple complex traits in outbred mice by ultra-low-coverage sequencing. Nature genetics. 2016;48(8):912.

38. Davies RW, Flint J, Myers S, Mott R. Rapid genotype imputation from sequence without reference panels. Nature genetics. 2016;48(8):965.

39. Sadler AM, Bailey SJ. Repeated daily restraint stress induces adaptive behavioural changes in both adult and juvenile mice. Physiology & behavior. 2016;167:313–23.

40. Biscarini F, Bovenhuis H, Van Der Poel J, Rodenburg T, Jungerius A, Van Arendonk J. Across-line SNP association study for direct and associative effects on feather damage in laying hens. Behavior genetics. 2010;40(5):715–27.

41. Casale FP, Rakitsch B, Lippert C, Stegle O. Efficient set tests for the genetic analysis of correlated traits. Nature methods. 2015;12(8):755.

42. Sudmant PH, Rausch T, Gardner EJ, Handsaker RE, Abyzov A, Huddleston J, et al. An integrated map of structural variation in 2,504 human genomes. Nature. 2015;526(7571):75.

43. Beavis W. QTL analyses: power, precision and accuracy. Molecular Dissection of Complex Traits. Edited by: AH P. 1997. CRC Press, New York.

44. Beavis W, Beavis W. The power and deceit of QTL experiments: lessons from comparative QTL studies. 1994.

45. Parker CC, Gopalakrishnan S, Carbonetto P, Gonzales NM, Leung E, Park YJ, et al. Genome-wide association study of behavioral, physiological and gene expression traits in outbred CFW mice. Nature genetics. 2016;48(8):919.

46. Commons KG, Cholanians AB, Babb JA, Ehlinger DG. The Rodent Forced Swim Test Measures Stress-Coping Strategy, Not Depression-like Behavior. ACS Chemical Neuroscience. 2017;8(5):955–60.

47. Bouvier D, Corera AT, Tremblay MÈ, Riad M, Chagnon M, Murai KK, et al. Pre‐ synaptic and post‐synaptic localization of EphA4 and EphB2 in adult mouse forebrain. Journal of neurochemistry. 2008;106(2):682–95.

48. Murai KK, Nguyen LN, Irie F, Yamaguchi Y, Pasquale EB. Control of hippocampal dendritic spine morphology through ephrin-A3/EphA4 signaling. Nature neuroscience. 2003;6(2):153–60.

49. Li Y, Wang H, Wang X, Liu Z, Wan Q, Wang G. Differential expression of hippocampal EphA4 and ephrinA3 in anhedonic-like behavior, stress resilience, and antidepressant drug treatment after chronic unpredicted mild stress. Neuroscience letters. 2014;566:292–7.

50. Zhang J-c, Yao W, Qu Y, Nakamura M, Dong C, Yang C, et al. Increased EphA4-ephexin1 signaling in the medial prefrontal cortex plays a role in depression-like phenotype. Scientific Reports. 2017;7(1):7133.

51. Coba M, Ramaker M, Ho E, Thompson S, Komiyama N, Grant S, et al. Dlgap1 knockout mice exhibit alterations of the postsynaptic density and selective reductions in sociability. Scientific reports. 2018;8(1):1–12.

52. de Oliveira PC, Zaniboni CR, Carmona IM, Fonseca AR, Canto-de-Souza A. Preliminary behavioral assessment of cagemates living with conspecifics submitted to chronic restraint stress in mice. Neuroscience letters. 2017;657:204–10.

53. Bolger N, DeLongis A, Kessler RC, Wethington E. The contagion of stress across multiple roles. Journal of Marriage and the Family. 1989:175–83.

54. Haeffel GJ, Hames JL. Cognitive Vulnerability to Depression Can Be Contagious. Clinical Psychological Science. 2014;2(1):75–85.

55. Stickney JD, Morgan MM. Social housing promotes recovery of wheel running depressed by inflammatory pain and morphine withdrawal in male rats. Behavioural Brain Research. 2021;396:112912.

56. Clarke T-K, Adams MJ, Howard DM, Xia C, Davies G, Hayward C, et al. Genetic and shared couple environmental contributions to smoking and alcohol use in the UK population. Molecular psychiatry. 2019:1–11.

57. Ridaura VK, Faith JJ, Rey FE, Cheng J, Duncan AE, Kau AL, et al. Gut microbiota from twins discordant for obesity modulate metabolism in mice. Science. 2013;341(6150):1241214.

58. Sittig LJ, Carbonetto P, Engel KA, Krauss KS, Barrios-Camacho CM, Palmer AA. Genetic background limits generalizability of genotype-phenotype relationships. Neuron. 2016;91(6):1253–9.

59. Venerables W, Ripley B. Modern applied statistics with S. new york: Springer; 2002.

60. Bijma P, Muir WM, Ellen ED, Wolf JB, Van Arendonk JA. Multilevel selection 2: estimating the genetic parameters determining inheritance and response to selection. Genetics. 2007;175(1):289–99.

61. Lippert C, Casale F, Rakitsch B, Stegle O. LIMIX: genetic analysis of multiple traits. 2014.

62. Dabney A, Storey JD, Warnes G. qvalue: Q-value estimation for false discovery rate control. R package version. 2010;1(0).

63. Pruim RJ, Welch RP, Sanna S, Teslovich TM, Chines PS, Gliedt TP, et al. LocusZoom: regional visualization of genome-wide association scan results. Bioinformatics. 2010;26(18):2336–7.

64. Mulligan MK, Mozhui K, Prins P, Williams RW. GeneNetwork: a toolbox for systems genetics. Systems Genetics: Springer; 2017. p. 75–120.

65. Sloan Z, Arends D, Broman KW, Centeno A, Furlotte N, Nijveen H, et al. GeneNetwork: framework for web-based genetics. The Journal of Open Source Software. 2016;1(2):-.

66. Dottori M, Hartley L, Galea M, Paxinos G, Polizzotto M, Kilpatrick T, et al. EphA4 (Sek1) receptor tyrosine kinase is required for the development of the corticospinal tract. Proceedings of the National Academy of Sciences. 1998;95(22):13248–53.

67. Schneider CA, Rasband WS, Eliceiri KW. NIH Image to ImageJ: 25 years of image analysis. Nature methods. 2012;9(7):671–5.

