## Supplementary Note and Supplementary Figures for "Dissecting indirect genetic effects from peers in laboratory mice"

### Supplementary Note: Correlation between direct and social genotypes arising from using each individual as both focal individual and social partner.

A common analysis strategy to maximize sample size when all individuals are genetically heterogeneous and have been both genotyped and phenotyped is to consider each individual as both focal individual and social partner in the model. In this Supplementary Note we investigate the calibration of P values under the null (i.e. no IGE) under this design, when the tested locus gives rise to DGE. We simulate **strictly unrelated individuals**.

Simulations: $N$ = 1,000 individuals were randomly assigned to groups of two (pairs). Direct genotypes at a locus were sampled from the binomial distribution $B\ (2, 0.02)$ (the average minor allele frequency in CFW mice is 0.019[1]) and the social genotype of an individual was defined as the direct genotype of its group mate. Phenotypes were generated from DGE at that one locus plus independent and identically distributed (iid) noise. Local DGE were set to explain a certain fraction of the total variance (vg), which varied from 0 to 80% in the simulations. No IGE were simulated at the locus, nor any effect of genetic background or cage effects. 10,000 phenotypes were simulated for each combination of vg and conditioning or not.

Analysis of the simulated phenotypes: Each simulated phenotype was analysed using a standard linear model, testing for IGE arising from social genotypes at the locus, where social genotype is defined as the genotype of the group mate. Additionally, the model included a fixed effect for DGE (a strategy we refer to as "conditioning", bottom plots) or not (top plots). No random effect was included for genetic background or cage effects.

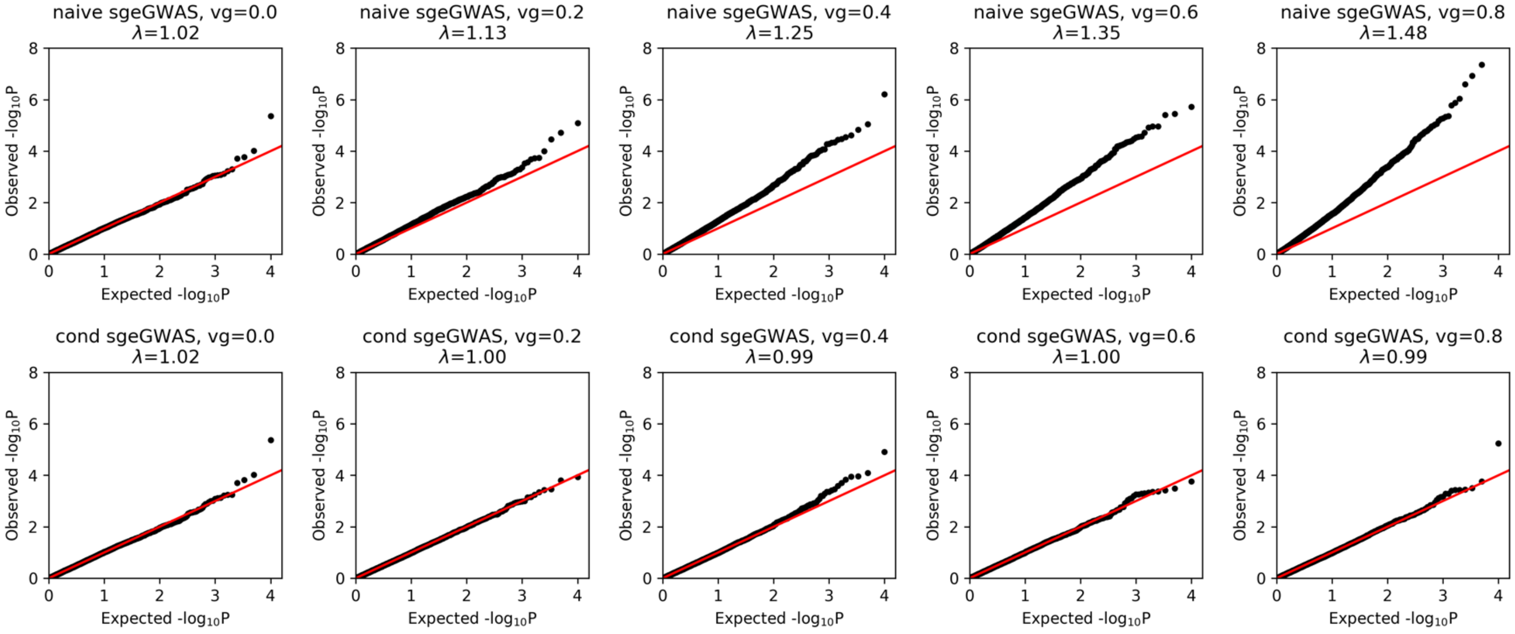


***Note Figure 1** QQ plots of IGE negative log P values. Null phenotypes (no IGE) were simulated from DGE at one locus and iid noise. IGE P values were obtained from models including a fixed effect for DGE at the locus (bottom plots) or not (top plots). The fraction of the total variance explained by the simulated DGE (vg) varied from 0 to 80% in the simulations. Each QQ shows 10,000 P values, one for each phenotype in the simulation set.*

<u>Results</u>: In null simulations (no IGE), strong DGE (high vg) lead to strong inflation of IGE P values in the absence of conditioning. This inflation is resolved (i.e. P values are calibrated) by conditioning on the direct genotype at the locus.

<u>Investigation of the origin of the inflation</u>: In the presence of DGE at a locus, the P value for the IGE test depends on the correlation between direct and social genotypes. We found that, even when all individuals are strictly unrelatedd (as simulated above), the correlation between direct and social genotypes takes more extreme values than the correlation between two independent draws of length $N$ from $B\,(2, 0.02)$ (Note Figure 2).

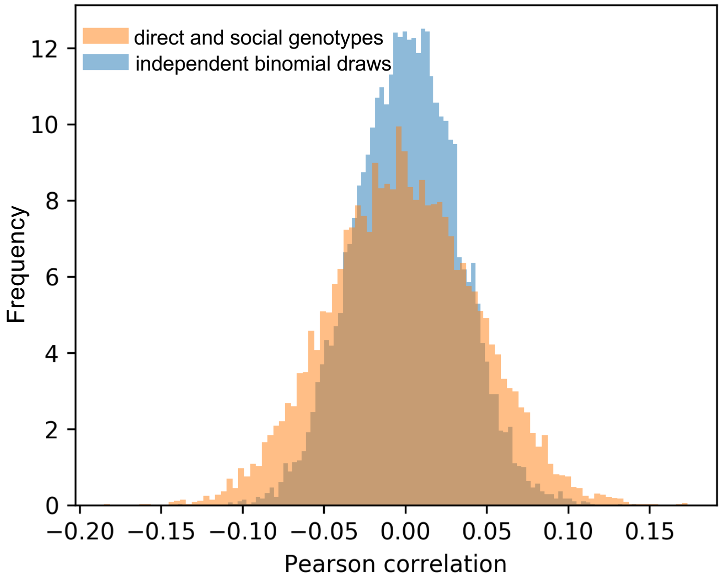


***Note Figure 2*** *Distribution of the Pearson correlation between direct and social genotypes (orange) and independent binomial draws. Direct genotypes, social genotypes, and the independent binomial draws are all of length $N$ = 1,000 (number of individuals).*

We now mathematically explain this observation. When $N$ strictly unrelated individuals are housed in pairs and each individual serves as both focal individual and social partner in the analysis, direct and social genotypes ($G_D$ and $G_S$ respectively) can be re-written as $G_D = [a, b]$ and $G_S = [b, a]$, where $a$ and $b$ are independent and identically distributed vectors of length $N/2$. Without loss of generality, let's assume that $E[a] = E[b] = 0$ and $Var[a] = Var[b] = 1$. Then we have:

$$Var[Corr(G_D, G_s)] \approx \frac{Var[G_D.G_S]}{N^2} \approx \frac{Var[2 \times a.b]}{N^2} \approx \frac{4\times(\frac{N}{2})}{N^2} \approx \frac{2}{N}.$$

In contrast, for independent vectors $X_1$ and $X_2$ of the same size as $G_D$ and $G_S$ (i.e. size $N$), we have:

$Var[Corr(X_1, X_2)] \approx \frac{Var[X_1.X_2]}{N^2} \approx \frac{1}{N}$.

Conclusion: When individuals serve as both focal individual and social partner and to the extent that DGE affect a phenotype, these results show the need for conditioning on direct genotypes in igeGWAS in order to obtain calibrated P values (under the null hypothesis of no IGE), **even when all individuals are strictly unrelated**.

Similarly, when individuals serve as both focal individual and social partner (which is not usually the case in human GWAS for example) and to the extent that IGE affect the phenotype, conditioning on social genotypes is required to obtain calibrated dgeGWAS P values, **even when all individuals are strictly unrelated**.

We adopt conditioning in all the analyses presented in the main text.

Code: The code used for this Supplementary Note is provided as an IPython Notebook (SupplNote.ipynb).

### Supplementary Figures

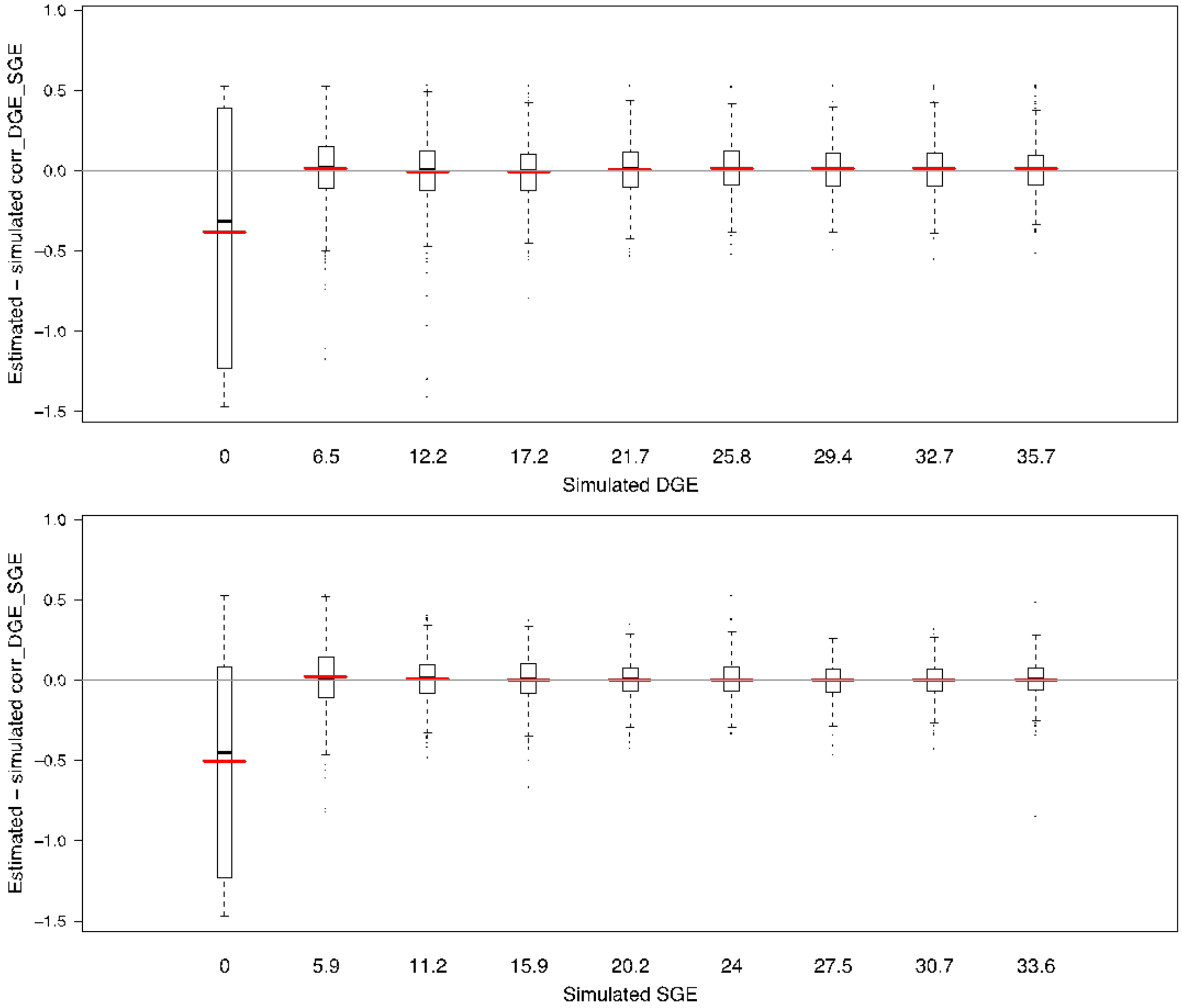


***Supplementary Figure 1*** *Estimation of the correlation between IGE and DGE (ρ, see Figure 1b) in simulations. The difference between the estimated and the true (simulated) value of ρ is shown (y axis) for various aggregate DGE (top, x axis) and IGE (bottom, x axis).*

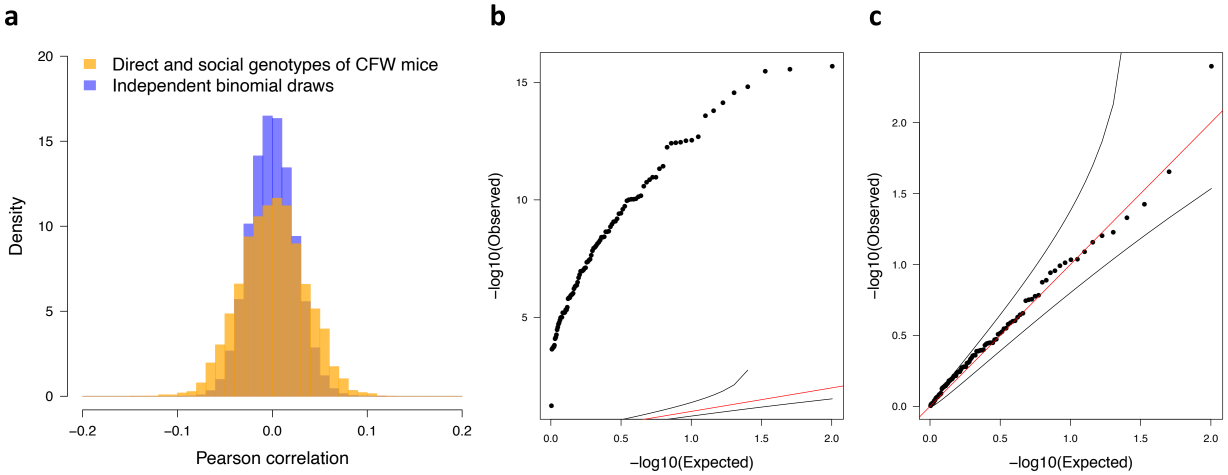


***Supplementary Figure 2*** *Correlations between direct and social genotypes of CFW mice, and implications for GWAS. (a) Pearson correlation between direct and social genotypes at each of the LD-pruned variants used in GWAS (orange) and between independent binomial draws with same minor allele frequency and length (blue). (b,c) QQ plot of IGE negative log P values from simulations where no IGE were simulated but large-effect DGE at the tested variant were simulated: genotypic correlations not accounted for (i.e. no conditioning) in panel b, genotypic correlations accounted for by conditioning in panel c.*

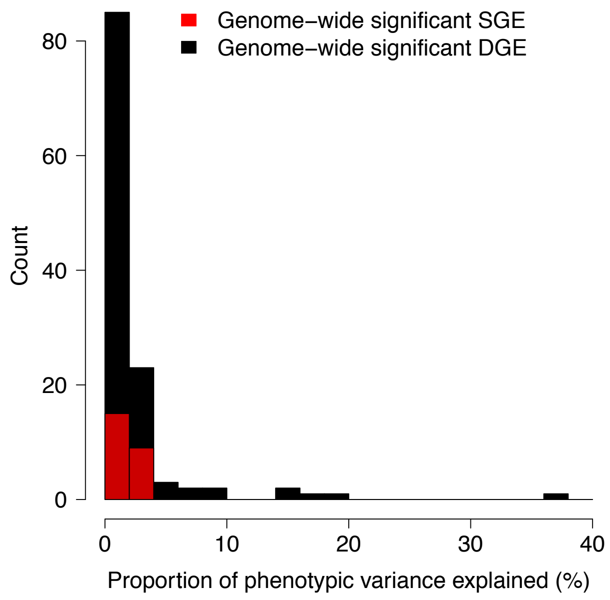


***Supplementary Figure 3*** *Proportion of phenotypic variance explained by significant DGE (black) and IGE (red) associations (FDR < 10%).*

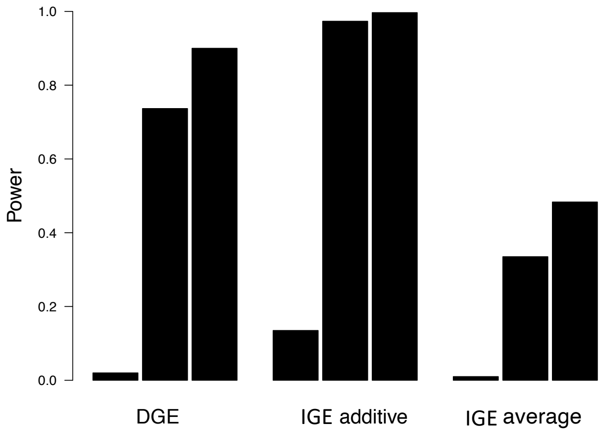


***Supplementary Figure 4*** *Power to detect DGE and IGE associations in simulations. Three types of local genetic effects were simulated: DGE, IGE arising from the sum of genetic effects across the two cage mates (additive model) and IGE arising from the average of genetic effects across the two cage mates (average model). The average model corresponds to a scenario where a mouse can only interact with one cage mate at a time, resulting in a dilution of social effects when more than one cage mate is present in the cage (two cage mates were present in this study). For each type of effect, results are shown (left to right) for variants with low MAF (MAF < 0.05), medium MAF (0.225<MAF<0.275) and high MAF (MAF>0.45) (MAF: minor allele frequency, defined based on direct genotypes). Power was calculated at a genome-wide significance threshold of negative log P 5.*

*To understand why the power varies the way it does, remember that the sample variances of the local genetic terms are the following:*

- $var(Gb_D) = 2p(1-p)\, b_D^2$ *for DGE*
- $var(ZGb_S) = 2Np(1-p)\, b_S^2$ *with N=2 for IGE simulated under the additive model*
- $var\left(\frac{Z}{N}Gb_S\right) = \frac{2Np(1-p)}{N^2} b_S^2$ *for IGE simulated under the proportional model.*

*Also note that the sample variance of the DGE term is the same as the sample variance of an IGE term with $N = 1$.*

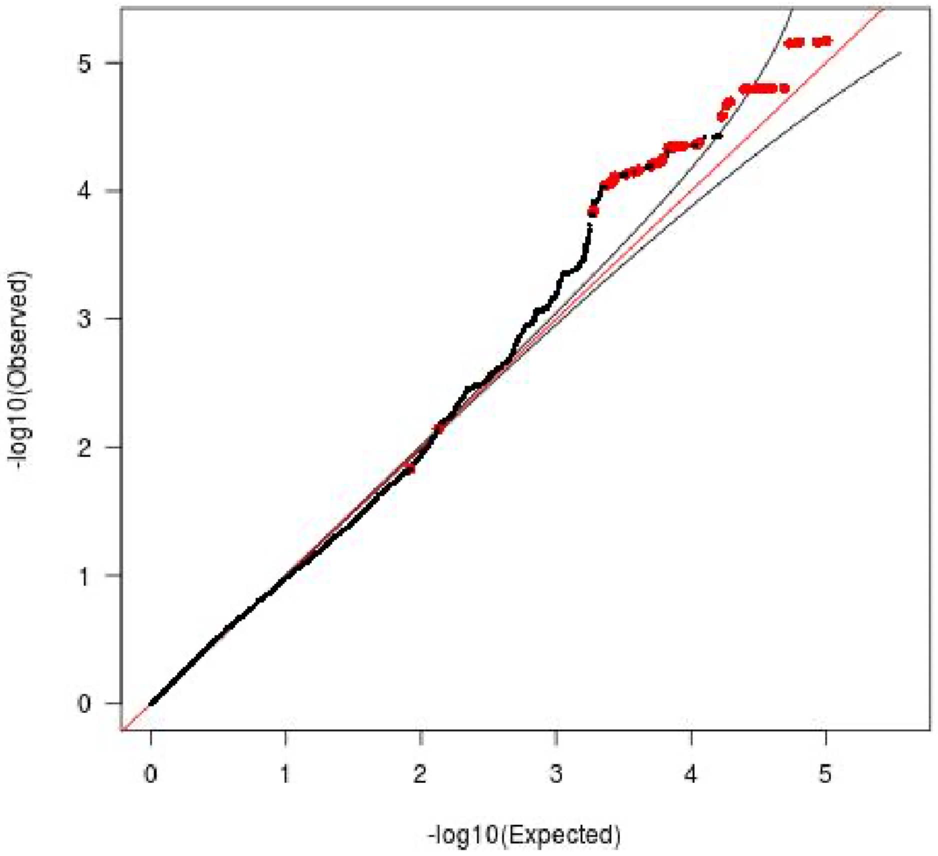


***Supplementary Figure 5*** *Quantile-quantile (QQ) plot for the igeGWAS P values for Immobility during the first two minutes of the forced swim test. The red colored dots correspond to variants that are in linkage disequilibrium ($R^2$ > 0.5) with the top variant at the significant IGE locus on chromosome 1 (FDR < 10%) for that measure. Epha4 is the only putative causal gene at this locus.*

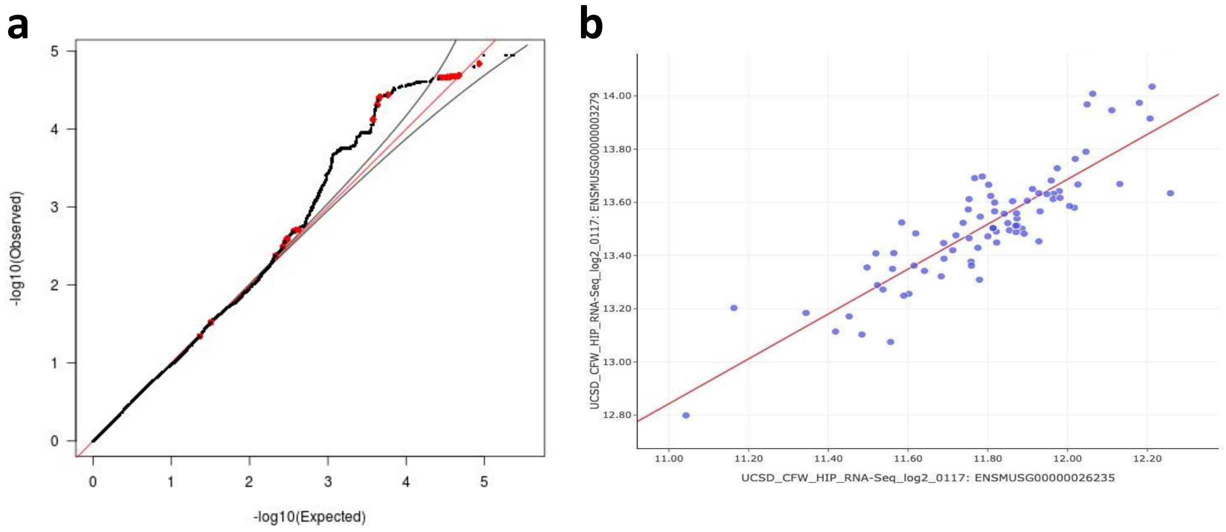


***Supplementary Figure 6*** *Information relevant to the role of Dlgap1 in giving rise to IGE on immobility during the last four minutes of the FST. (a) Quantile-quantile (QQ) plot of igeGWAS P values, showing in red the variants that are in linkage disequilibrium (*$R^2$ *> 0.5) with the top variant at the significant IGE locus on chromosome 17 (FDR < 10%), where Dlgap1 was identified as one of eight putative causal genes. The dots with observed -logP values greater than 3 that are not red correspond to variants that are at the other significant IGE locus, on chromosome 9, for that phenotype. (b) Correlation between the expression of Epha4 (x-axis) and Dlgap1 (y-axis) in the hippocampus of a dataset collected in an independent cohort of 79 male CFW mice (Figure from Genenetwork, see Methods).*

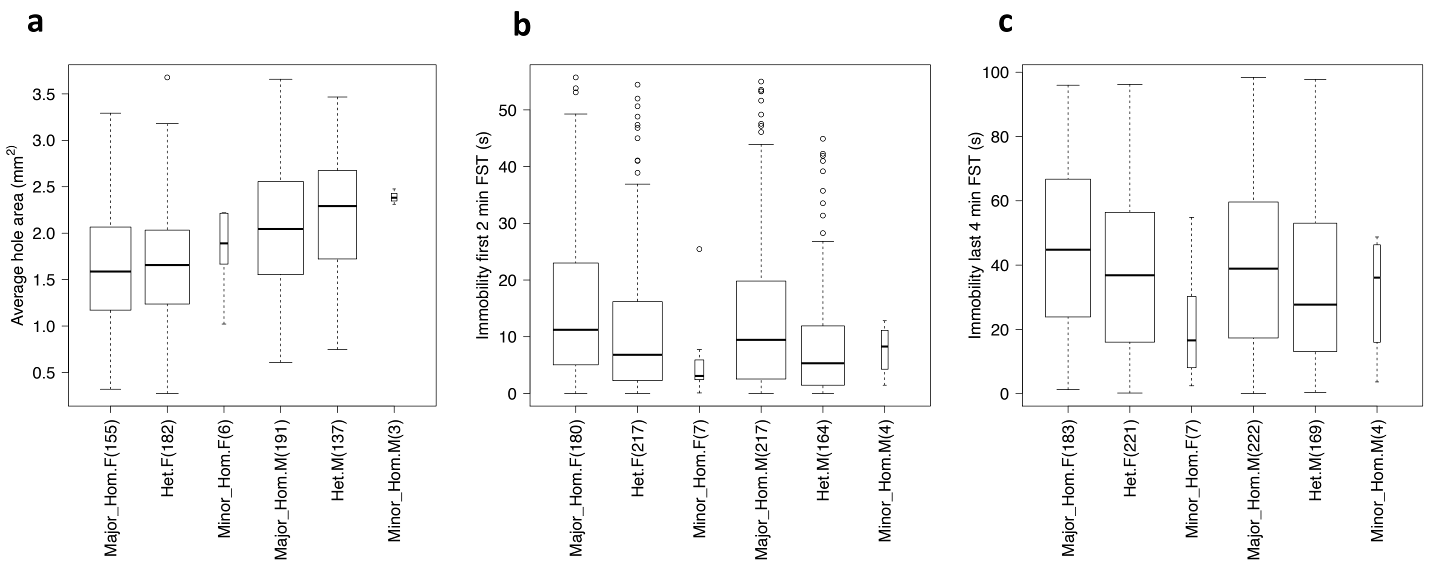


***Supplementary Figure 7*** *Phenotypes of the outbred CFW mice used in igeGWAS:* ***(a)*** *wound healing,* ***(b)*** *immobility during the first two minutes of the FST, and* ***(c)*** *immobility during the last four minutes of the FST. The data points (mice) are binned by genotype at the most significant variant at the Epha4 locus, namely chr1_77531018 for wound healing (as shown in Figure 4c) and chr1_76556104 for the FST measures (as shown in Figure 4a). In the x-axis label, Major and Minor refer to the major and minor alleles respectively, F and M to females and males, and the number in brackets is the number of mice in the bin. The boxes have different widths to reflect the different numbers of mice in each bin.*
