## Supplementary material for "Dissecting indirect genetic effects from peers in laboratory mice": locusZoom plots for all significant IGE loci: Bioch.Calcium_BWcorr_chr4_154482358_156984293_chr4_156222045.pdf

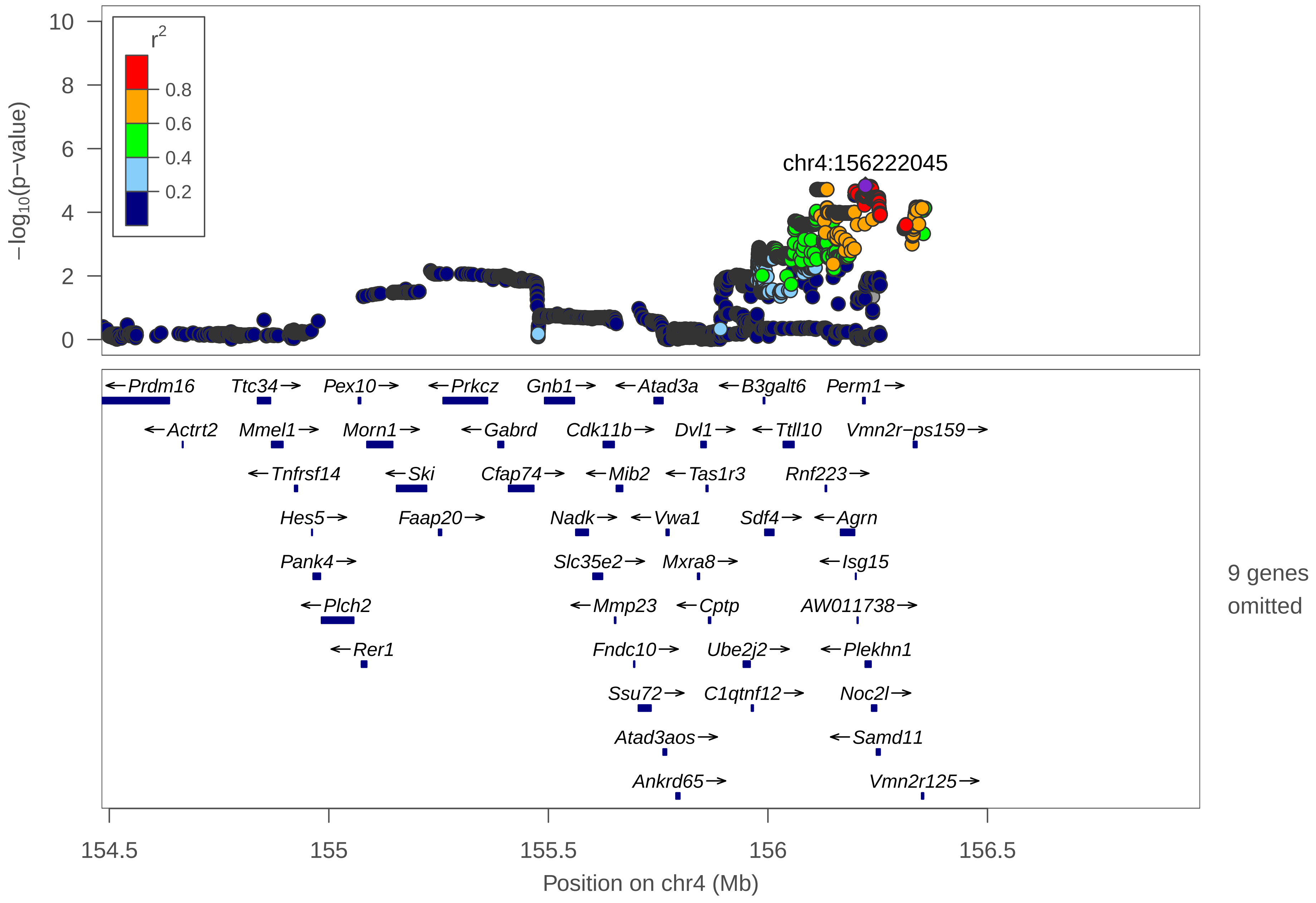

chr4:156222045
r2
0.8
0.6
0.4
0.2
−log10(p−value)
10
8
6
4
2
0
Prdm16
Ttc34
Pex10
Prkcz
Gnb1
Atad3a
B3galt6
Perm1
Actrt2
Mmel1
Morn1
Gabrd
Cdk11b
Dvl1
Ttll10
Vmn2r−ps159
Tnfrsf14
Ski
Cfap74
Mib2
Tas1r3
Rnf223
Hes5
Faap20
Nadk
Vwa1
Sdf4
Agrn
Pank4
Slc35e2
Mxra8
Isg15
Plch2
Mmp23
Cptp
AW011738
Rer1
Fndc10
Ube2j2
Plekhn1
Ssu72
C1qtnf12
Noc2l
Atad3aos
Samd11
Ankrd65
Vmn2r125
9 genes omitted
154.5
155
155.5
156
156.5
Position on chr4 (Mb)

| | |
|---|---|
| date: | Thu Apr 1 18:56:53 2021 |
| build: | mice |
| display range: | chr4:154482358−156984293 [154482358−156984293] |
| hilite range: | 0 − 0 [ 0 − 0 ] |
| reference SNP: | chr4:156222045 |
| number of SNPs plotted: | 7085 |
| min P−value: | 1.47E−5 [chr4:156222045] |
| max P−value: | 9.96E−1 [chr4:156218167] |
| omitted Genes: | Mrpl20, Ccnl2, Aurkaip1 |
| omitted Genes: | Mxra8os, Ints11, Pusl1 |
| omitted Genes: | Acap3, Tnfrsf4, Tnfrsf18 |

Make more plots at http://csg.sph.umich.edu/locuszoom/
