## Supplementary material for "Dissecting indirect genetic effects from peers in laboratory mice": locusZoom plots for all significant IGE loci: Bioch.Calcium_BWcorr_chr14_20575786_22075786_chr14_21512958.pdf

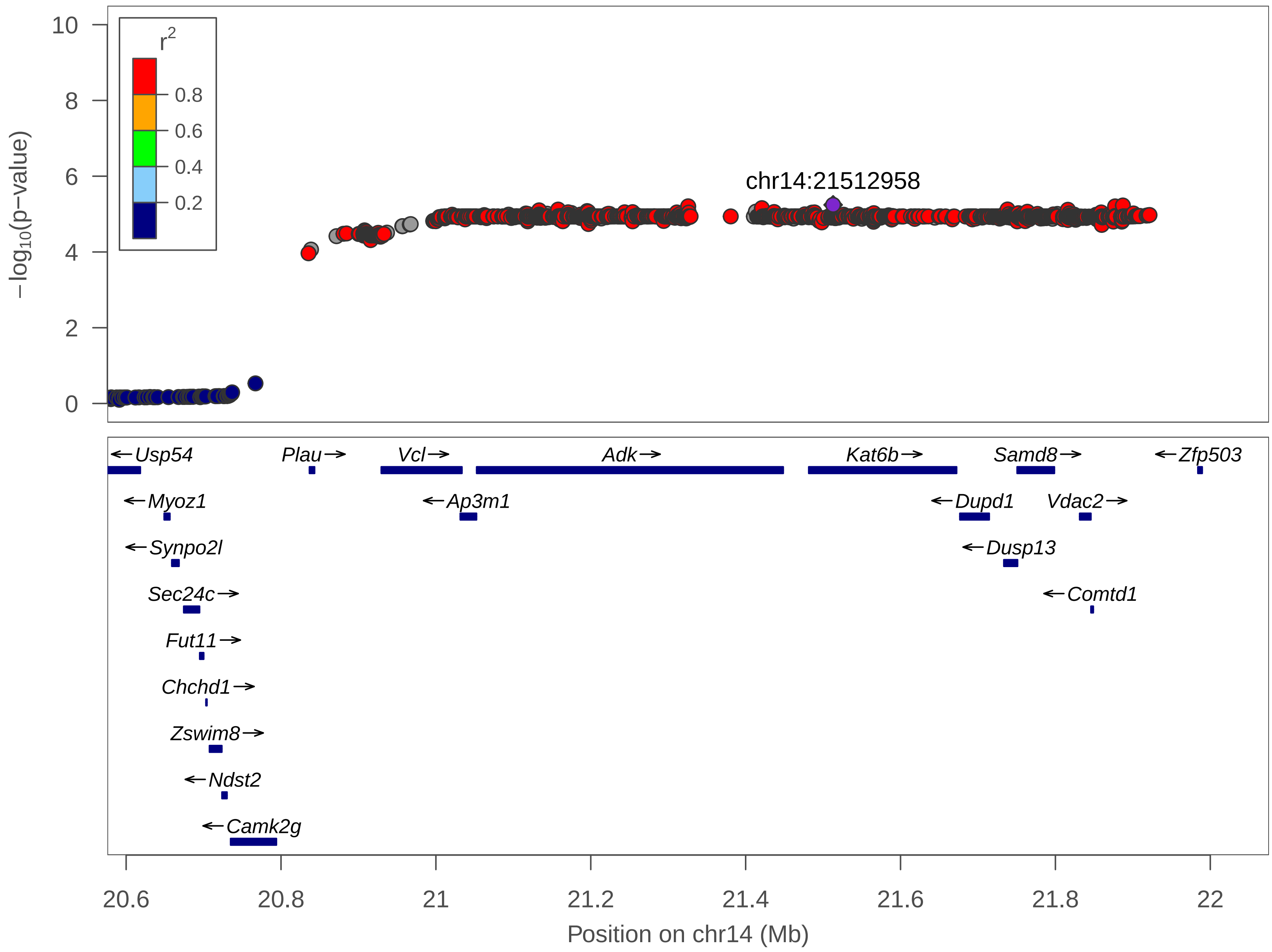

10
8
6
4
2
0
−log10(p−value)
r2
0.8
0.6
0.4
0.2
chr14:21512958
←Usp54
Plau→
Vcl→
Adk→
Kat6b→
Samd8→
←Zfp503
←Myoz1
←Ap3m1
←Dupd1
Vdac2→
←Synpo2l
←Dusp13
Sec24c→
←Comtd1
Fut11→
Chchd1→
Zswim8→
←Ndst2
←Camk2g
20.6
20.8
21
21.2
21.4
21.6
21.8
22
Position on chr14 (Mb)

| | |
|---|---|
| date: | Thu Apr 1 18:56:29 2021 |
| build: | mice |
| display range: | chr14:20575786−22075786 [20575786−22075786] |
| hilite range: | 0 − 0 [ 0 − 0 ] |
| reference SNP: | chr14:21512958 |
| number of SNPs plotted: | 2404 |
| min P−value: | 5.71E−6 [chr14:21512958] |
| max P−value: | 7.98E−1 [chr14:20591214] |

Make more plots at http://csg.sph.umich.edu/locuszoom/
