## Supplementary material for "Dissecting indirect genetic effects from peers in laboratory mice": locusZoom plots for all significant IGE loci: Bioch.Chloride_BWcorr_chr2_8774584_10274584_chr2_9531510.pdf

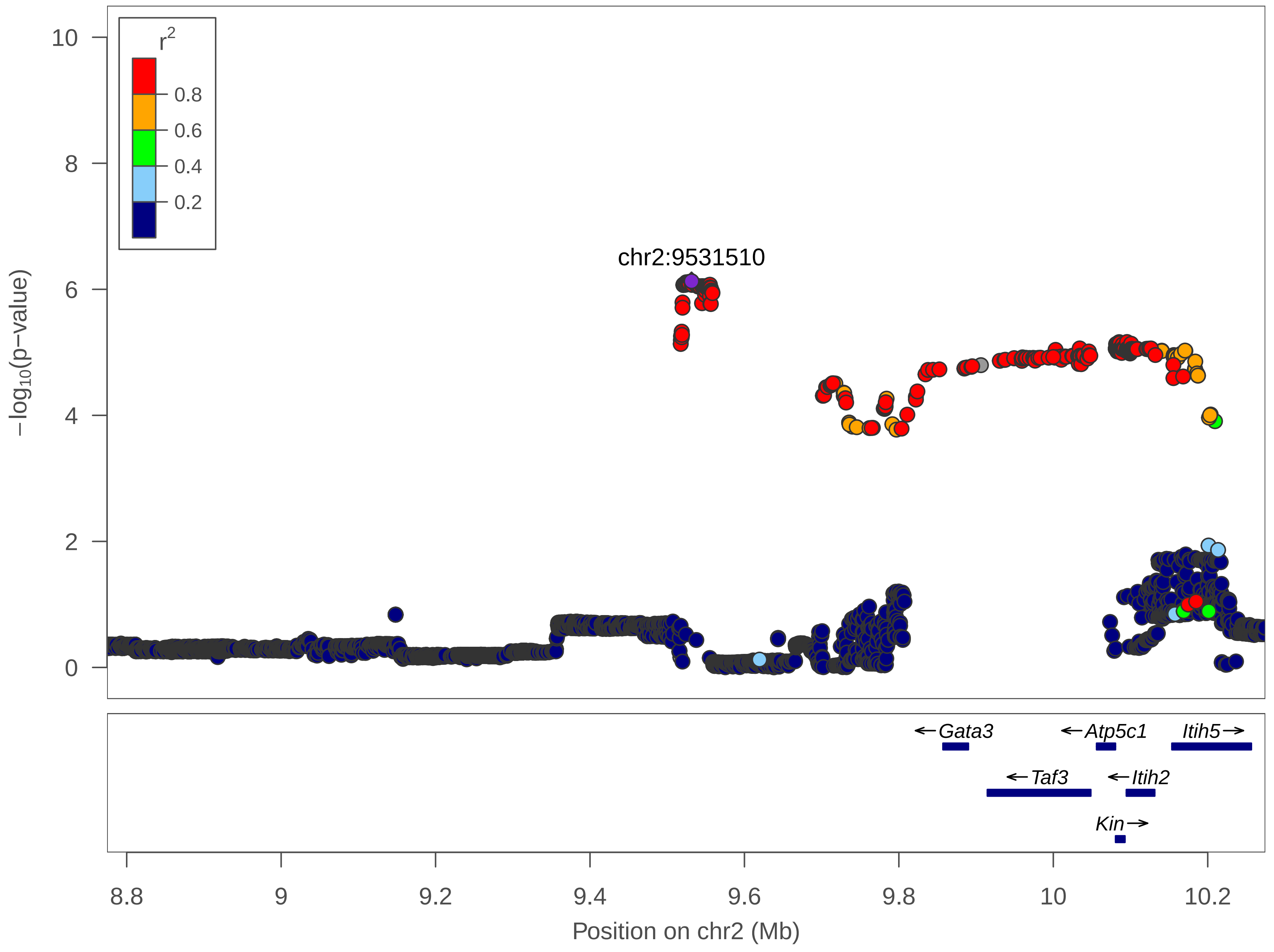

r2
0.8
0.6
0.4
0.2
10
8
6
4
2
0
−log10(p−value)
chr2:9531510
←Gata3
←Atp5c1
Itih5→
←Taf3
←Itih2
Kin→
8.8
9
9.2
9.4
9.6
9.8
10
10.2
Position on chr2 (Mb)

| | |
|---|---|
| date: | Thu Apr 1 18:57:19 2021 |
| build: | mice |
| display range: | chr2:8774584−10274584 [8774584−10274584] |
| hilite range: | 0 − 0 [ 0 − 0 ] |
| reference SNP: | chr2:9531510 |
| number of SNPs plotted: | 4578 |
| min P−value: | 7.45E−7 [chr2:9531510] |
| max P−value: | 9.96E−1 [chr2:9638150] |

Make more plots at http://csg.sph.umich.edu/locuszoom/
