## Supplementary material for "Dissecting indirect genetic effects from peers in laboratory mice": locusZoom plots for all significant IGE loci: BMC.Kurt_chr12_111918979_117155338_chr12_113713376.pdf

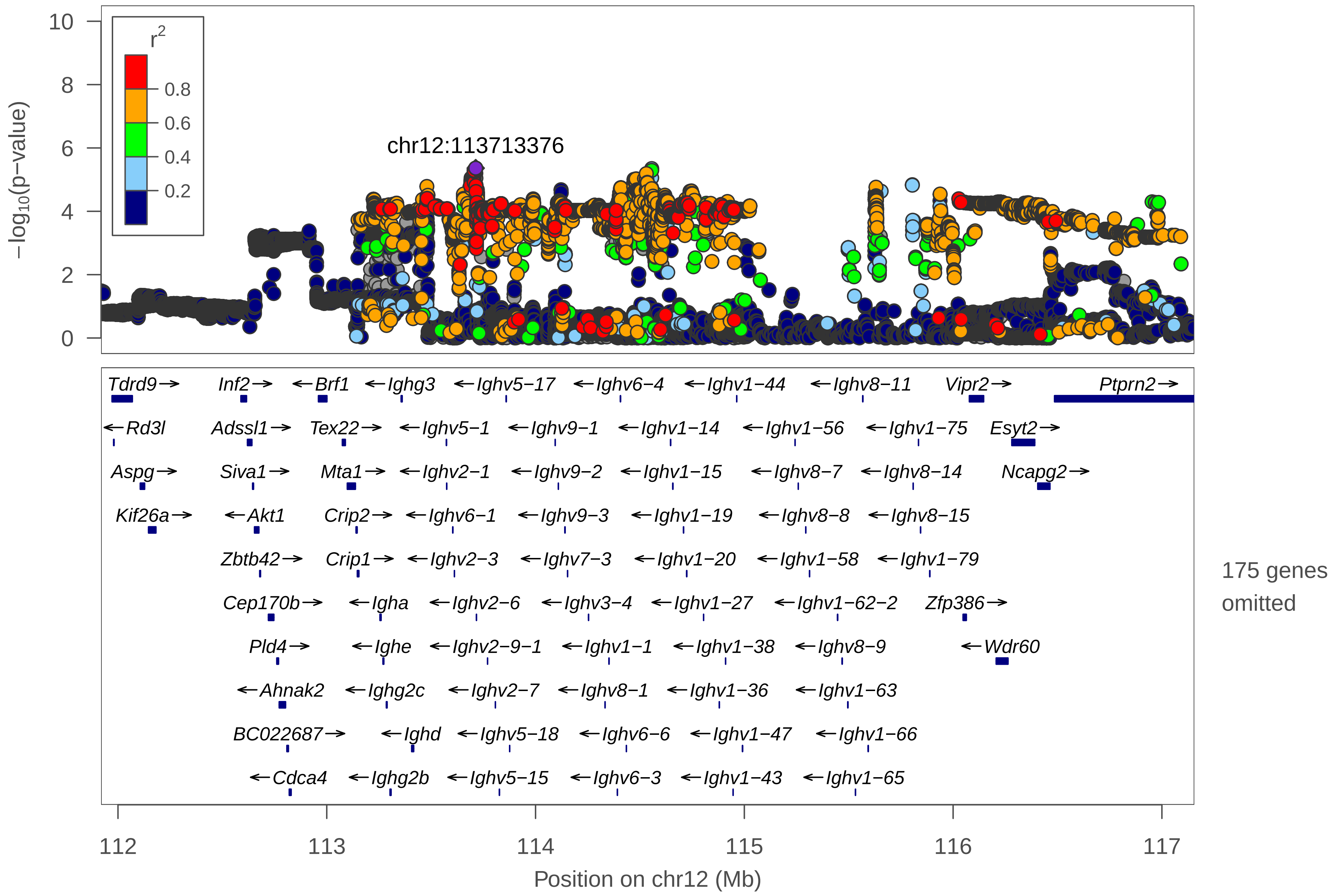

chr12:113713376
r2
0.8
0.6
0.4
0.2
−log10(p−value)
0
2
4
6
8
10
Tdrd9→
Inf2→
←Brf1
←Ighg3
←Ighv5−17
←Ighv6−4
←Ighv1−44
←Ighv8−11
Vipr2→
Ptprn2→
←Rd3l
Adssl1→
Tex22→
←Ighv5−1
←Ighv9−1
←Ighv1−14
←Ighv1−56
←Ighv1−75
Esyt2→
Aspg→
Siva1→
Mta1→
←Ighv2−1
←Ighv9−2
←Ighv1−15
←Ighv8−7
←Ighv8−14
Ncapg2→
Kif26a→
←Akt1
Crip2→
←Ighv6−1
←Ighv9−3
←Ighv1−19
←Ighv8−8
←Ighv8−15
Zbtb42→
Crip1→
←Ighv2−3
←Ighv7−3
←Ighv1−20
←Ighv1−58
←Ighv1−79
Cep170b→
←Igha
←Ighv2−6
←Ighv3−4
←Ighv1−27
←Ighv1−62−2
Zfp386→
Pld4→
←Ighe
←Ighv2−9−1
←Ighv1−1
←Ighv1−38
←Ighv8−9
←Wdr60
←Ahnak2
←Ighg2c
←Ighv2−7
←Ighv8−1
←Ighv1−36
←Ighv1−63
BC022687→
←Ighd
←Ighv5−18
←Ighv6−6
←Ighv1−47
←Ighv1−66
←Cdca4
←Ighg2b
←Ighv5−15
←Ighv6−3
←Ighv1−43
←Ighv1−65
175 genes omitted
112
113
114
115
116
117
Position on chr12 (Mb)

date: Fri Apr 2 10:54:20 2021

build: mice

display range: chr12:111918979−117155338 [111918979−117155338]

hilite range: 0 − 0 [ 0 − 0 ]

reference SNP: chr12:113713376

number of SNPs plotted: 49917

min P−value: 4.34E−6 [chr12:113713376]

max P−value: 10E−1 [chr12:116448505]

omitted Genes: Gpr132, Jag2, Nudt14

omitted Genes: Btbd6, Pacs2, Ighg1

omitted Genes: Ighm, Ighj4, Ighj3

omitted Genes: Ighj2, Ighj1, Ighd4−1

omitted Genes: Ighd3−2, Ighd5−6, Ighd2−8

omitted Genes: Ighd5−5, Ighd2−7, Ighd5−8

omitted Genes: Ighd5−4, Ighd2−6, Ighd5−7

omitted Genes: Ighd5−3, Ighd2−5, Ighd5−2

omitted Genes: Ighd2−4, Ighd6−2, Ighd2−3

omitted Genes: Ighd6−1, Ighd1−1, Adam6b

omitted Genes: Ighd3−1, Ighd5−1, Adam6a

omitted Genes: Ighv5−2, Ighv2−2, Ighv5−3

omitted Genes: Ighv5−4, Ighv5−5, Ighv5−6

omitted Genes: Ighv5−7, Ighv2−4, Ighv5−8

Make more plots at http://csg.sph.umich.edu/locuszoom/

omitted Genes: Ighv5−9, Ighv5−10, Ighv2−5
