## Supplementary material for "Dissecting indirect genetic effects from peers in laboratory mice": locusZoom plots for all significant IGE loci: BMC.osteoporosis_chr11_6269042_7769042_chr11_7033585.pdf

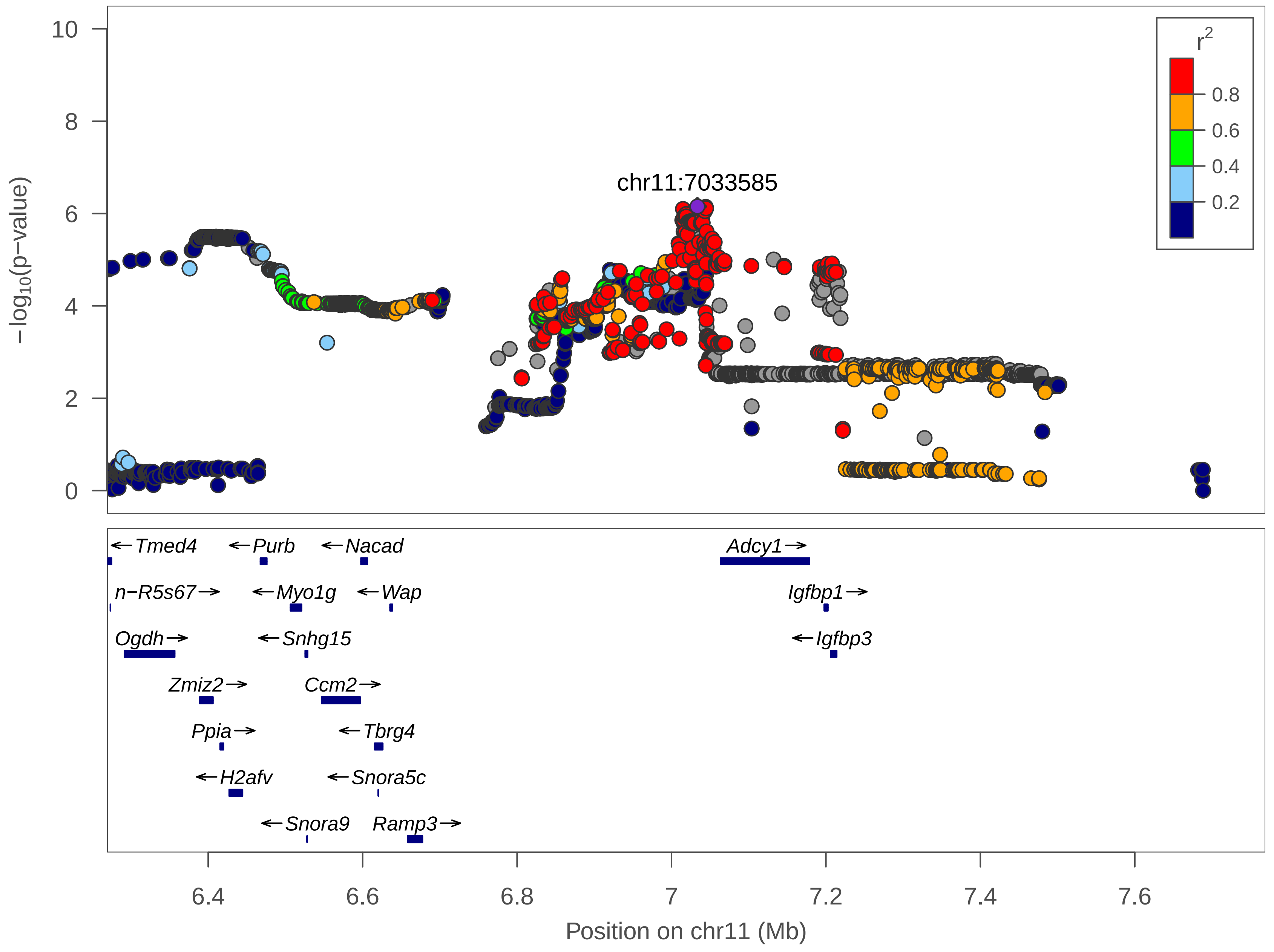

chr11:7033585
r2
0.8
0.6
0.4
0.2
−log10(p−value)
10
8
6
4
2
0
←Tmed4
←Purb
←Nacad
Adcy1→
n−R5s67→
←Myo1g
←Wap
Igfbp1→
Ogdh→
←Snhg15
←Igfbp3
Zmiz2→
Ccm2→
Ppia→
←Tbrg4
←H2afv
←Snora5c
←Snora9
Ramp3→
6.4
6.6
6.8
7
7.2
7.4
7.6
Position on chr11 (Mb)

| | |
|---|---|
| date: | Thu Apr 1 18:56:05 2021 |
| build: | mice |
| display range: | chr11:6269042−7769042 [6269042−7769042] |
| hilite range: | 0 − 0 [ 0 − 0 ] |
| reference SNP: | chr11:7033585 |
| number of SNPs plotted: | 3580 |
| min P−value: | 7.04E−7 [chr11:7033585] |
| max P−value: | 9.99E−1 [chr11:7688509] |

Make more plots at http://csg.sph.umich.edu/locuszoom/
