## Supplementary material for "Dissecting indirect genetic effects from peers in laboratory mice": locusZoom plots for all significant IGE loci: Cardio.ECG.Tpeak_Tend_BWcorr_chr19_17107339_21380478_chr19_18647680.pdf

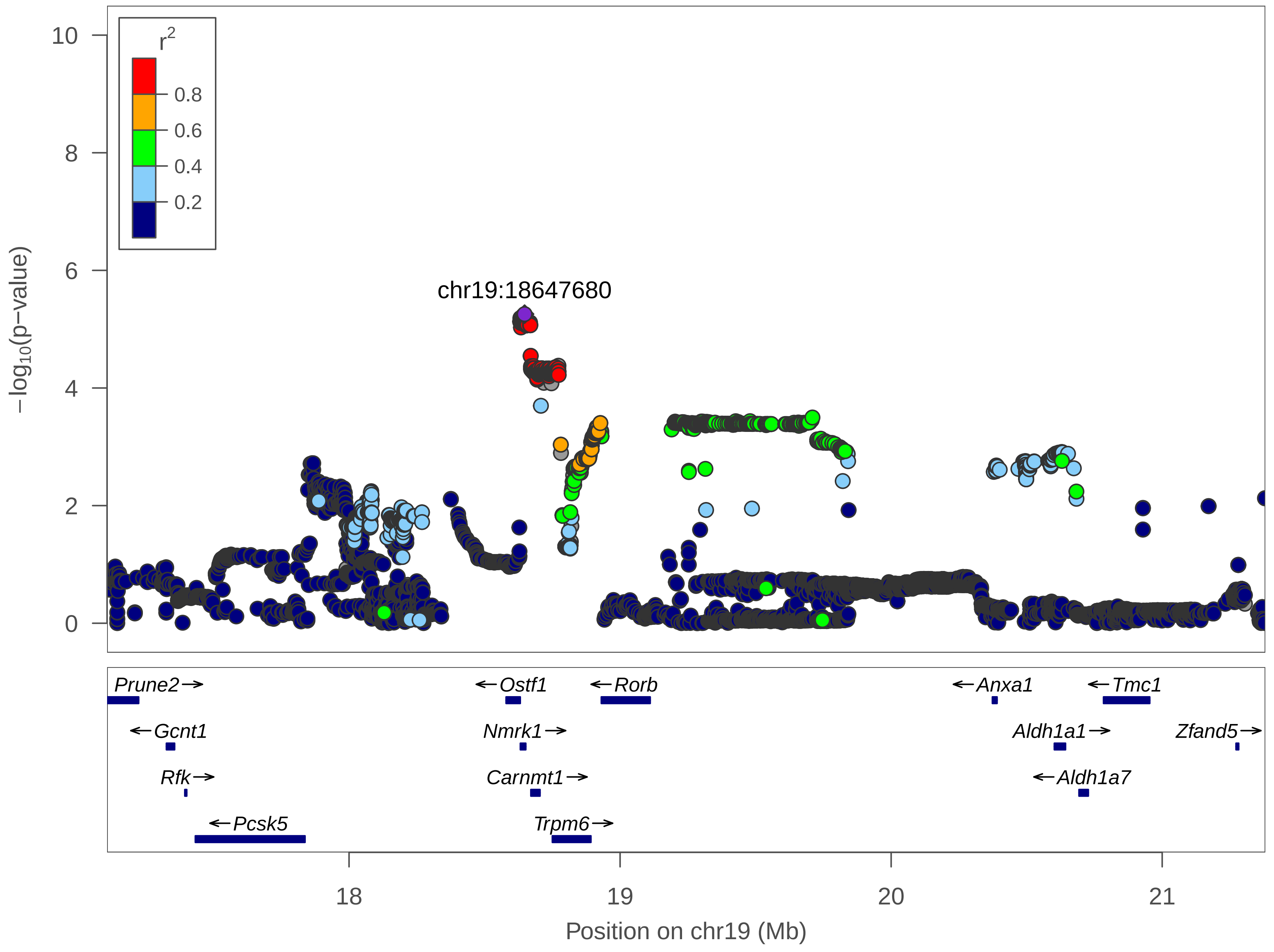

r2
0.8
0.6
0.4
0.2
10
8
6
4
2
0
−log10(p−value)
chr19:18647680
Prune2→
←Gcnt1
Rfk→
←Pcsk5
←Ostf1
Nmrk1→
Carnmt1→
Trpm6→
←Rorb
←Anxa1
←Tmc1
Aldh1a1→
←Aldh1a7
Zfand5→
18
19
20
21
Position on chr19 (Mb)

| | |
|---|---|
| date: | Thu Apr 1 18:58:27 2021 |
| build: | mice |
| display range: | chr19:17107339−21380478 [17107339−21380478] |
| hilite range: | 0 − 0 [ 0 − 0 ] |
| reference SNP: | chr19:18647680 |
| number of SNPs plotted: | 14523 |
| min P−value: | 5.55E−6 [chr19:18647680] |
| max P−value: | 10E−1 [chr19:19284658] |

Make more plots at http://csg.sph.umich.edu/locuszoom/
