## Supplementary material for "Dissecting indirect genetic effects from peers in laboratory mice": locusZoom plots for all significant IGE loci: FACS.CD3posCD4pos_chr12_43053888_47044310_chr12_44996254.pdf

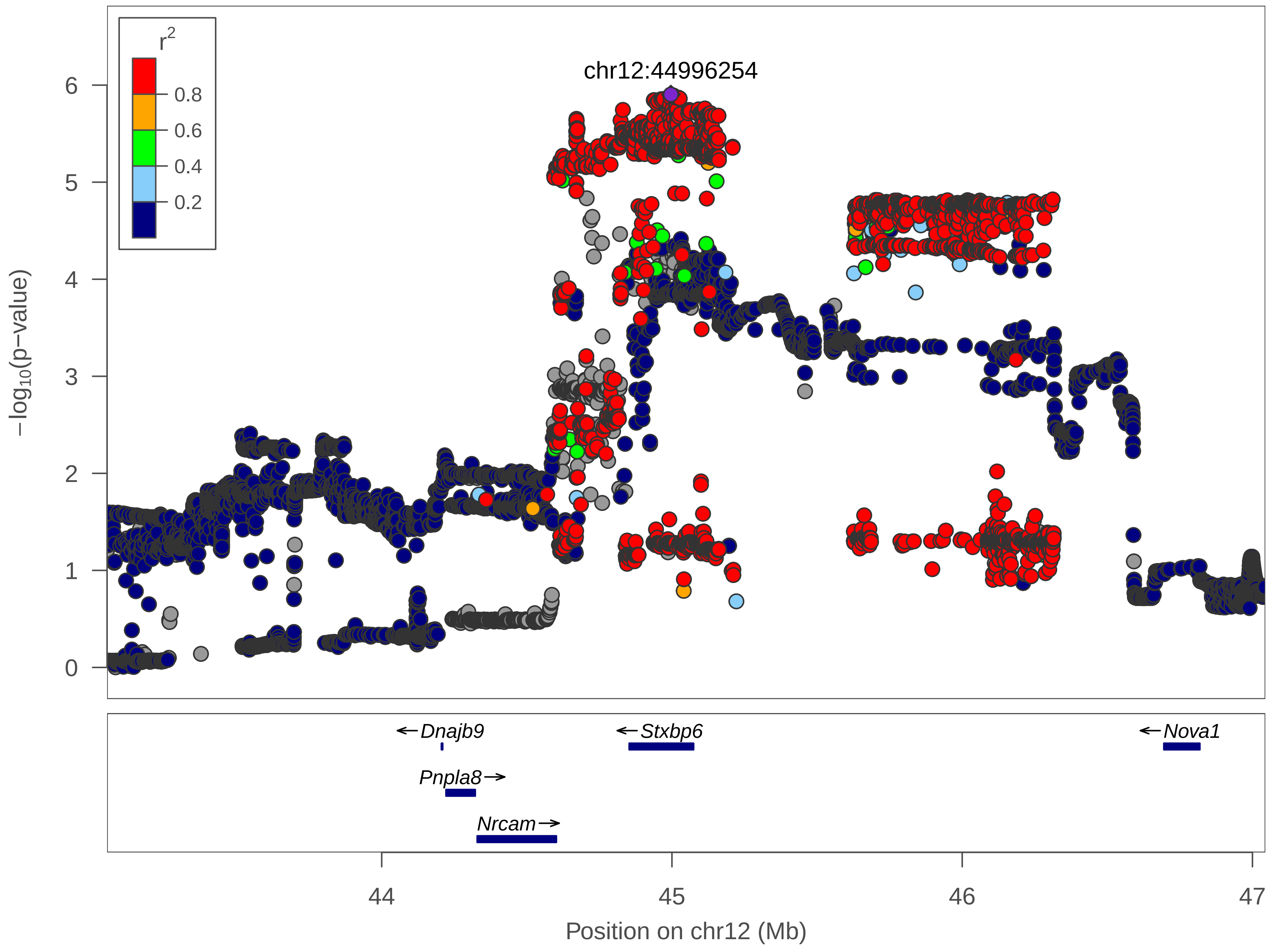

r2
0.8
0.6
0.4
0.2
chr12:44996254
6
5
4
3
2
1
0
−log10(p−value)
←Dnajb9
←Stxbp6
←Nova1
Pnpla8→
Nrcam→
44
45
46
47
Position on chr12 (Mb)

| | |
|---|---|
| date: | Fri Apr 2 15:53:40 2021 |
| build: | mice |
| display range: | chr12:43053888−47044310 [43053888−47044310] |
| hilite range: | 0 − 0 [ 0 − 0 ] |
| reference SNP: | chr12:44996254 |
| number of SNPs plotted: | 14460 |
| min P−value: | 1.25E−6 [chr12:44996254] |
| max P−value: | 9.97E−1 [chr12:43083124] |

Make more plots at http://csg.sph.umich.edu/locuszoom/
