## Supplementary material for "Dissecting indirect genetic effects from peers in laboratory mice": locusZoom plots for all significant IGE loci: Haem.abs_neuts_chr1_34981852_36481852_chr1_35729352.pdf

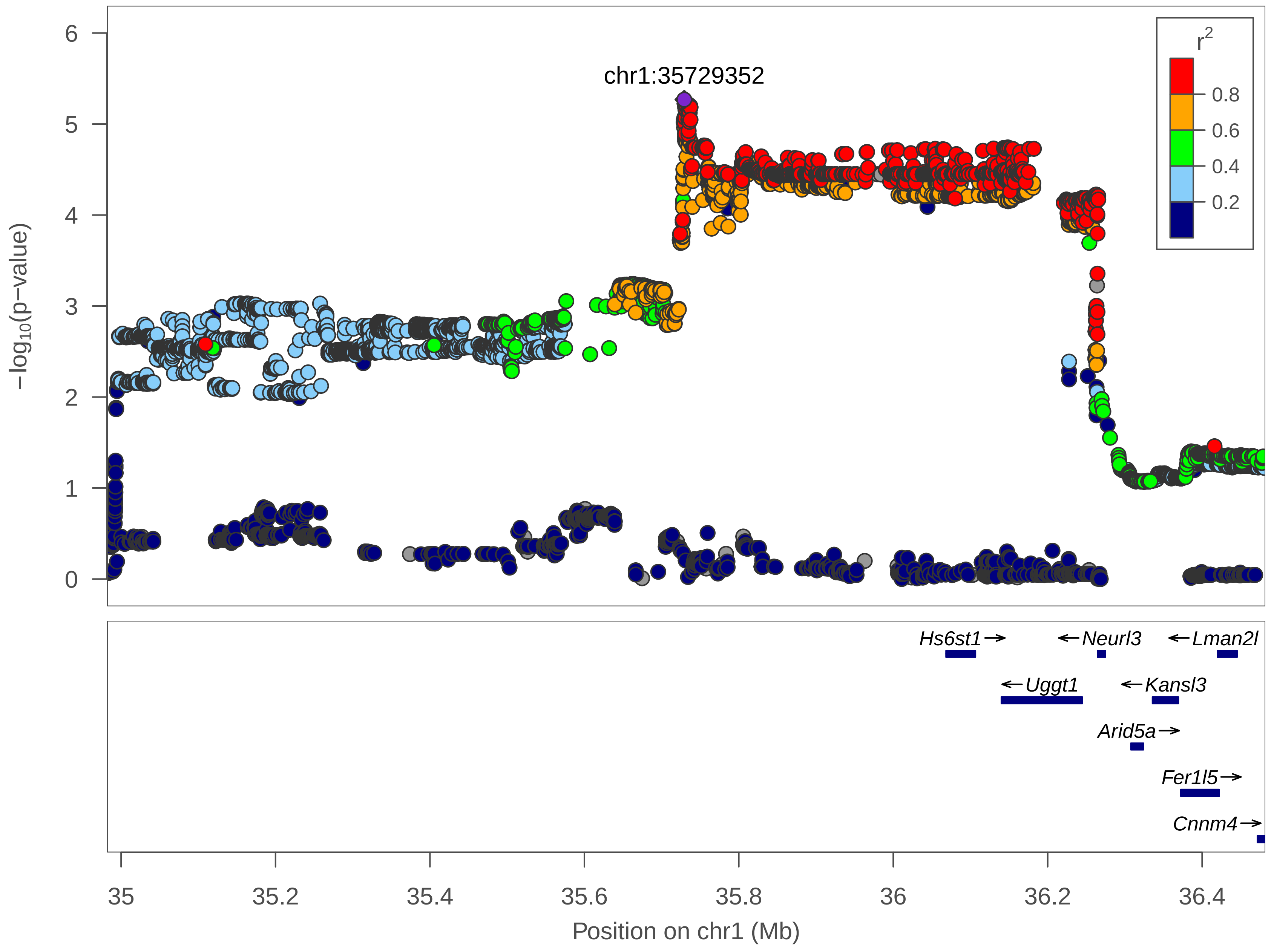

chr1:35729352
$r^2$
0.8
0.6
0.4
0.2
$-\log_{10}$(p-value)
0
1
2
3
4
5
6
Hs6st1
Neurl3
Lman2l
Uggt1
Kansl3
Arid5a
Fer1l5
Cnnm4
35
35.2
35.4
35.6
35.8
36
36.2
36.4
Position on chr1 (Mb)

| | |
|---|---|
| date: | Fri Apr 2 15:54:00 2021 |
| build: | mice |
| display range: | chr1:34981852−36481852 [34981852−36481852] |
| hilite range: | 0 − 0 [ 0 − 0 ] |
| reference SNP: | chr1:35729352 |
| number of SNPs plotted: | 7743 |
| min P−value: | 5.39E−6 [chr1:35729352] |
| max P−value: | 10E−1 [chr1:36010960] |

Make more plots at http://csg.sph.umich.edu/locuszoom/
