## Supplementary material for "Dissecting indirect genetic effects from peers in laboratory mice": locusZoom plots for all significant IGE loci: Hypoxia.f_NR_BWcorr_chr12_112992891_114492891_chr12_113742891.pdf

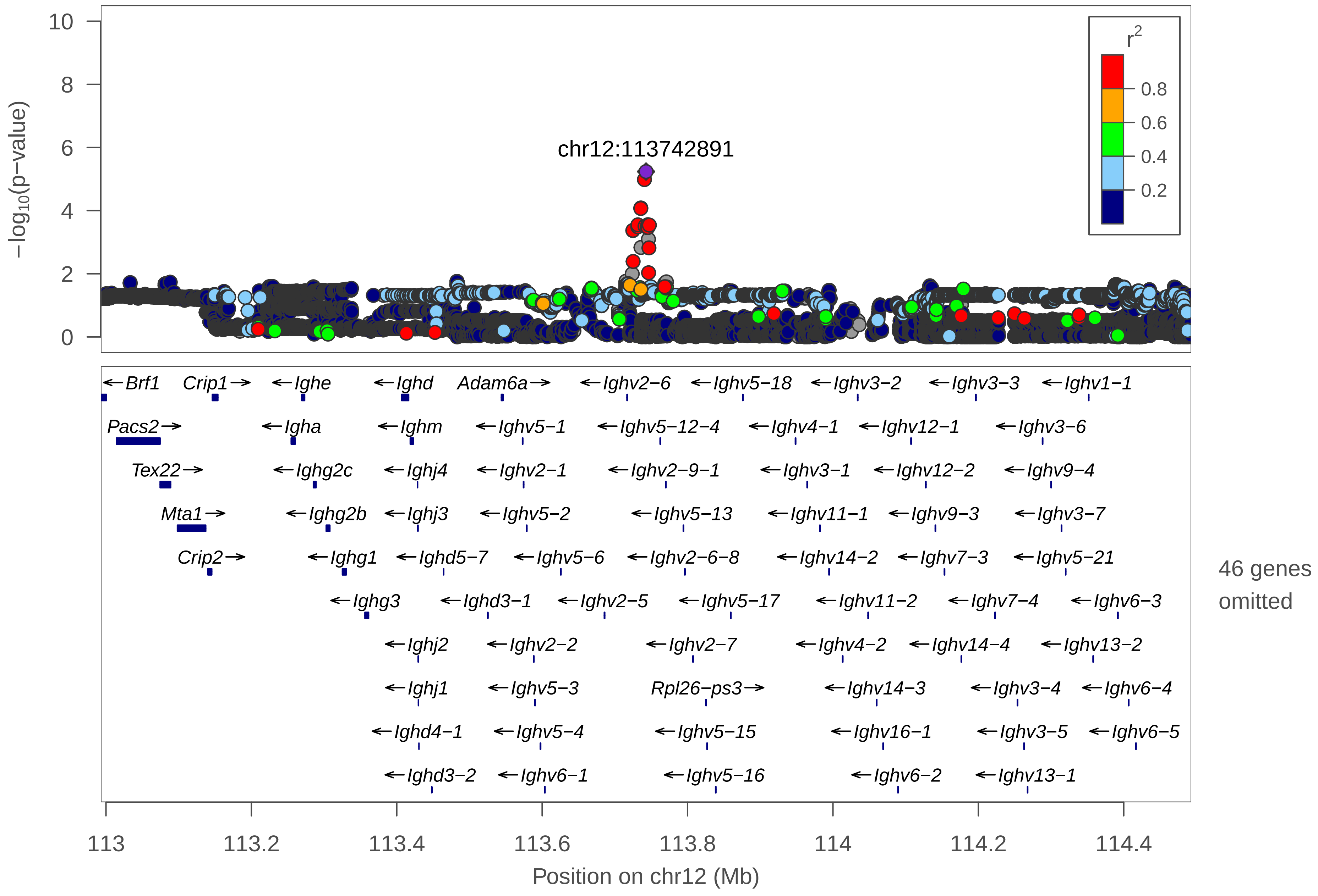

chr12:113742891
r2
0.8
0.6
0.4
0.2
−log10(p−value)
10
8
6
4
2
0
←Brf1
Crip1→
←Ighe
←Ighd
Adam6a→
←Ighv2−6
←Ighv5−18
←Ighv3−2
←Ighv3−3
←Ighv1−1
Pacs2→
←Igha
←Ighm
←Ighv5−1
←Ighv5−12−4
←Ighv4−1
←Ighv12−1
←Ighv3−6
Tex22→
←Ighg2c
←Ighj4
←Ighv2−1
←Ighv2−9−1
←Ighv3−1
←Ighv12−2
←Ighv9−4
Mta1→
←Ighg2b
←Ighj3
←Ighv5−2
←Ighv5−13
←Ighv11−1
←Ighv9−3
←Ighv3−7
Crip2→
←Ighg1
←Ighd5−7
←Ighv5−6
←Ighv2−6−8
←Ighv14−2
←Ighv7−3
←Ighv5−21
←Ighg3
←Ighd3−1
←Ighv2−5
←Ighv5−17
←Ighv11−2
←Ighv7−4
←Ighv6−3
←Ighj2
←Ighv2−2
←Ighv2−7
←Ighv4−2
←Ighv14−4
←Ighv13−2
←Ighj1
←Ighv5−3
Rpl26−ps3→
←Ighv14−3
←Ighv3−4
←Ighv6−4
←Ighd4−1
←Ighv5−4
←Ighv5−15
←Ighv16−1
←Ighv3−5
←Ighv6−5
←Ighd3−2
←Ighv6−1
←Ighv5−16
←Ighv6−2
←Ighv13−1
46 genes omitted
113
113.2
113.4
113.6
113.8
114
114.2
114.4
Position on chr12 (Mb)

date: Thu Apr 1 18:57:50 2021

build: mice

display range: chr12:112992891−114492891 [112992891−114492891]

hilite range: 0 − 0 [ 0 − 0 ]

reference SNP: chr12:113742891

number of SNPs plotted: 21165

min P−value: 5.79E−6 [chr12:113742891]

max P−value: 10E−1 [chr12:114421905]

omitted Genes: Ighd5−6, Ighd2−8, Ighd5−5

omitted Genes: Ighd2−7, Ighd5−8, Ighd5−4

omitted Genes: Ighd2−6, Ighd5−3, Ighd2−5

omitted Genes: Ighd5−2, Ighd2−4, Ighd6−2

omitted Genes: Ighd2−3, Ighd6−1, Ighd1−1

omitted Genes: Adam6b, Ighd5−1, Ighv2−3

omitted Genes: Ighv5−5, Ighv5−7, Ighv2−4

omitted Genes: Ighv5−8, Ighv5−9, Ighv5−10

omitted Genes: Ighv5−11, Ighv5−12, Ighv5−9−1

omitted Genes: Ighv2−8, Ighv2−9, Ighv5−19

omitted Genes: Ighv7−1, Ighv7−2, Ighv14−1

omitted Genes: Ighv9−1, Ighv9−2, Ighv15−1

omitted Genes: Ighv3−8, Ighv8−1, Ighv12−3

omitted Genes: Ighv6−6, Ighv6−7, Ighv8−2

Make more plots at http://csg.sph.umich.edu/locuszoom/

omitted Genes: Ighv1−2, Ighv10−1, Ighv1−3
