## Supplementary material for "Dissecting indirect genetic effects from peers in laboratory mice": locusZoom plots for all significant IGE loci: Neuro.DCX_BWcorr_chr12_2426906_3926906_chr12_3176906.pdf

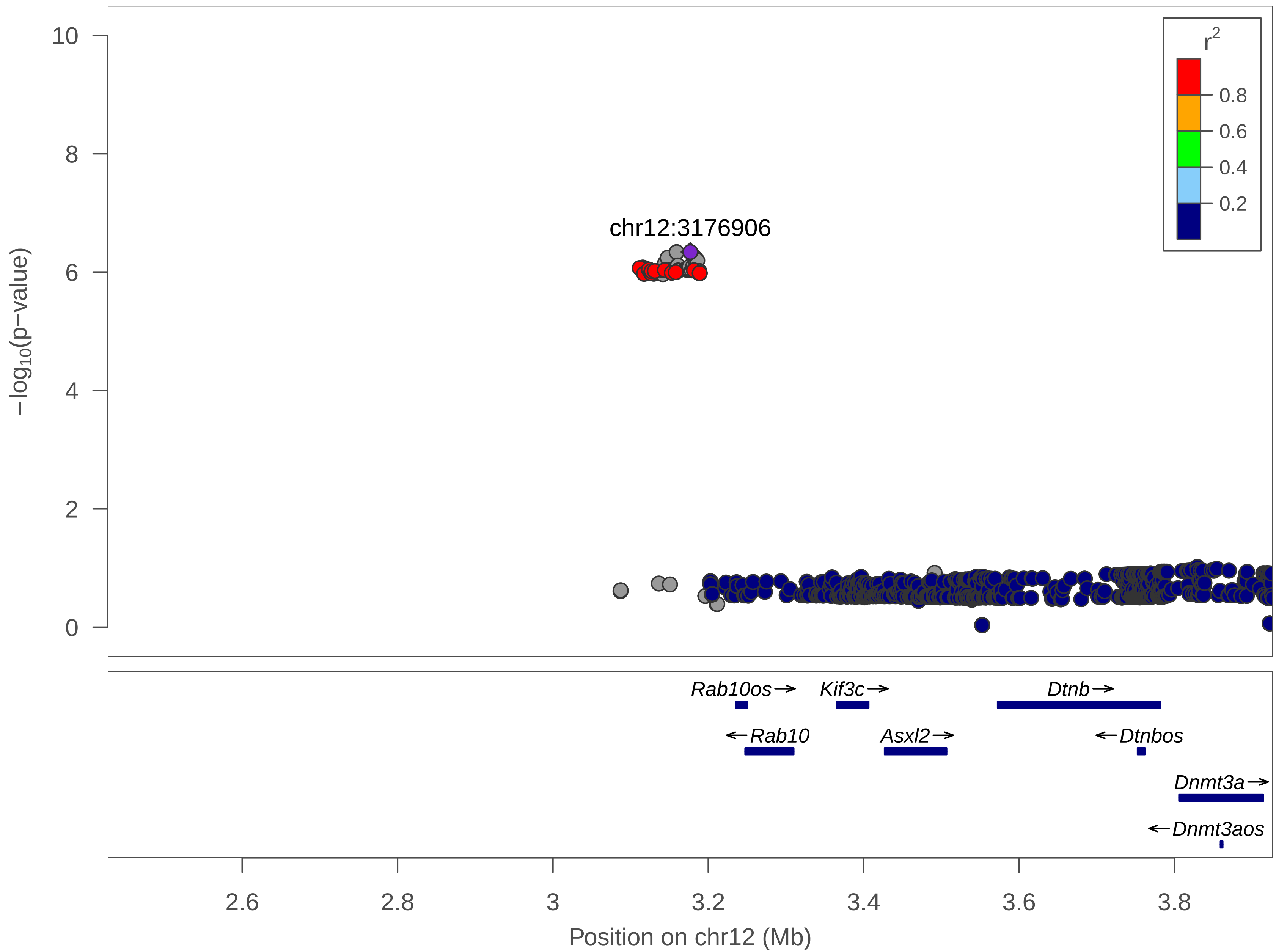

chr12:3176906
r2
0.8
0.6
0.4
0.2
10
8
6
4
2
0
−log10(p−value)
Rab10os→
Kif3c→
Dtnb→
←Rab10
Asxl2→
←Dtnbos
Dnmt3a→
←Dnmt3aos
2.6
2.8
3
3.2
3.4
3.6
3.8
Position on chr12 (Mb)

| | |
|---|---|
| date: | Thu Apr 1 19:03:48 2021 |
| build: | mice |
| display range: | chr12:2426906−3926906 [2426906−3926906] |
| hilite range: | 0 − 0 [ 0 − 0 ] |
| reference SNP: | chr12:3176906 |
| number of SNPs plotted: | 703 |
| min P−value: | 4.58E−7 [chr12:3176906] |
| max P−value: | 9.26E−1 [chr12:3552581] |

Make more plots at http://csg.sph.umich.edu/locuszoom/
