## Supplementary material for "Dissecting indirect genetic effects from peers in laboratory mice": locusZoom plots for all significant IGE loci: Neuro.Ki67_BWcorr_chr1_69394500_72089531_chr1_71318343.pdf

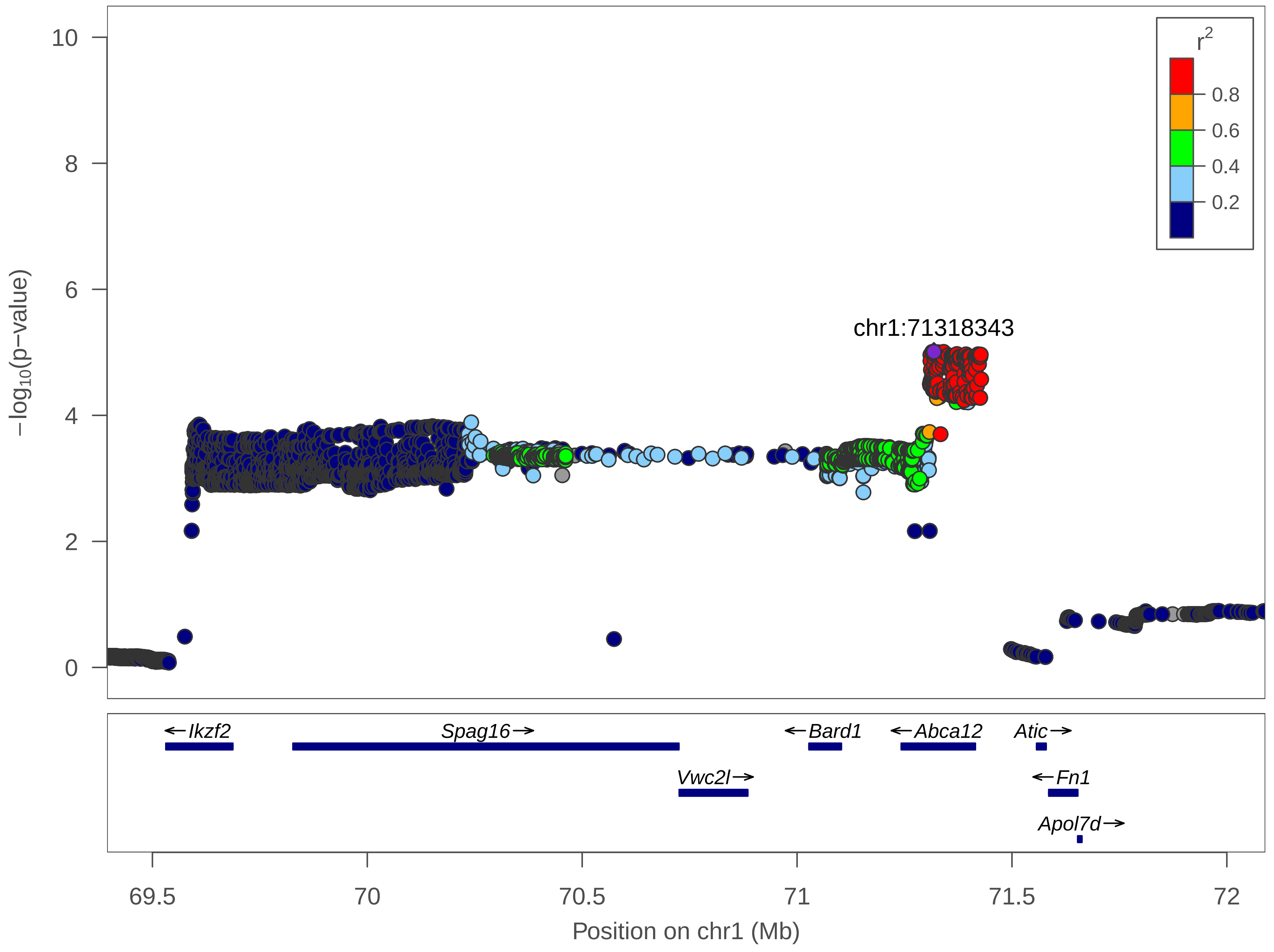

r^2
0.8
0.6
0.4
0.2
10
8
6
4
2
0
−log10(p−value)
chr1:71318343
←Ikzf2
Spag16→
←Bard1
←Abca12
Atic→
Vwc2l→
←Fn1
Apol7d→
69.5
70
70.5
71
71.5
72
Position on chr1 (Mb)

| | |
|---|---|
| date: | Thu Apr 1 19:03:28 2021 |
| build: | mice |
| display range: | chr1:69394500−72089531 [69394500−72089531] |
| hilite range: | 0 − 0 [ 0 − 0 ] |
| reference SNP: | chr1:71318343 |
| number of SNPs plotted: | 5523 |
| min P−value: | 9.81E−6 [chr1:71318343] |
| max P−value: | 8.39E−1 [chr1:69538277] |

Make more plots at http://csg.sph.umich.edu/locuszoom/
