## Supplementary material for "Dissecting indirect genetic effects from peers in laboratory mice": locusZoom plots for all significant IGE loci: Neuro.Ki67_BWcorr_chr10_17537371_21564231_chr10_19064386.pdf

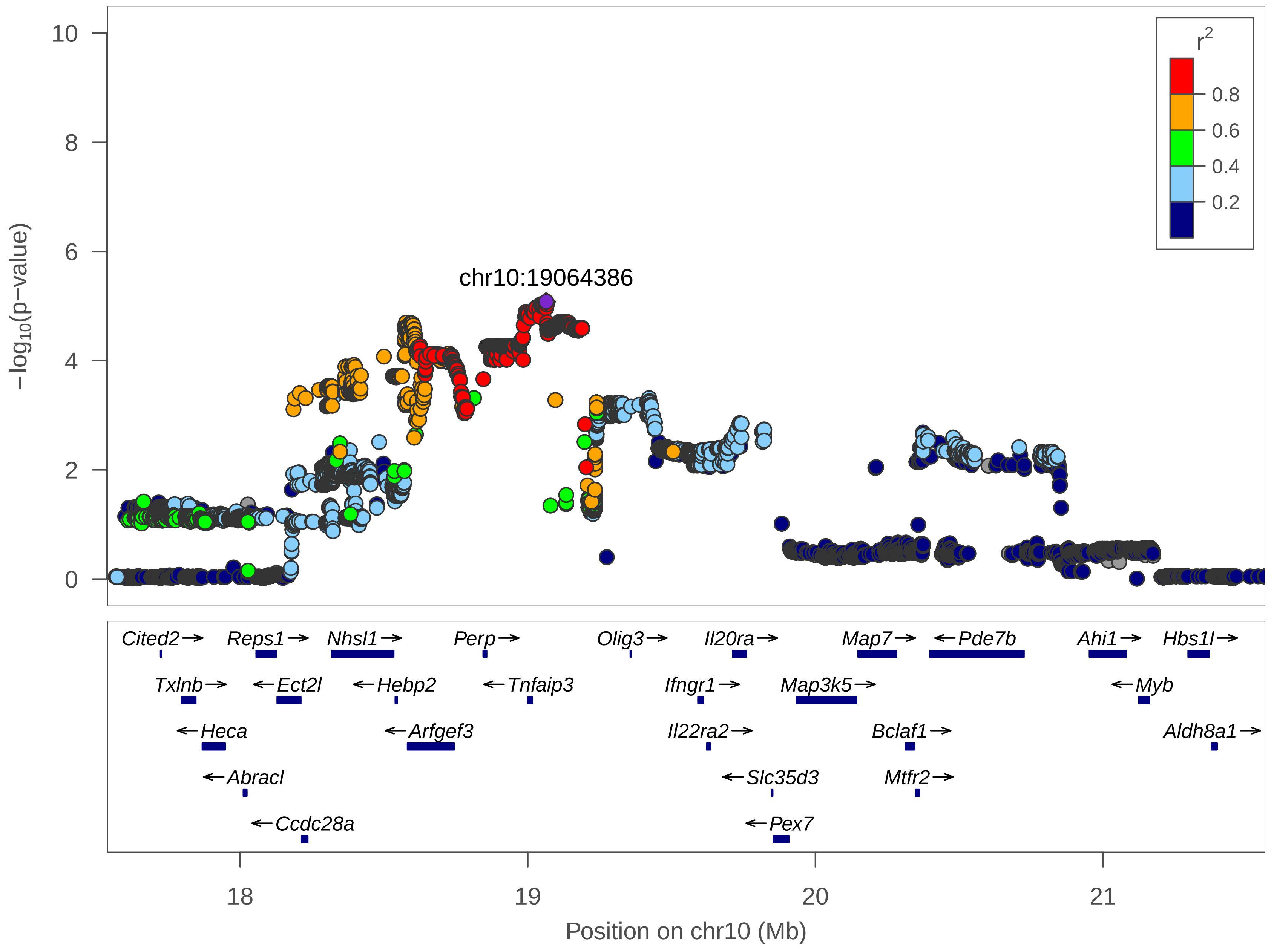

chr10:19064386
r2
0.8
0.6
0.4
0.2
−log10(p−value)
0
2
4
6
8
10
Cited2→
Reps1→
Nhsl1→
Perp→
Olig3→
Il20ra→
Map7→
←Pde7b
Ahi1→
Hbs1l→
Txlnb→
←Ect2l
←Hebp2
←Tnfaip3
Ifngr1→
Map3k5→
←Myb
←Heca
←Arfgef3
Il22ra2→
Bclaf1→
Aldh8a1→
←Abracl
←Slc35d3
Mtfr2→
←Ccdc28a
←Pex7
18
19
20
21
Position on chr10 (Mb)

| | |
|---|---|
| date: | Thu Apr 1 19:03:04 2021 |
| build: | mice |
| display range: | chr10:17537371−21564231 [17537371−21564231] |
| hilite range: | 0 − 0 [ 0 − 0 ] |
| reference SNP: | chr10:19064386 |
| number of SNPs plotted: | 10993 |
| min P−value: | 8.31E−6 [chr10:19064386] |
| max P−value: | 9.85E−1 [chr10:21117644] |

Make more plots at http://csg.sph.umich.edu/locuszoom/
