## Supplementary material for "Dissecting indirect genetic effects from peers in laboratory mice": locusZoom plots for all significant IGE loci: PAS.First5_chr10_2378239_3878239_chr10_3128410.pdf

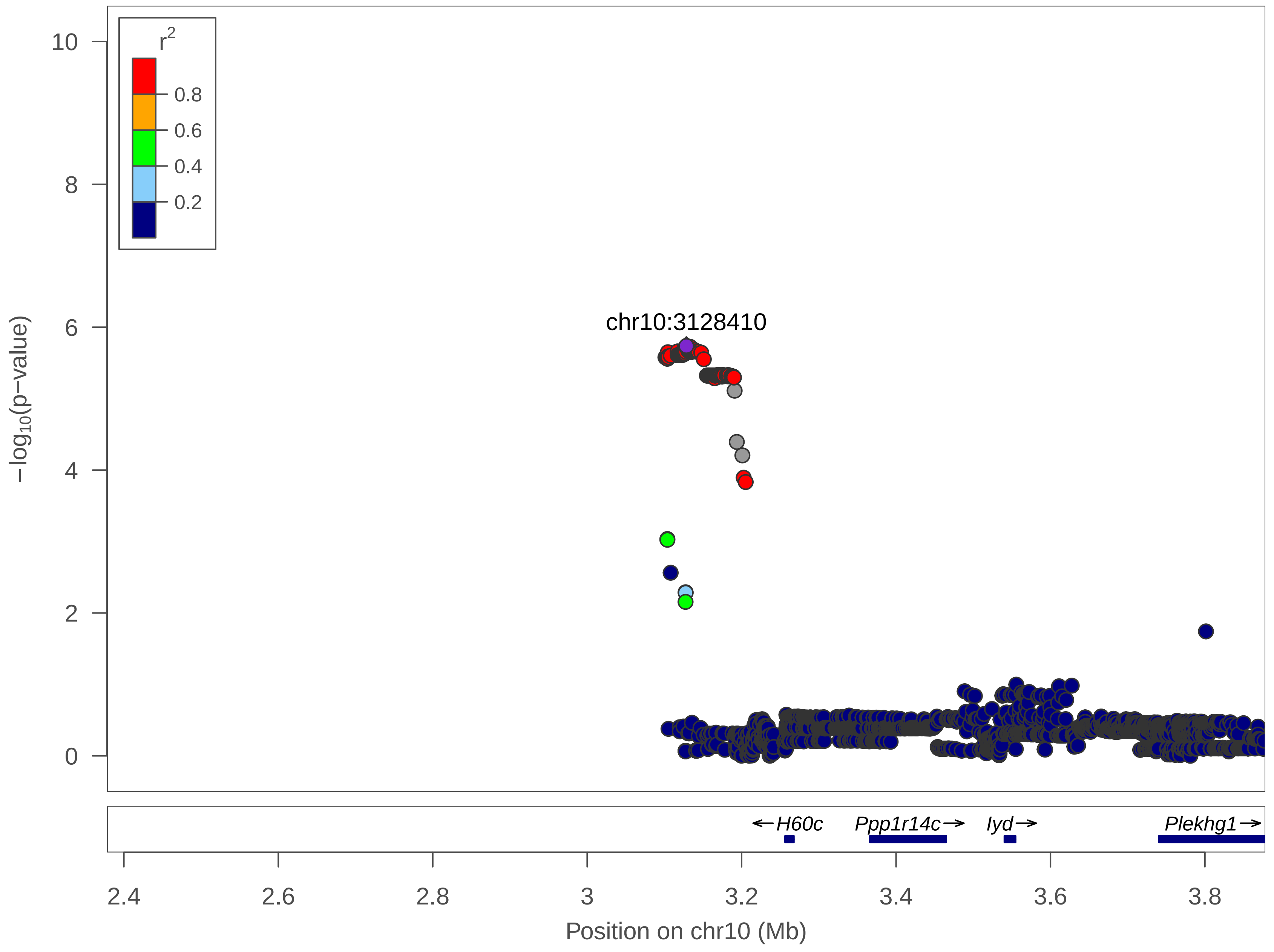
r2
0.8
0.6
0.4
0.2
10
8
6
4
2
0
−log10(p−value)
chr10:3128410
H60c
Ppp1r14c
Iyd
Plekhg1
2.4
2.6
2.8
3
3.2
3.4
3.6
3.8
Position on chr10 (Mb)

| | |
|---|---|
| date: | Thu Apr 1 18:55:42 2021 |
| build: | mice |
| display range: | chr10:2378239−3878239 [2378239−3878239] |
| hilite range: | 0 − 0 [ 0 − 0 ] |
| reference SNP: | chr10:3128410 |
| number of SNPs plotted: | 3074 |
| min P−value: | 1.83E−6 [chr10:3128410] |
| max P−value: | 9.97E−1 [chr10:3781106] |

Make more plots at http://csg.sph.umich.edu/locuszoom/
