## Supplementary material for "Dissecting indirect genetic effects from peers in laboratory mice": locusZoom plots for all significant IGE loci: PST.Immobility.First2min_chr1_75762651_78078619_chr1_76556104.pdf

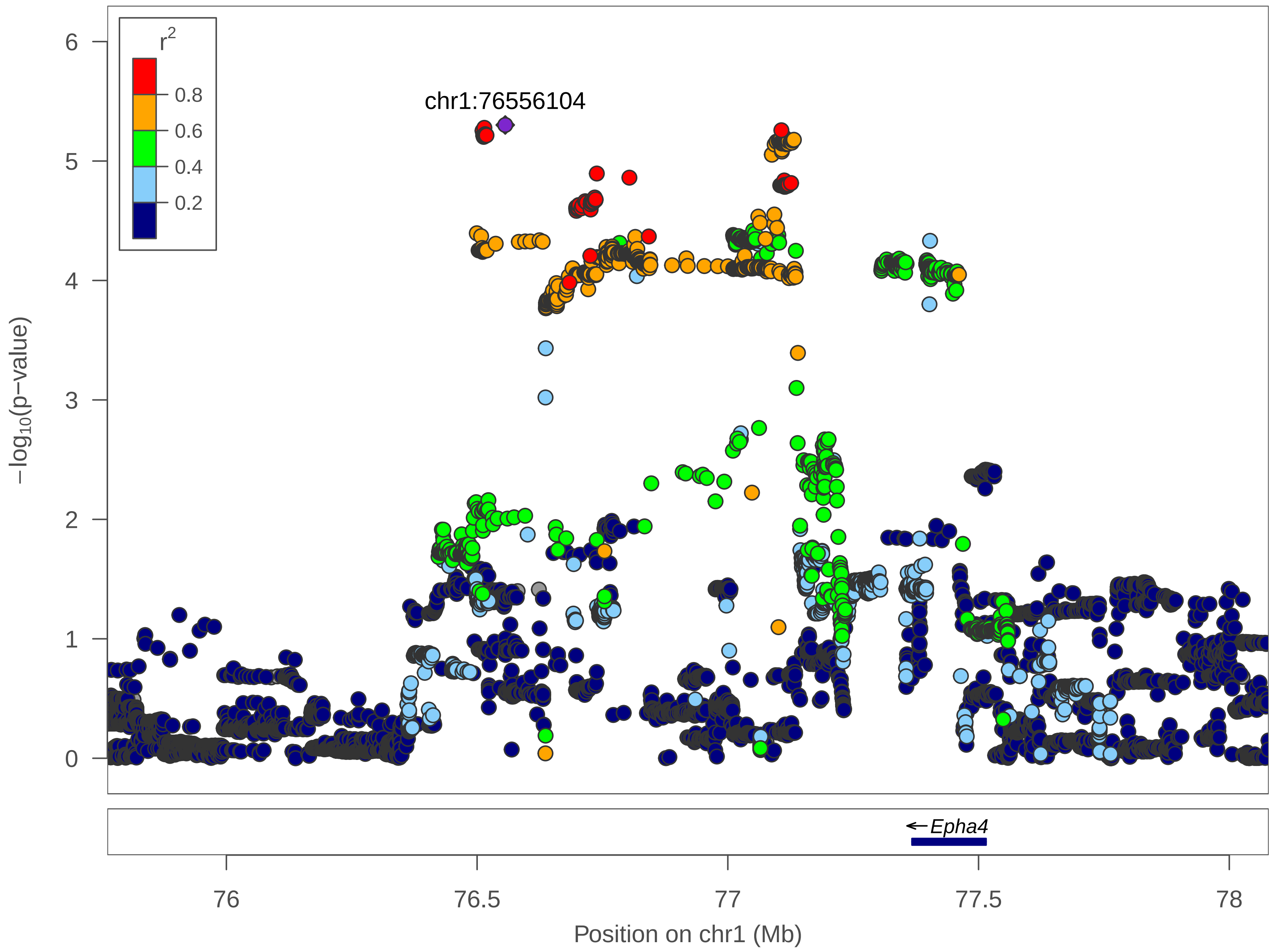
r2
0.8
0.6
0.4
0.2
chr1:76556104
6
5
4
3
2
1
0
−log10(p−value)
Epha4
76
76.5
77
77.5
78
Position on chr1 (Mb)

| | |
|---|---|
| date: | Fri Apr 2 15:56:51 2021 |
| build: | mice |
| display range: | chr1:75762651−78078619 [75762651−78078619] |
| hilite range: | 0 − 0 [ 0 − 0 ] |
| reference SNP: | chr1:76556104 |
| number of SNPs plotted: | 10189 |
| min P−value: | 4.99E−6 [chr1:76556104] |
| max P−value: | 10E−1 [chr1:77763171] |

Make more plots at http://csg.sph.umich.edu/locuszoom/
