## Supplementary material for "Dissecting indirect genetic effects from peers in laboratory mice": locusZoom plots for all significant IGE loci: PST.Immobility.Last4min_chr9_86736657_90358438_chr9_88502417.pdf

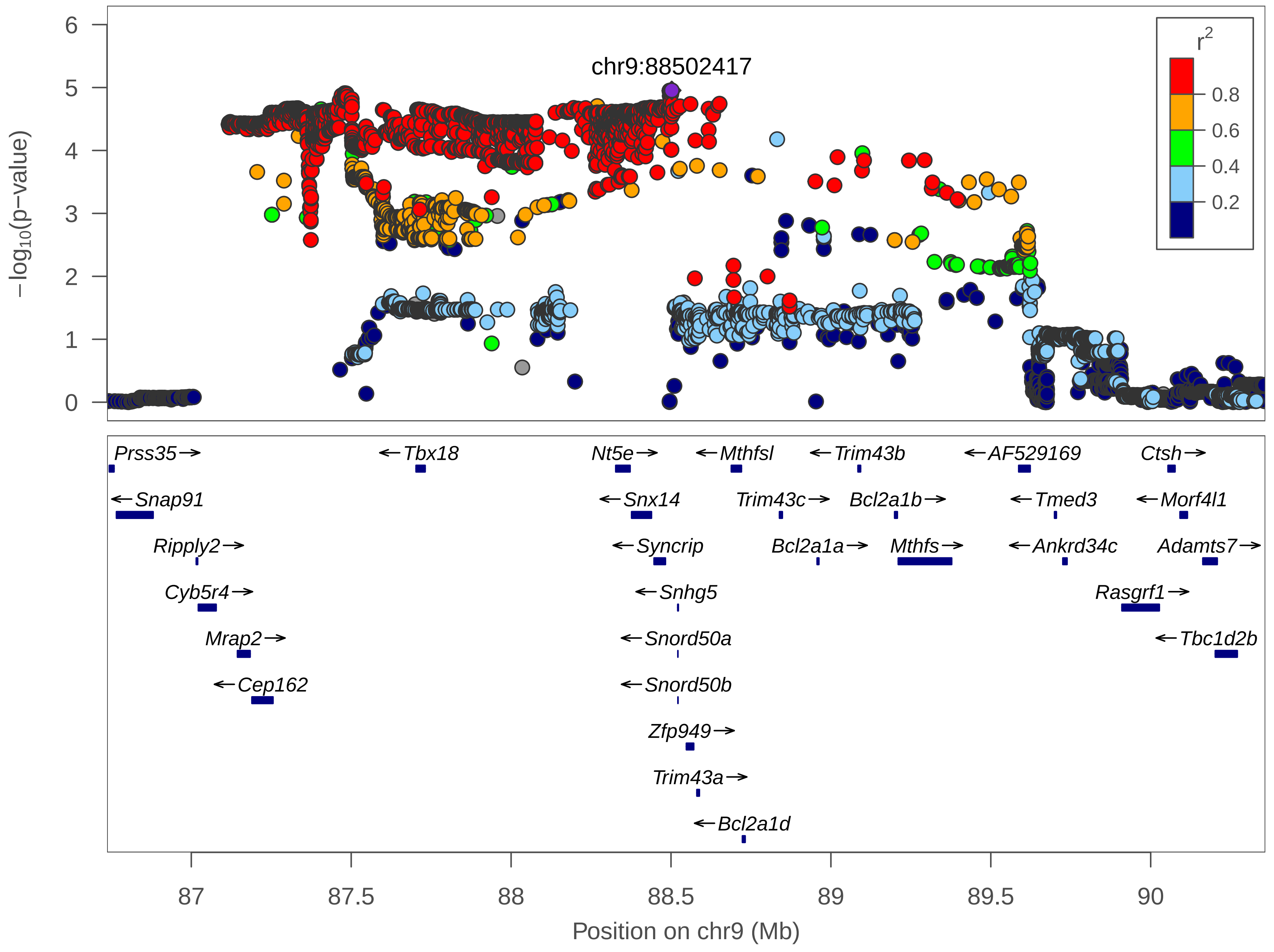

chr9:88502417
r2
0.8
0.6
0.4
0.2
−log10(p−value)
0
1
2
3
4
5
6
Prss35
Snap91
Ripply2
Cyb5r4
Mrap2
Cep162
Tbx18
Nt5e
Snx14
Syncrip
Snhg5
Snord50a
Snord50b
Zfp949
Trim43a
Bcl2a1d
Mthfsl
Trim43c
Bcl2a1a
Trim43b
Bcl2a1b
Mthfs
AF529169
Tmed3
Ankrd34c
Rasgrf1
Ctsh
Morf4l1
Adamts7
Tbc1d2b
87
87.5
88
88.5
89
89.5
90
Position on chr9 (Mb)

| | |
|---|---|
| date: | Fri Apr 2 15:56:19 2021 |
| build: | mice |
| display range: | chr9:86736657−90358438 [86736657−90358438] |
| hilite range: | 0 − 0 [ 0 − 0 ] |
| reference SNP: | chr9:88502417 |
| number of SNPs plotted: | 7620 |
| min P−value: | 1.11E−5 [chr9:88502417] |
| max P−value: | 10E−1 [chr9:89670316] |

Make more plots at http://csg.sph.umich.edu/locuszoom/
