## Supplementary material for "Dissecting indirect genetic effects from peers in laboratory mice": locusZoom plots for all significant IGE loci: PST.Immobility.Last4min_chr17_70527261_72790006_chr17_71304770.pdf

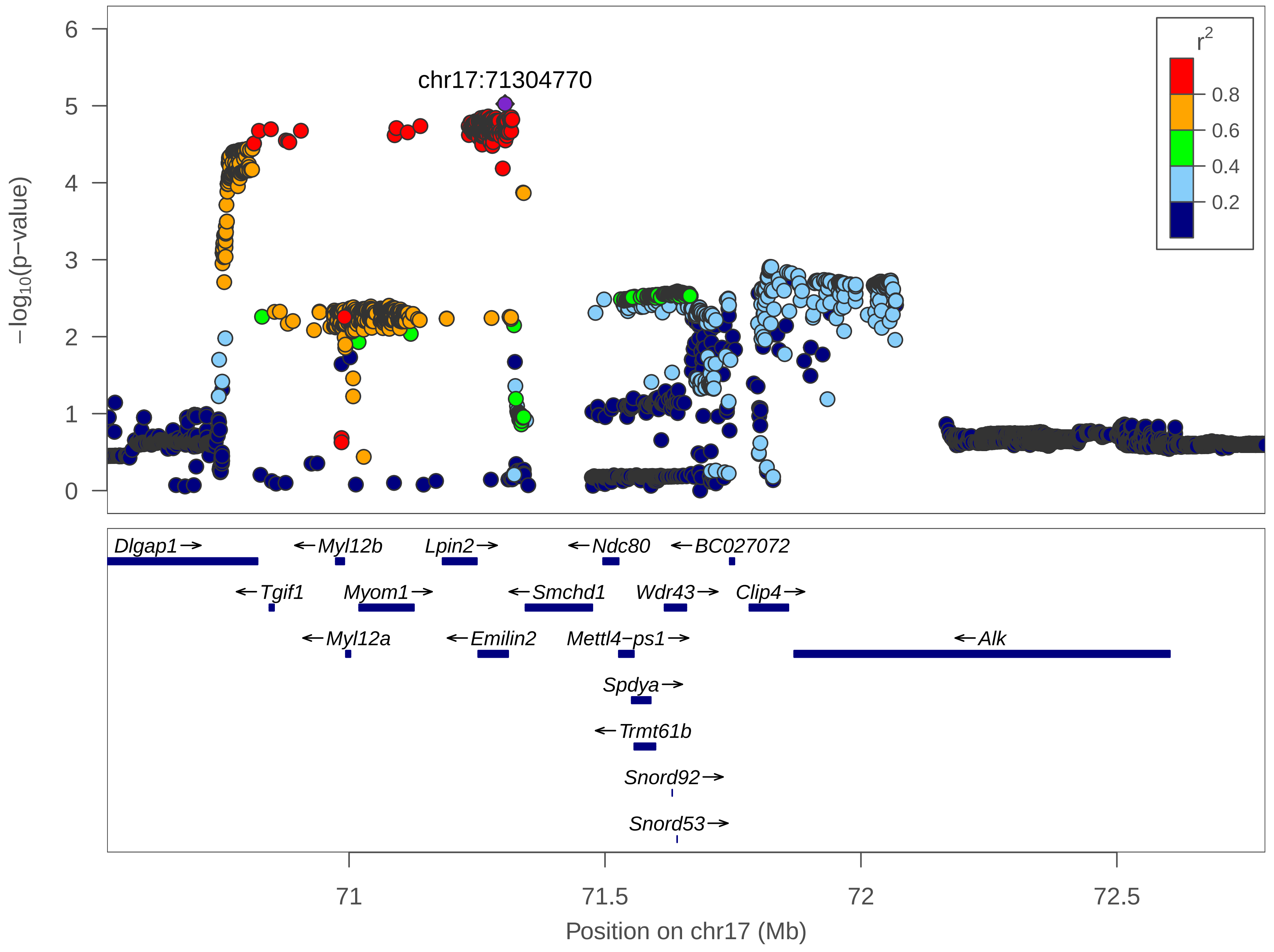

chr17:71304770
r2
0.8
0.6
0.4
0.2
−log10(p−value)
0
1
2
3
4
5
6
Dlgap1→
←Myl12b
Lpin2→
←Ndc80
←BC027072
←Tgif1
Myom1→
←Smchd1
Wdr43→
Clip4→
←Myl12a
←Emilin2
Mettl4−ps1→
←Alk
Spdya→
←Trmt61b
Snord92→
Snord53→
71
71.5
72
72.5
Position on chr17 (Mb)

| | |
|---|---|
| date: | Fri Apr 2 15:56:35 2021 |
| build: | mice |
| display range: | chr17:70527261−72790006 [70527261−72790006] |
| hilite range: | 0 − 0 [ 0 − 0 ] |
| reference SNP: | chr17:71304770 |
| number of SNPs plotted: | 6939 |
| min P−value: | 9.45E−6 [chr17:71304770] |
| max P−value: | 9.96E−1 [chr17:71685917] |

Make more plots at http://csg.sph.umich.edu/locuszoom/
