## Supplementary material for "Dissecting indirect genetic effects from peers in laboratory mice": locusZoom plots for all significant IGE loci: Sleep.s12h_D_BWcorr_chr11_31954132_36659978_chr11_35911554.pdf

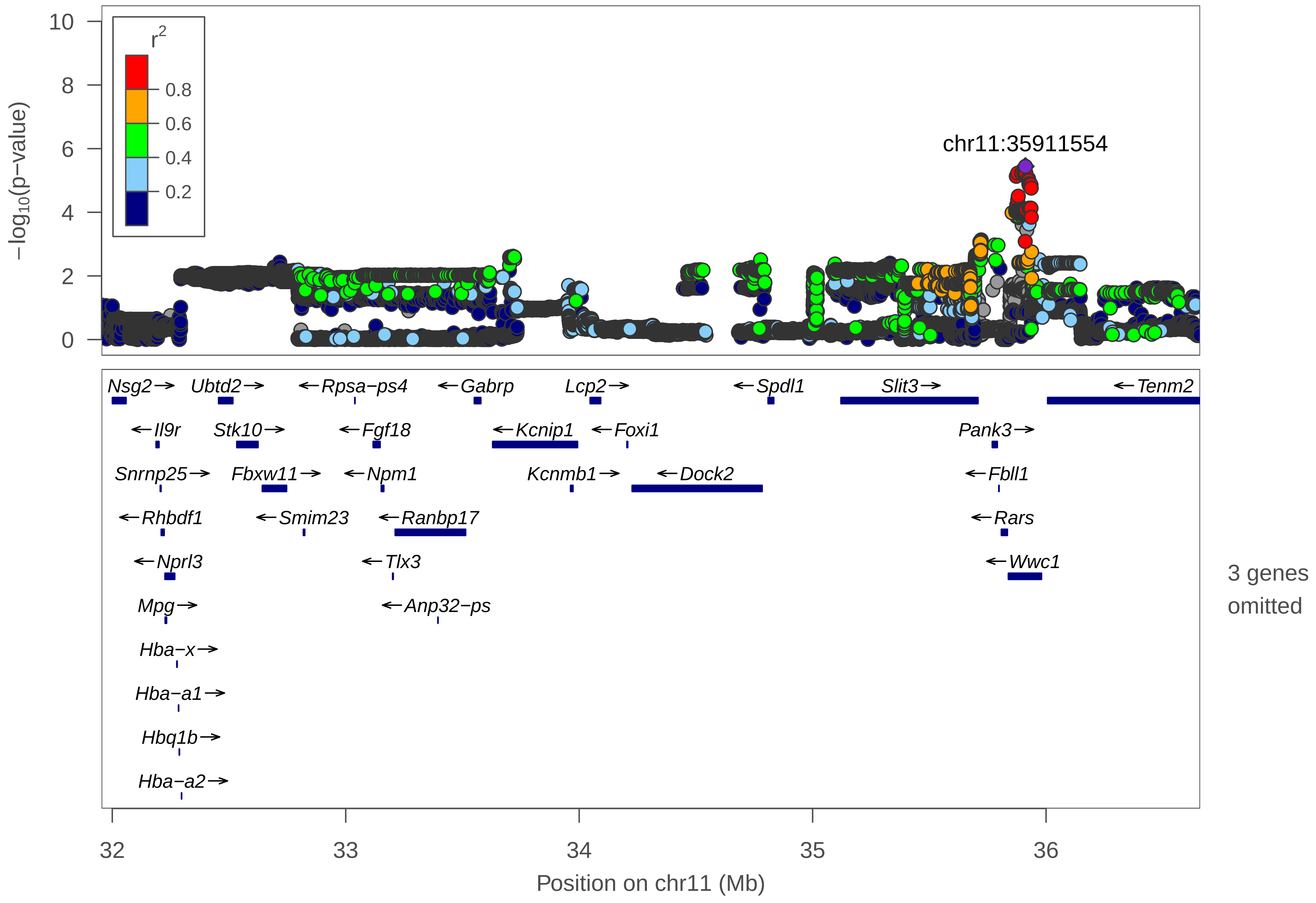

10
8
6
4
2
0
$-\log_{10}$(p−value)
$r^2$
0.8
0.6
0.4
0.2
chr11:35911554
Nsg2→
Ubtd2→
←Rpsa−ps4
←Gabrp
Lcp2→
←Spdl1
Slit3→
←Tenm2
←Il9r
Stk10→
←Fgf18
←Kcnip1
←Foxi1
Pank3→
Snrnp25→
Fbxw11→
←Npm1
Kcnmb1→
←Dock2
←Fbll1
←Rhbdf1
←Smim23
←Ranbp17
←Rars
←Nprl3
←Tlx3
←Wwc1
Mpg→
←Anp32−ps
Hba−x→
Hba−a1→
Hbq1b→
Hba−a2→
3 genes omitted
32
33
34
35
36
Position on chr11 (Mb)

| | |
|---|---|
| date: | Thu Apr 1 19:01:03 2021 |
| build: | mice |
| display range: | chr11:31954132−36659978 [31954132−36659978] |
| hilite range: | 0 − 0 [ 0 − 0 ] |
| reference SNP: | chr11:35911554 |
| number of SNPs plotted: | 42876 |
| min P−value: | 3.61E−6 [chr11:35911554] |
| max P−value: | 9.99E−1 [chr11:33611923] |
| omitted Genes: | Hbq1a, Sh3pxd2b, Efcab9 |

Make more plots at http://csg.sph.umich.edu/locuszoom/
