## Supplementary material for "Dissecting indirect genetic effects from peers in laboratory mice": locusZoom plots for all significant IGE loci: Sleep.s12h_D_BWcorr_chr13_104477636_107448960_chr13_106719493.pdf

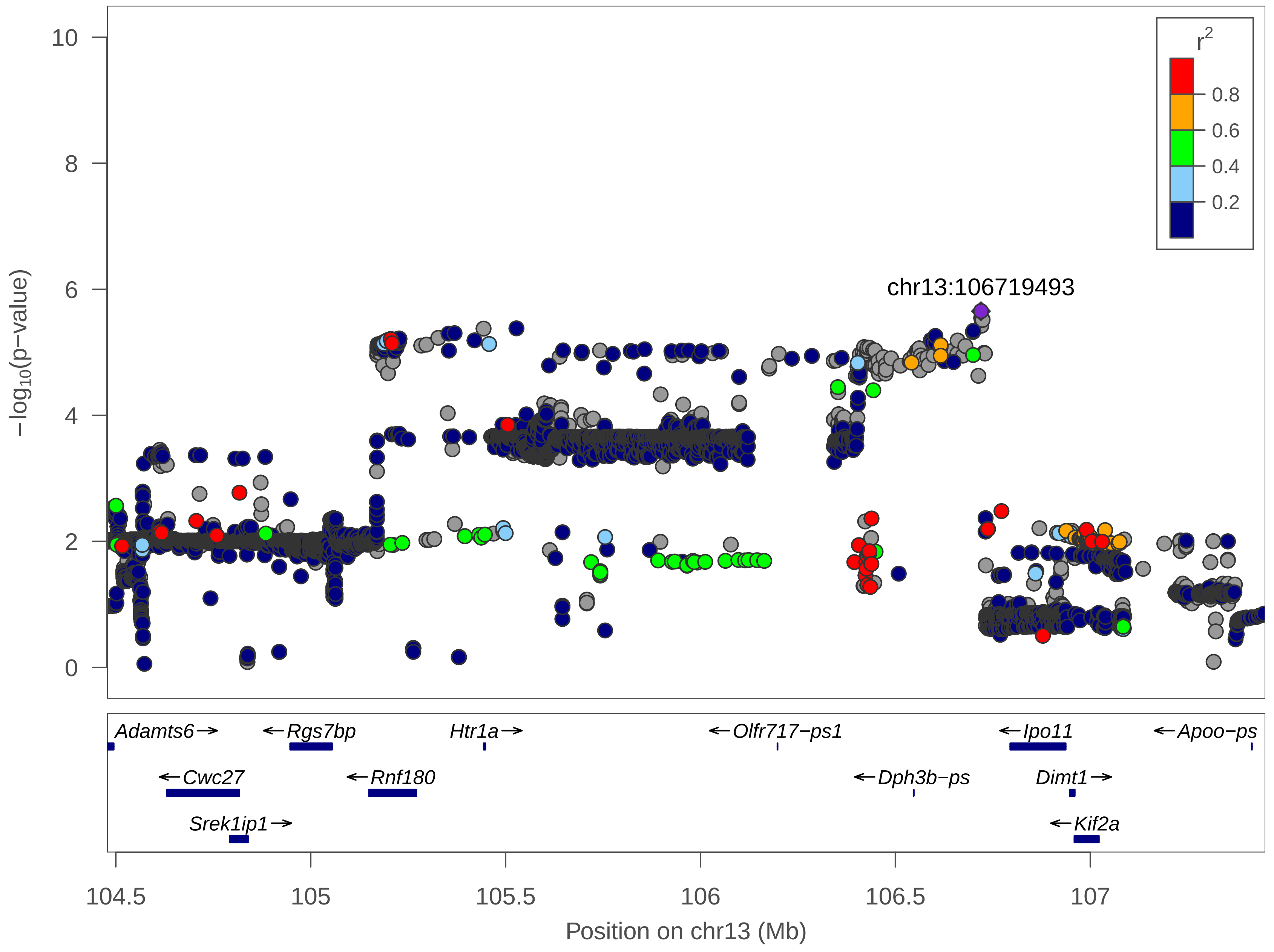

r2
0.8
0.6
0.4
0.2
chr13:106719493
−log10(p−value)
10
8
6
4
2
0
Adamts6
Rgs7bp
Htr1a
Olfr717−ps1
Ipo11
Apoo−ps
Cwc27
Rnf180
Dph3b−ps
Dimt1
Srek1ip1
Kif2a
104.5
105
105.5
106
106.5
107
Position on chr13 (Mb)

| | |
|---|---|
| date: | Thu Apr 1 19:00:26 2021 |
| build: | mice |
| display range: | chr13:104477636−107448960 [104477636−107448960] |
| hilite range: | 0 − 0 [ 0 − 0 ] |
| reference SNP: | chr13:106719493 |
| number of SNPs plotted: | 11137 |
| min P−value: | 2.22E−6 [chr13:106719493] |
| max P−value: | 8.76E−1 [chr13:104573375] |

Make more plots at http://csg.sph.umich.edu/locuszoom/
