## Supplementary material for "Dissecting indirect genetic effects from peers in laboratory mice": locusZoom plots for all significant IGE loci: Sleep.s12h_L_BWcorr_chr6_134458975_135958975_chr6_135204878.pdf

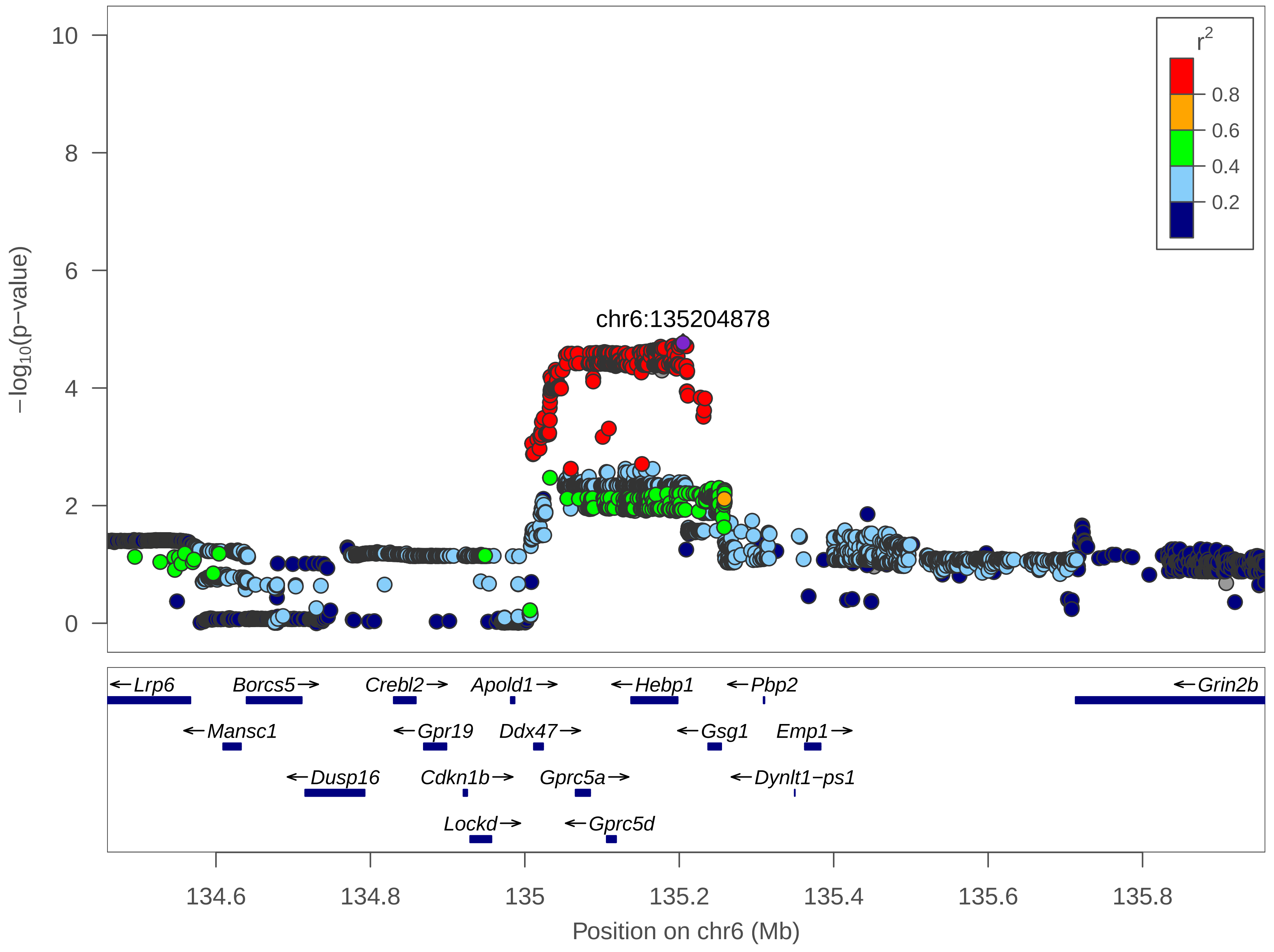

r²
0.8
0.6
0.4
0.2
chr6:135204878
−log10(p−value)
10
8
6
4
2
0
Lrp6
Borcs5
Crebl2
Apold1
Hebp1
Pbp2
Grin2b
Mansc1
Gpr19
Ddx47
Gsg1
Emp1
Dusp16
Cdkn1b
Gprc5a
Dynlt1−ps1
Lockd
Gprc5d
134.6
134.8
135
135.2
135.4
135.6
135.8
Position on chr6 (Mb)

| | |
|---|---|
| date: | Thu Apr 1 18:58:56 2021 |
| build: | mice |
| display range: | chr6:134458975−135958975 [134458975−135958975] |
| hilite range: | 0 − 0 [ 0 − 0 ] |
| reference SNP: | chr6:135204878 |
| number of SNPs plotted: | 5086 |
| min P−value: | 1.71E−5 [chr6:135204878] |
| max P−value: | 9.99E−1 [chr6:134730181] |

Make more plots at http://csg.sph.umich.edu/locuszoom/
