## Supplementary material for "Dissecting indirect genetic effects from peers in laboratory mice": locusZoom plots for all significant IGE loci: Sleep.s12h_L_BWcorr_chr11_34737784_36237784_chr11_35458419.pdf

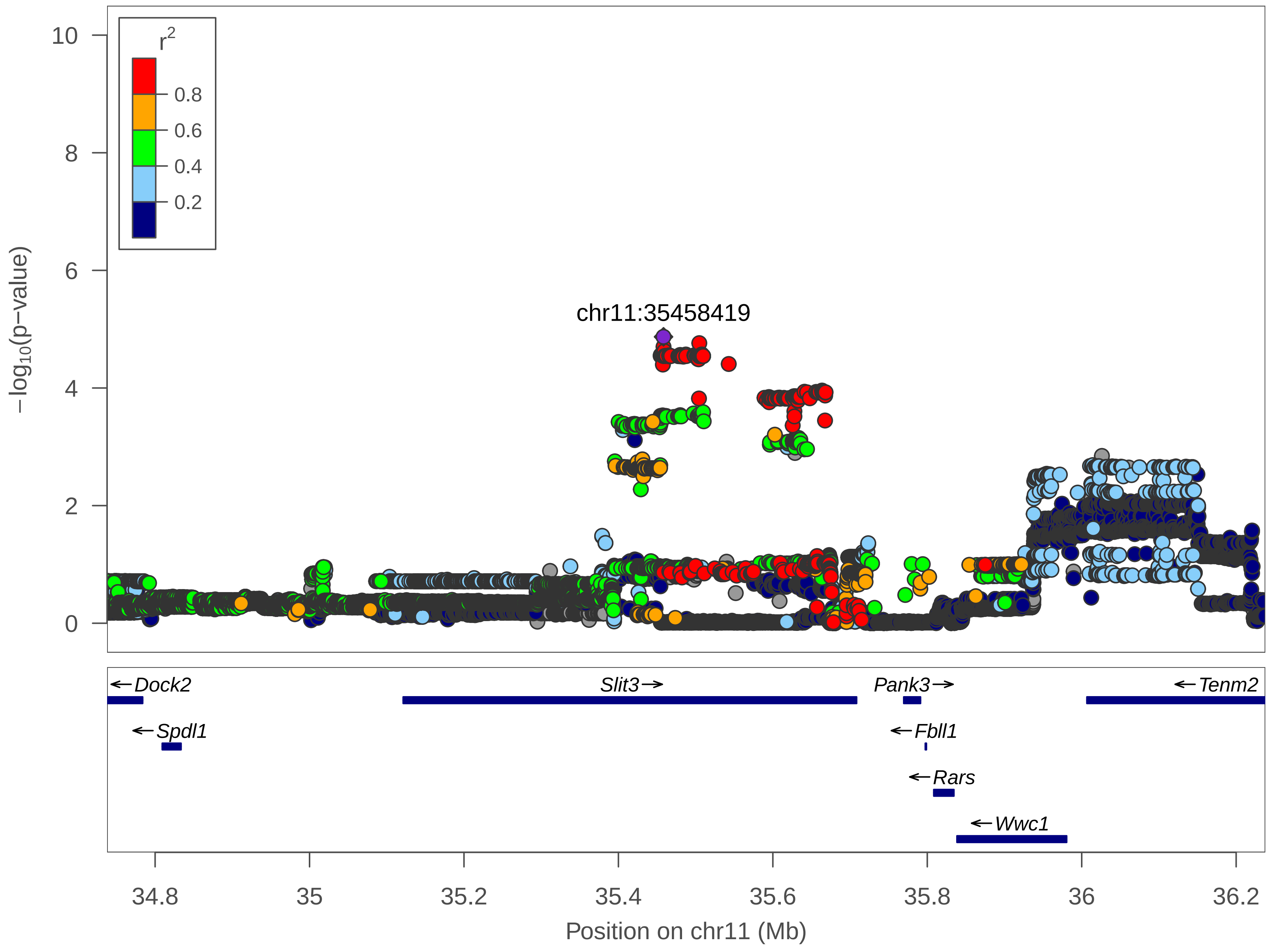

r2
0.8
0.6
0.4
0.2
10
8
6
4
2
0
−log10(p−value)
chr11:35458419
Dock2
Slit3
Pank3
Tenm2
Spdl1
Fbll1
Rars
Wwc1
34.8
35
35.2
35.4
35.6
35.8
36
36.2
Position on chr11 (Mb)

| | |
|---|---|
| date: | Thu Apr 1 18:59:56 2021 |
| build: | mice |
| display range: | chr11:34737784−36237784 [34737784−36237784] |
| hilite range: | 0 − 0 [ 0 − 0 ] |
| reference SNP: | chr11:35458419 |
| number of SNPs plotted: | 15878 |
| min P−value: | 1.35E−5 [chr11:35458419] |
| max P−value: | 10E−1 [chr11:35680122] |

Make more plots at http://csg.sph.umich.edu/locuszoom/
