## Supplementary material for "Dissecting indirect genetic effects from peers in laboratory mice": locusZoom plots for all significant IGE loci: Sleep.s12h_L_BWcorr_chr19_55003132_56503132_chr19_55282275.pdf

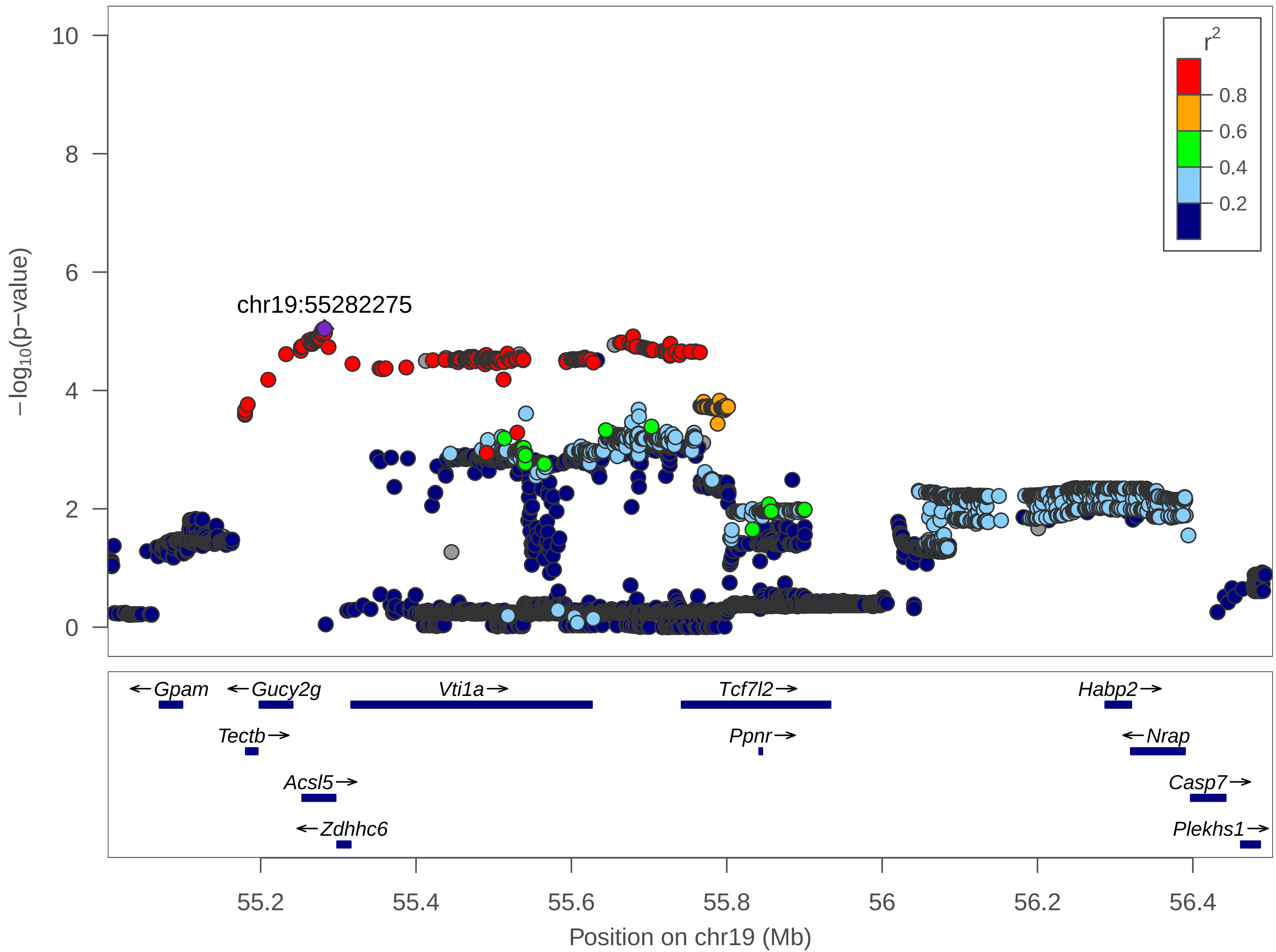

10
8
6
4
2
0
−log10(p−value)
chr19:55282275
r2
0.8
0.6
0.4
0.2
←Gpam
←Gucy2g
Vti1a→
Tcf7l2→
Habp2→
Tectb→
Ppnr→
←Nrap
Acsl5→
Casp7→
←Zdhhc6
Plekhs1→
55.2
55.4
55.6
55.8
56
56.2
56.4
Position on chr19 (Mb)

| | |
|---|---|
| date: | Thu Apr 1 18:59:24 2021 |
| build: | mice |
| display range: | chr19:55003132−56503132 [55003132−56503132] |
| hilite range: | 0 − 0 [ 0 − 0 ] |
| reference SNP: | chr19:55282275 |
| number of SNPs plotted: | 8674 |
| min P−value: | 9.08E−6 [chr19:55282275] |
| max P−value: | 9.99E−1 [chr19:55717146] |

Make more plots at http://csg.sph.umich.edu/locuszoom/
