## Supplementary material for "Dissecting indirect genetic effects from peers in laboratory mice": locusZoom plots for all significant IGE loci: Sleep.VAR_1h_BWcorr_chr3_142826479_144326479_chr3_143575069.pdf

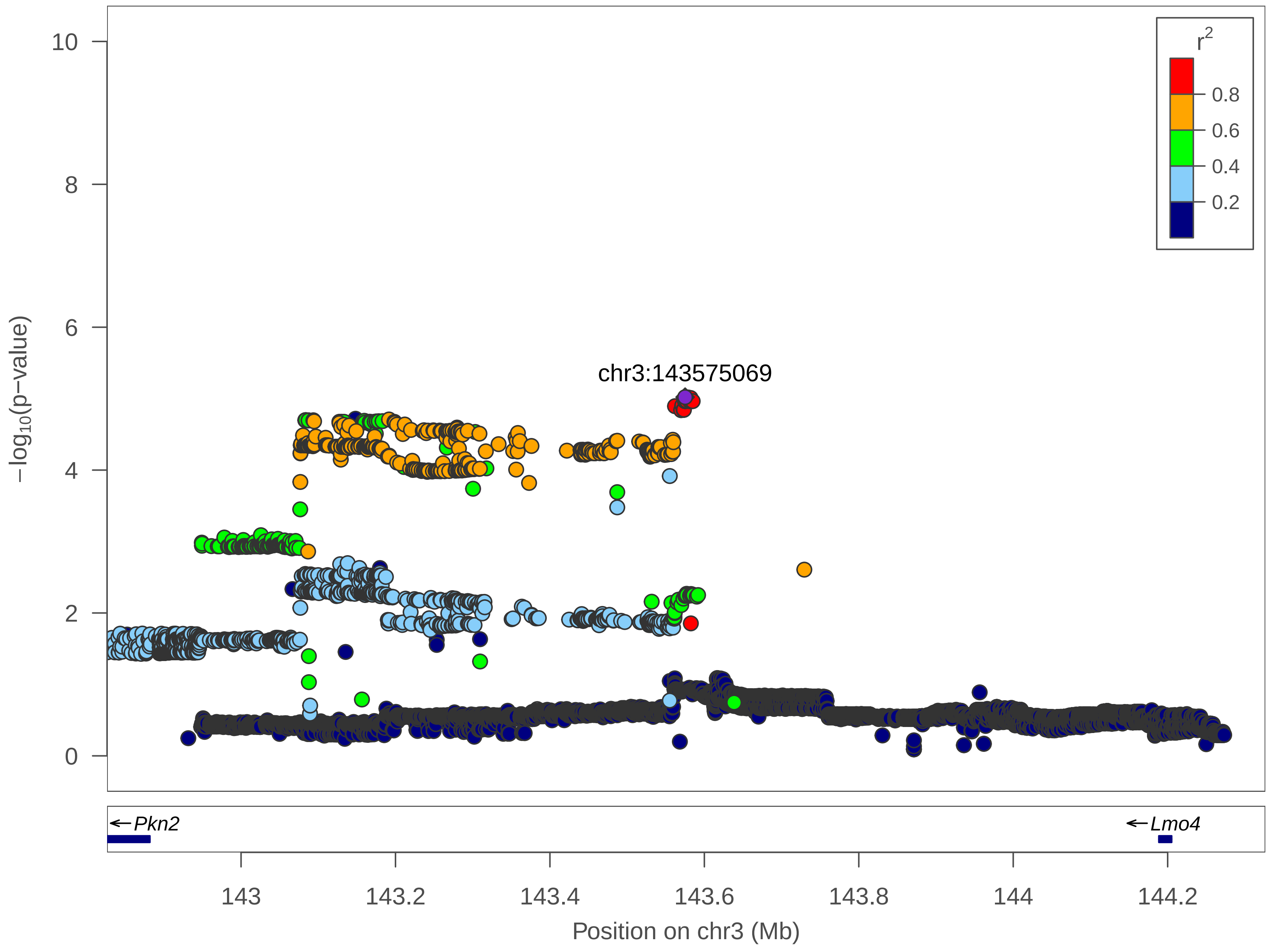

r²
0.8
0.6
0.4
0.2
10
8
6
4
2
0
−log10(p−value)
chr3:143575069
←Pkn2
←Lmo4
143
143.2
143.4
143.6
143.8
144
144.2
Position on chr3 (Mb)

| | |
|---|---|
| date: | Thu Apr 1 19:01:58 2021 |
| build: | mice |
| display range: | chr3:142826479−144326479 [142826479−144326479] |
| hilite range: | 0 − 0 [ 0 − 0 ] |
| reference SNP: | chr3:143575069 |
| number of SNPs plotted: | 11766 |
| min P−value: | 9.53E−6 [chr3:143575069] |
| max P−value: | 8.12E−1 [chr3:143871387] |

Make more plots at http://csg.sph.umich.edu/locuszoom/
