## Supplementary material for "Dissecting indirect genetic effects from peers in laboratory mice": locusZoom plots for all significant IGE loci: Sleep.VAR_1h_BWcorr_chr12_10091326_14517234_chr12_10840541.pdf

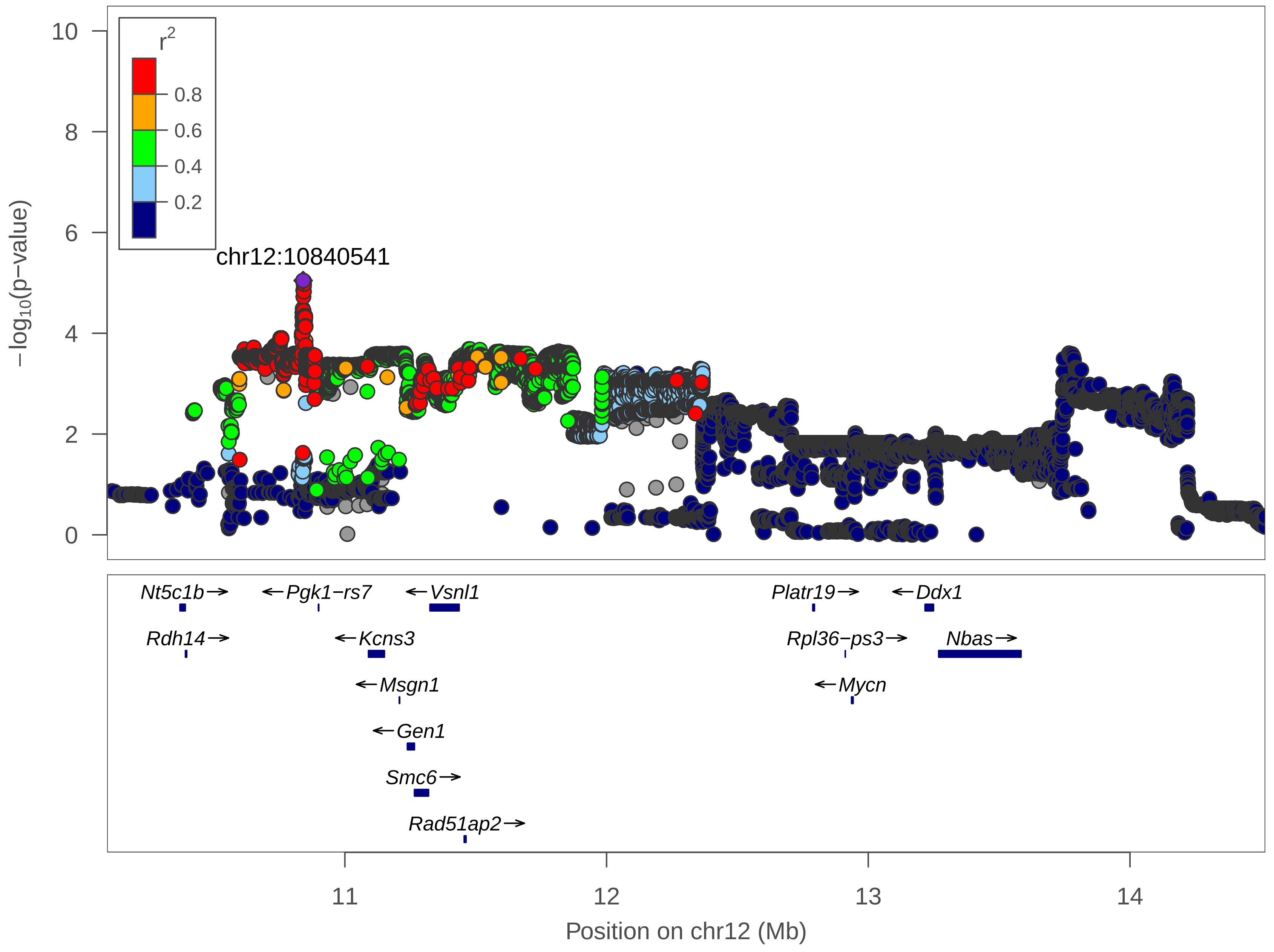

r2
0.8
0.6
0.4
0.2
chr12:10840541
10
8
6
4
2
0
−log10(p−value)
Nt5c1b→
←Pgk1−rs7
←Vsnl1
Platr19→
←Ddx1
Rdh14→
←Kcns3
Rpl36−ps3→
Nbas→
←Msgn1
←Mycn
←Gen1
Smc6→
Rad51ap2→
11
12
13
14
Position on chr12 (Mb)

| | |
|---|---|
| date: | Thu Apr 1 19:02:32 2021 |
| build: | mice |
| display range: | chr12:10091326−14517234 [10091326−14517234] |
| hilite range: | 0 − 0 [ 0 − 0 ] |
| reference SNP: | chr12:10840541 |
| number of SNPs plotted: | 33430 |
| min P−value: | 9E−6 [chr12:10840541] |
| max P−value: | 9.97E−1 [chr12:13165173] |

Make more plots at http://csg.sph.umich.edu/locuszoom/
